# An extremophilic *Nocardiopsis* strain from Great Salt Lake expands the taxonomic range of mycolic acid biosynthesis

**DOI:** 10.1101/2025.10.11.681818

**Authors:** Abigail F. Scott, Tianyun Jin, Elijah R. Bring Horvath, Nyssa K. Krull, Reiko Iizumi, Austin Bender, Brian T. Murphy, William Fenical, Jaclyn M. Winter

**Affiliations:** Department of Pharmacology and Toxicology, University of Utah, Salt Lake City, UT, USA; Center for Marine Biotechnology and Biomedicine, Scripps Institution of Oceanography, University of California at San Diego, San Diego, CA, USA; Department of Pharmaceutical Sciences, College of Pharmacy, University of Illinois at Chicago, Chicago, IL, USA; Center for Biomolecular Sciences, College of Pharmacy, University of Illinois at Chicago, Chicago, IL, USA; Skaggs School of Pharmacy and Pharmaceutical Sciences, University of California at San Diego, San Diego, CA, USA; Moores Comprehensive Cancer Center, University of California at San Diego, San Diego, California, United States

## Abstract

Mycolic acids, long-chain fatty acids that form the characteristic and relatively impermeable mycomembrane, have long been considered a defining chemotaxonomic feature of the order *Mycobacteriales*. Here, we report that *Nocardiopsis bonnevillensis,* a new type strain isolated from the hypersaline environment of Great Salt Lake, is the first organism outside of this order known to produce mycolic acids. Lipid profiling, acid-fast staining, isoniazid sensitivity, and genome mining confirmed hallmark features of mycolic acid biosynthesis. Evaluation of the strain’s metabolic capabilities led to the isolation of bonnevanoside, a thiophenyl nonulopyranoside reported here for the first time from a natural source. Moreover, additional Great Salt Lake-derived *Nocardiopsis* isolates exhibited acid-fast staining, suggesting that this trait may be more widespread than previously recognized. Altogether, these findings expand the taxonomic distribution of mycolic acid biosynthesis, challenge long-standing chemotaxonomic boundaries, and highlight the potential ecological significance of mycolate-containing envelopes in supporting bacterial survival in extreme environments.

## MAIN

Mycolic acids are very long-chain fatty acids that form a distinctive, relatively impermeable outer membrane known as the mycomembrane. This feature has long been considered unique to members of the order *Mycobacteriales*, where mycolic acid biosynthesis evolved through adaptations of primary fatty acid synthesis pathways. While most bacteria synthesize fatty acids *de novo* using the type II fatty acid synthase system (FAS-II), *Mycobacteriales* encode both FAS-I and FAS-II systems, with FAS-II elongating fatty acids into meromycolate chains. In contrast, members of *Corynebacteriaceae* produce mycolic acids in the apparent absence of a canonical FAS-II, relying instead on a multifunctional FAS-I to support both initiation and elongation steps of fatty acid biosynthesis^1,2^.

Mycolic acid production also requires specialized downstream enzymes, including PKS13, which catalyzes the condensation of the α-alkyl branch with the activated meromycolate chain to afford mature mycolic acids (Fig.1A). Disruption of *pks*13 abolishes mycolic acid biosynthesis in *Corynebacterium glutamicum*, emphasizing its essential biosynthetic role^3^. Given the complexity of the biosynthetic pathway and the lack of gene clustering, horizontal gene transfer of the complete mycolic acid biosynthetic pathway is unlikely. Thus, it is unsurprising that mycolic acid biosynthesis has, until now, been confined to members of the order *Mycobacteriales*. However, because the pathway evolved from conserved primary metabolism and exhibits lineage-specific adaptations, it remains plausible that mycolic acid biosynthesis could have arisen independently in other bacterial taxa through convergent evolution.

**Figure 1.**
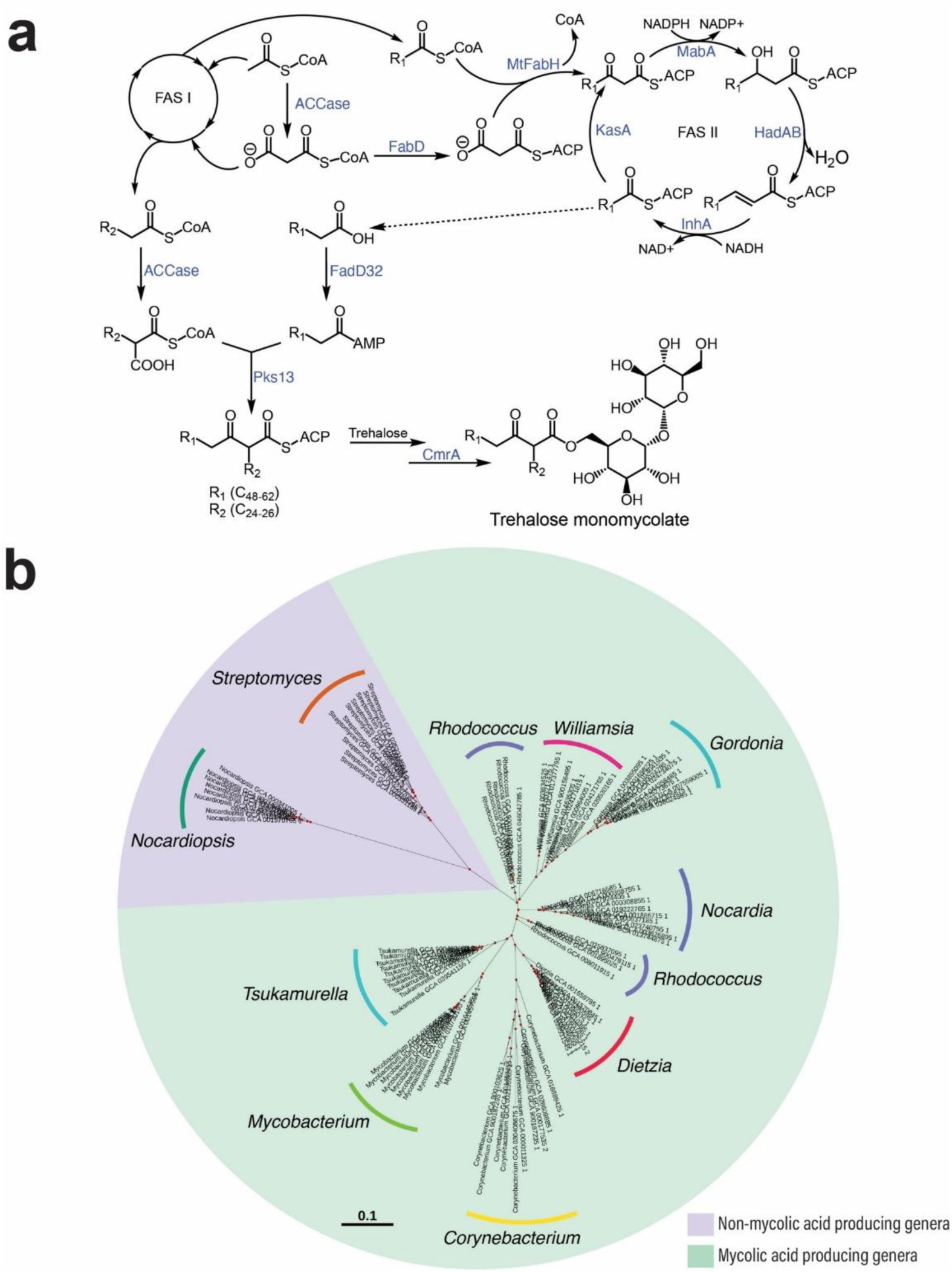
Mycolic acid biosynthesis and taxonomic distribution across Actinomycetota. **a**, Schematic of mycolic acid biosynthesis in *Mycobacterium* species, which utilize both FAS-I and FAS-II systems. Enzymes involved in meromycolate chain elongation and final condensation are shown, along with their associated cofactors and substrates. Adapted from Marrakchi *et al*.^2^. ACP, acyl carrier protein; CoA, Co-enzyme A; CmrA, mycolic acid reductase; KasA, meromycolic acid-3-oxoacyl-ACP synthase; FAD32, fatty-acyl-AMP ligase; ACCase, acyl-CoA carboxylase; MtFabH, beta-ketoacyl-ACP synthase; MabA, meromycolic acid 3-oxoacyl-ACP reductase; hadAB, meromycolic acid-3-hydroxy-acyl-ACP dehydratase; InhA, meromycolic acid enoyl-ACP reductase. **b**, Phylogenetic tree of representative genera within the *Actinomycetota* phylum, annotated with known mycolic acid producers (shaded in cyan) and their corresponding overall chain lengths. While members of the order *Mycobacteriales,* such as *Mycobacterium, Tsukamurella, Nocardia,* and *Rhodococcus,* are well-established mycolic acid producers, genera such as *Streptomyces* and *Nocardiopsis* (shaded in purple) are characterized by the absence of mycolic acids.

Following synthesis, mycolic acids are transported out of the cytoplasm and incorporated into the outer membrane as either free lipids or esterified to arabinogalactan^2^. This distinctive mycomembrane contributes to pathogen persistence, protects against desiccation, and provides intrinsic resistance to antibiotics and environmental stresses^4,5^. In pathogens such as *Mycobacterium tuberculosis*, these properties are critical for host survival, and the mycolic acid biosynthetic pathway has therefore become an important drug target. For example, the antitubercular drug isoniazid inhibits InhA, an enoyl reductase involved in meromycolate chain biosynthesis (Fig. 1A)^6,7^. However, the protective roles of mycolic acids are not limited to pathogenesis and may also reflect broader adaptations that support microbial survival in extreme environments.

*Actinomycetota* isolated from marine and saline environments are well known for their ability to produce biologically active and structurally diverse metabolites. The search for novel chemical scaffolds has increasingly turned toward microbial communities in extreme ecosystems, where selective pressures can drive both taxonomic diversification and chemical novelty. Our initial efforts to explore *Actinomycetota* from Great Salt Lake (GSL), an extreme hypersaline environment, uncovered a number of previously uncharacterized taxa^8^. Within this context, the genus *Nocardiopsis* stands out as particularly promising for natural product discovery. Members of this genus have been isolated from environments ranging from compost and indoor air to marine sediments and plant tissues, but they are most commonly associated with saline and alkaline ecosystems^9–16^. Their remarkable adaptability to such extreme conditions, the relatively small number of characterized species, and the growing interest in extremophilic Actinomycetota position *Nocardiopsis* as a valuable genus for expanding our understanding of microbial diversity and discovering novel bioactive metabolites.

GSL is a terminal, hypersaline lake in Northern Utah, USA and one of the most saline bodies of water on Earth^17,18^. A railroad causeway bisects the lake into two distinct regions, the North arm (Gunnison Bay) and South arm (Gilbert Bay), and restricts the mixing of water between the two arms. Most freshwater enters the South arm, where salinity typically ranges from 8‒17%, whereas salinity in the North arm commonly exceeds 27%^19^. In addition to salinity, GSL is situated at ∼1280 m above sea level and experiences dramatic seasonal water temperature fluctuations (∼0.5°C to ∼45°C)^18,20^. Microorganisms inhabiting GSL must therefore tolerate not only hyper salinity and temperature extremes but also occasional exposure to anthropogenic inputs such as treated municipal wastwater^20,21^. These selective pressures likely favor microbes with specialized cellular structures, including modified cell envelopes, and metabolic strategies that promote environmental resilience.

Here, we describe one such polyextremophile, *N. bonnevillensis*, a new type strain isolated from GSL sediment. This strain represents the first reported example of a mycolic acid-producing organism outside the order *Mycobacteriales* (Fig. 1B). Multiple lines of evidence, such as mass spectrometric lipid analyses, acid-fast staining, isoniazid sensitivity, and genome mining that revealed homologs of all essential mycolic acid biosynthesis genes, confirm this unexpected capacity. Given the substantial metabolic investment and genetic complexity associated with mycolic acid production, we next investigated whether these lipids confer a functional advantage under GSL’s extreme environmental conditions. Extending this analysis to additional *Nocardiopsis* isolates from GSL revealed several more acid-fast positive strains, suggesting that mycolic acid production may be more widespread within this genus than previously recognized. Their broad phylogenetic distribution raises important questions about the taxonomic range, ecological significance, and evolutionary origins of this adaptation.

## RESULTS

### Isolation and identification of a novel type strain

*Nocardiopsis bonnevillensis* was isolated from sediment collected in July 2021 along the shoreline of Farmington Bay, Great Salt Lake (41° 3′ 58″ N, 112° 13′ 48″ W), within Antelope Island State Park, Utah. Farmington Bay, located in the southeastern portion of the lake, receives ∼50% of its water input from municipal treatment plants and is connected to the mainland by the Antelope Island Causeway^22^. The Farmington Bay sediment sample displayed an electrical conductivity of 40.6 milli-Siemens (mS), compared to 5.90 mS for a marine sediment sample collected from a near-shore site adjacent to the Scripps Institution of Oceanography pier in La Jolla, California (32° 52′ 00″ N, 117° 15′ 26″ W), which was processed in the same manner.

### Whole-genome sequencing and phylogenomic characterization

To obtain a whole-genome sequence of *N. bonnevillensis*, genomic DNA was extracted using an MP Biomedicals FastDNA SPIN Kit according to the manufacturer’s instructions and sequenced by Plasmidsaurus using Oxford Nanopore. CheckM analysis of the assembled genome indicated 100.0% completeness and 0.5% contamination^23^. The 6.1 Mb genome was assembled into six contigs with an average coverage depth of 29x. To determine the taxonomic placement of *N. bonnevillensis*, we performed both whole-genome and 16S ribosomal RNA (rRNA) phylogenomic analyses. Average nucleotide identity (ANI) calculated using autoMLST’s curated reference database^24^, identified *N. ganjiahuensis* DSM_45031 as the closest match at 89.9% ANI, which is well below the 95‒96% species delineation threshold. Digital DNA-DNA hybridization (dDDH) analysis using the Type Strain Genome Server (TYGS)^25^ further supported this placement. The most similar genomes were *N. terrae* strains KCTC 19431 and DSM 45157 (NCBI accession numbers GCF_014651695 and GCF_014874035, respectively) (Fig. 2A), with a calculated dDDH value of 64.4% (formula d4; range: 64.4–70.2%), also below the 70% species cutoff.

**Figure 2:**
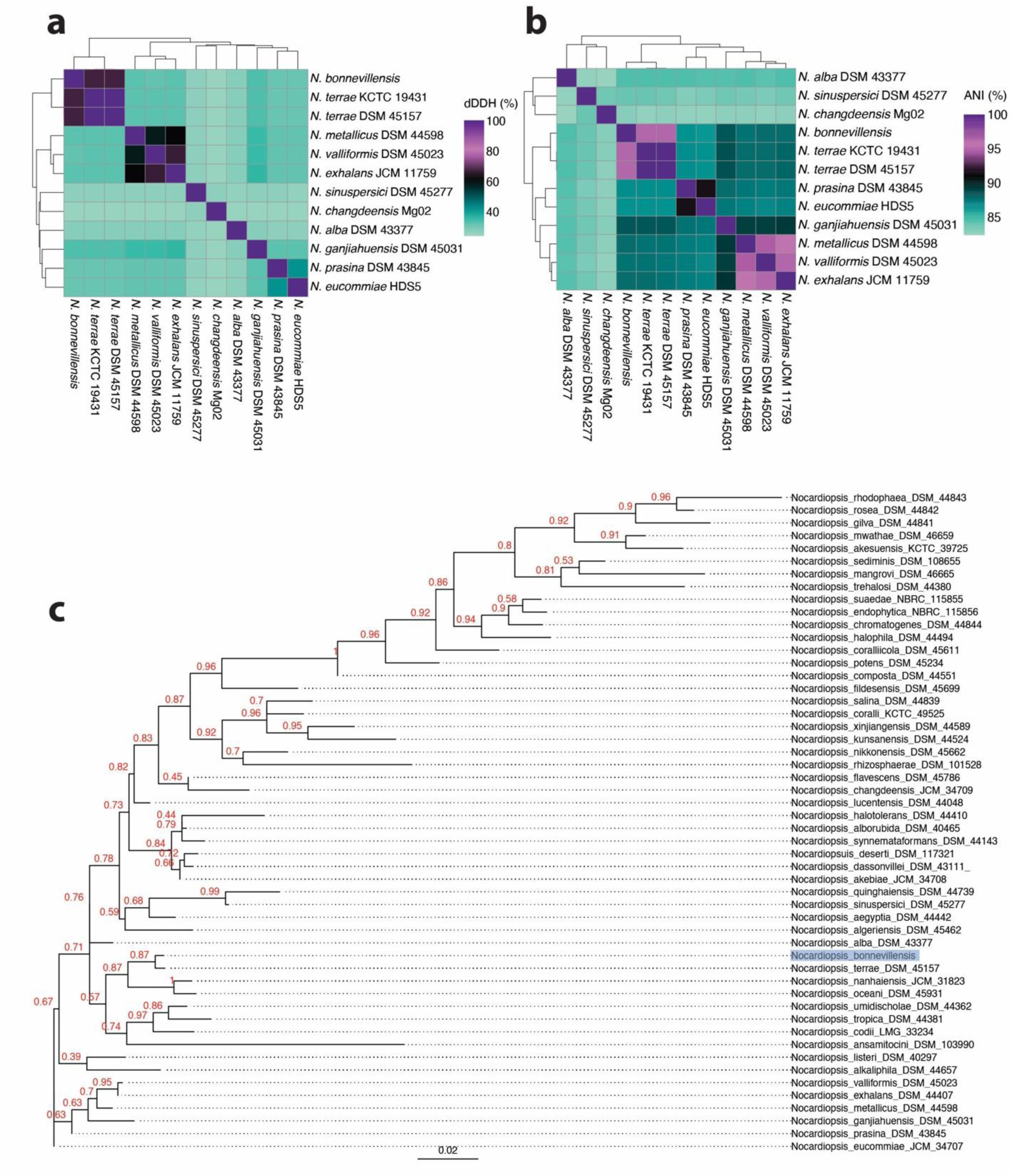
16S and whole genome comparisons of *N. bonnevillensis* to *Nocardiopsis* species. **a**, Heatmap of %dDDH of select *Nocardiopsis* spp. and *N. bonnevillensis.* **b**, Heat map of %ANI of select *Nocardiopsis* spp. and *N. bonnevillensis*; **c**, Maximum likelihood tree of 16S rRNA sequences from the 51 validly published and correctly named *Nocardiopsis* spp. and *N. bonnevillensis* created with RAxML^48^. *N. bonnevillensis* is highlighted in blue.

Because the *N. terrae* genomes were absent from the autoMLST database, we expanded our analysis to include 133 publicly available *Nocardiopsis* genomes from NCBI Datasets^26^. Pairwise ANI calculations confirmed the dDDH results: *N. bonnevillensis* shares 96.1% ANI with both *N. terrae* KCTC 19431 and DSM 45157 (Fig. 2B), narrowly exceeding the conventional ANI threshold for species-level distinction. To refine its placement, we restricted our phylogenomic dataset to 110 *Nocardiopsis* genomes exhibiting ≥ 82% ANI to *N. bonnevillensis*. Extracted 16S rRNA sequences were aligned and used to construct a maximum likelihood phylogenetic tree. Consistent with dDDH and ANI analyses, *N. bonnevillensis* clustered with *N. terrae* KCTC 19431 and DSM 45157 (Supplementary Fig. 1). To further enhance its taxonomic position, we conducted a whole genome phylogenomic analysis using PhyloPhlAn^27^, which leverages 360 universal marker genes. This maximum-likelihood tree placed *N. bonnevillensis* within the *N. terrae* clade, but as a distinct lineage near the species-level boundary (Extended Data Fig. 1). To understand *N. bonnevillensis’s* position among the 51 validly published and correctly named *Nocardiopsis* type strains, a maximum likelihood phylogenetic tree was constructed using their 16S rRNA sequences (Fig. 2C).

### Chemotaxonomic characterization supports novel species description

Based on the predicted taxonomic novelty, *N. bonnevillensis* was selected for in-depth chemotaxonomic analysis. These analyses were performed by DSMZ (Deutsche Sammlung von Mikroorganismen und Zellkulturen, Braunschweig, Germany) and included profiling of cellular fatty acids, respiratory quinones, polar lipids, whole-cell sugars, polyamines, diaminopimelic acid (Dpm) isomers, peptidoglycan structural elucidation (Extended Data Table 1), mycolic acids (Extended Data Table 2), matrix-assisted laser desorption/ionization (MALDI) Biotyper classification, Active Pharmaceutical Ingredient (API) Zym for enzymatic profiling (Extended Data Table 3), and API 50 CHB carbon utilization (Extended Data Table 4). MALDI Biotyper classification yielded genus-but not species-level resolution. On a scale of 0.00–3.00, the highest score was 1.87, consistent with a “probable genus identification” as *Nocardiopsis* sp., while the second-best score of 1.49 fell within the “not reliable identification” range. Together, these results support the designation of *N. bonnevillensis* as a novel species within the genus *Nocardiopsis*.

### Evaluation of biosynthetic potential and novel compound identification

To explore the metabolic capabilities of *N. bonnevillensis,* a crude extract from large-scale fermentation was screened for antimicrobial activity against a panel of Gram-positive and Gram-negative pathogens, including methicillin-resistant *Staphylococcus aureus* (MRSA) TCH1516, *Escherichia coli* LptD4213, *Pseudominas aeruginosa* PA01, *Bacillus oceanisediminis* CNY997, *Serratia marcescens* SNA111, and *Vibrio harveyi* SNA117. The extract exhibited moderate to strong activity against MRSA, *E. coli*, and *B. oceanisediminis*, but showed no detectable inhibition of *S. marcescens* or *V. harveyi* (Supplemental Result 4 and Supplementary Table 3).

To connect this activity with biosynthetic machinery, we mined the *N. bonnevillensis* genome using antiSMASH^28^, which revealed 19 putative biosynthetic gene clusters (BGCs) (Extended Data Fig. 2A, Supplementary Table 2). All 19 BGCs showed ≤ 85% similarity to characterized clusters in the MiBIG v.3 database^29^, classifying them as “uncharacterized” and suggesting that *N. bonnevillensis* harbors substantial untapped biosynthetic potential.

Guided by these findings, we used bioactivity-guided fractionation and purified a thiophenyl nonulopyranoside, bonnevanoside (**1**) (Extended Data Fig. 2B, Supplementary Tables 4–6 and Supplementary Fig. 4–13). Although bonnevanoside displayed no measurable activity against MRSA or *E. coli* at concentrations up to 100 µg/mL, this compound has been previously synthesized and is reported here for the first time from a natural source. The isolation of **1** is evidence of the biosynthetic novelty of *N. bonnevillensis* and highlights the potential of extremophilic *Nocardiopsis* species to produce structurally unique metabolites that expand the known chemical space of this genus. Together with its unusual lipid profile, these findings suggest that the chemical strategies employed by *N. bonnevillensis* may play an important role in adapting to the selective pressures of the Great Salt Lake environment.

### Identification of mycolic acids and the required biosynthetic genes

The most striking result from the chemotaxonomic analysis of *N. bonnevillensis* was the detection of mycolic acids, confirmed via electrospray ionization quadrupole time-of-flight mass spectrometry (ESI-QTOF-MS). Mass spectra matched the exact masses of known mycolic acid standards, and the lipid profile included α-mycolic acids C_79_H_154_O_3_ (3.14%) and C_81_H_158_O_3_ (10.19%), as well as dicarboxy- or dihydroxy-mycolic acids C_61_H_118_O_5_ (37.76%), C_62_H_120_O_5_ (25.68%), and C_64_H_124_O_5_ (23.23%) (Extended Data Table 2). Members of the genus *Nocardiopsis* are characterized by the absence of mycolic acids^30^, making this, to the best of our knowledge, the first report of mycolic acid production outside the order *Mycobacteriales*.

Acid-fast staining further supported these findings. Although the standard Ziehl–Neelsen protocol ^31^ yielded inconclusive results, *N. bonnevillensis* retained a pink hue between the destain and counterstain steps. Because species outside the family *Mycobacteriaceae* are often weakly acid-fast or stain negative with acid-alcohol destains, we applied a modified Ziehl-Neelsen method using an alcohol-free 1% sulfuric acid destain^32,33^. This approach produced a clear acid-fast positive result (Fig. 3a).

**Figure 3:**
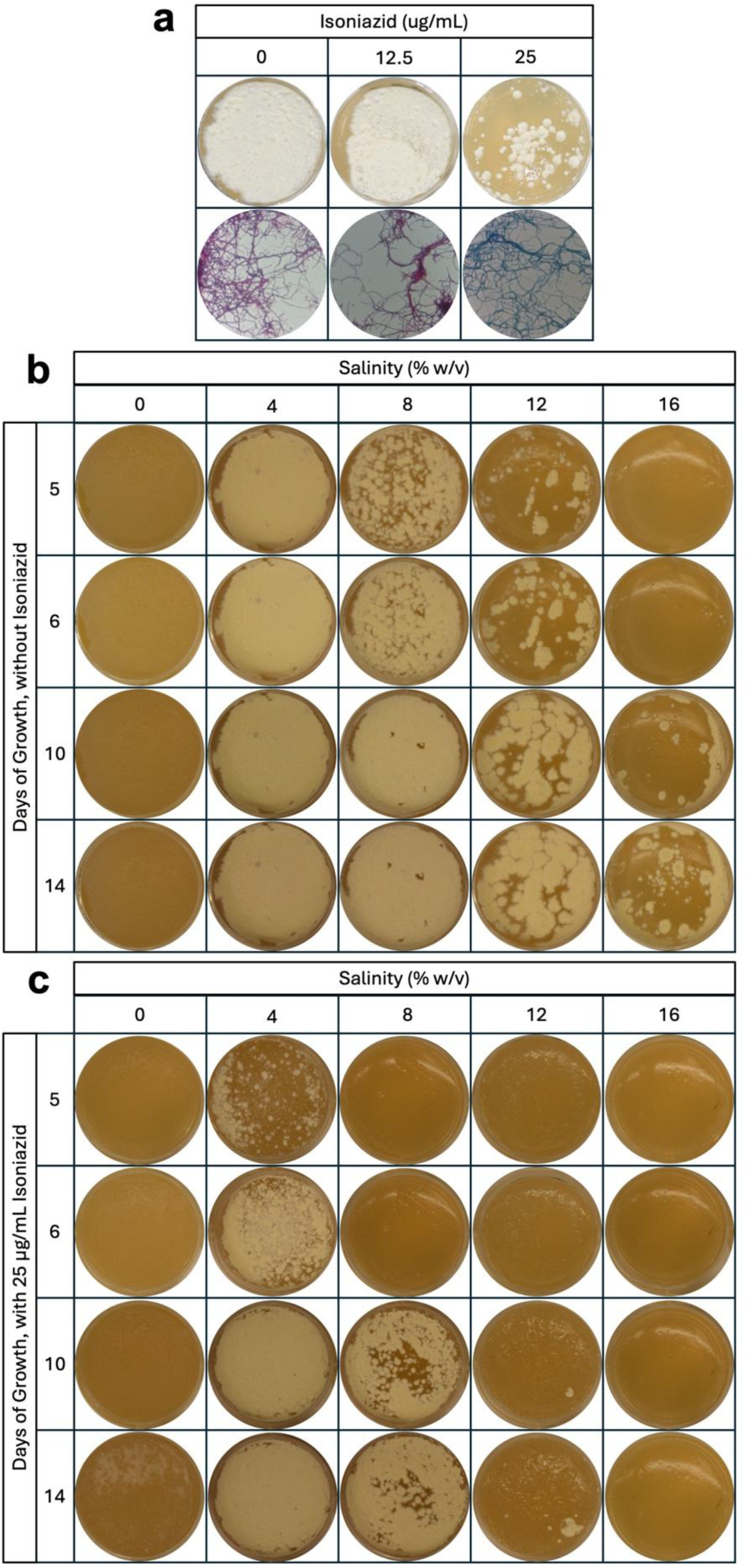
Effects of isoniazid and salinity on the growth of *N. bonnevillensis* **a**, Growth of *N. bonnevillensis* on YPM agar (8% salinity) after 6 days of incubation in the presence of 0, 12.5, or 25 µg/mL isoniazid (top), with corresponding acid-fast staining results shown below. **b**, Growth of *N. bonnevillensis* on YPM agar supplemented with increasing salinity concentrations (0–16% w/v) over a 14-day incubation period. **c**, Growth of *N. bonnevillensis* on YPM agar containing 25 µg/mL isoniazid under the same salinity conditions (0–16% w/v) over a 14-day incubation period.

Functional assays demonstrated that mycolic acid biosynthesis is essential for *N. bonnevillensis* growth under selective conditions. Cultures grown on YPM agar (8% salinity) supplemented with increasing concentrations of isoniazid (INH), an inhibitor of the InhA meromycolic acid enoyl-ACP reductase (Fig. 1a), exhibited minimal growth at ≥25 µg/mL after five days of incubation at 30 °C (Fig. 3a, Extended Data Figure 3). Acid-fast staining remained positive at 0 and 12.5 µg/mL of INH but was negative at 25 µg/mL, indicating that mycolic acid biosynthesis was inhibited under these conditions.

Environmental stress assays linked mycolic acid production to salinity tolerance. In the absence of INH, *N. bonnevillensis* grew across a broad salinity range (0–16%) (Fig. 3b, Supplementary Fig.14). In contrast, the presence of 25 µg/mL INH markedly reduced salinity tolerance, most notably at ≥ 8% salinity (Fig. 3c, Supplementary Fig.14). Growth and sporulation were comparable with or without INH on low-salt media and at 4% salinity but were severely reduced at 12% and 16% after 14 days of incubation. These results support the conclusion that mycolic acids contribute to salinity tolerance in *N. bonnevillensis*.

Comparative genomic analyses confirmed that the genes required for mycolic acid biosynthesis are present in the *N. bonnevillensis* genome. A curated set of protein sequences from 15 representative mycolate-producing species (Supplementary Table 7) was retrieved from the Kyoto Encyclopedia of Genes and Genomes (KEGG) ortholog tables for pathway modules M00886 (meromycolic acid biosynthesis) and M00887 (mycolic acid biosynthesis) and queried against the *N. bonnevillensis* genome using SeqForge (v0.0.1)^34^. Functional annotation with eggnog-mapper^35^ identified homologs of all essential biosynthetic genes (Table 1). The predicted Pks13 homolog was annotated using InterPro (European Bioinformatics Institute, EBI)^36^ and was found to contain an acyl transferase, ketosynthase, acyl carrier protein, and thioestersase domain with the expected organization consistent with canonical Pks13 architecture.^3^ Orthologs of *kas*B, *had*C, and *had*D were not detected in *N. bonnevillensis*, but these genes are restricted to the *Mycobacteriaceae* family (Fig. 1b) and are functionally equivalent to *kas*A, *had*A and *had*B. Interestingly, a FAS-I homolog was also not detected in the *N. bonnevillensis* genome.

**Table 1:** Homologs of known mycolic acid biosynthesis genes identified in N. bonnevillensis. S. piniformis (Skermania pinensis DSM 43998), M. smegmatis (Mycobacterium smegmatis MC2 155), D. timorensis (Dietzia timorensis ID05-A0528), T. fengzijianii (Tomitella fengzijianii HY188), W. herbipolensis (Williamsia herbipolensis NBC 00319), M. koreense (Mycobacterium koreense JCM 19956), R. equi (Rhodococcus equi 103S), R. jostii (Rhodococcus jostii RHA1), M. sinensis (Mycobacterium sinense JDM601), C. glutamicum (Corynebacterium glutamicum ATCC 13032).

| KEGG Ortholog | Gene Product | Protein function | Sequence Similarity (origin) | Match Accession Number | Similarity/ Identity (%) | Query Cover |
| --- | --- | --- | --- | --- | --- | --- |
| K02078 | AcpP | acyl carrier protein | *S. pinensis* | CP079105 | 86/70 | 100 |
| K11263 | BccA | $\alpha$ subunit of ACC | *M. smegmatis* | CP000480 | 77/65 | 100 |
| K18472 | AccD6 | $\beta$ subunit of ACC | *D. timorensis* | ANI92066 | 59/43 | 90 |
| K00645 | FabD | [ACP]- S-malonyltransferase | *T. fengzijianii* | CP041765 | 67/55 | 92 |
| K11608 | MtfabH | β-ketoacyl-ACP synthase III | *R. equi* | CBH50061 | 64/51 | 97 |
| K11609 | KasA | meromycolic acid 3-oxoacyl-ACP synthase I | *T. fengzijianii* | QDQ97938 | 67/50 | 99 |
| K11610 | MabA | meromycolic acid 3-oxoacyl-ACP reductase | *W. herbipolensis* | WUM20658 | 76/66 | 100 |
| K27071 | HadA | meromycolic acid (3R)-3hydroxyacyl-ACP dehydratase | *R. equi* | CBH49676 | 54/34 | 85 |
| K27072 | HadB | meromycolic acid (3R)-3hydroxyacyl-ACP dehydratase | *M. koreense* | BBY55061 | 70/50 | 99 |
| K11611 | InhA | meromycolic acid enoyl-ACP reductase | *R. jostii* | ABG98978 | 71/54 | 98 |
| K27070 | KasB | meromycolic acid 3-oxoacyl-ACP synthase II | - | - | - | - |
| K27073 | HadC | meromycolic acid (3R)-3hydroxyacyl-ACP dehydratase | - | - | - | - |
| K27074 | HadD | meromycolic acid (3R)-3hydroxyacyl-ACP dehydratase | - | - | - | - |
| K27093 | AccD4 | $\beta$ subunit of ACC | *R. equi* | CBH46355 | 70/53 | 97 |
| K27094 | AccD5 | $\beta$ subunit of ACC | *R. jostii* | ABG99963 | 80/74 | 100 |
| K27095 | AccE5 | $\varepsilon$ subunit of ACC | *M. sinense* | AEF37040 | 56/43 | 82 |
| K12428 | FadD32 | fatty acyl-AMP ligase | *M. koreense* | BBY56042 | 38/27 | 77 |
| K12437 | Pks13 | polyketide synthase | *C. glutamicum* | BAC00265 | 44/30 | 45 |
| K27306 | CmrA | mycolic acid reductase | *M. smegmatis* | ABK71182 | 50/38 | 94 |
| K11533 | FAS-I | fatty acid synthase | - | - | - | - |

### Identification of additional acid-fast positive Great Salt Lake-derived *Nocardiopsis* spp

From various GSL sediment samples, 13 additional bacterial isolates (Fig. 4a) were identified as *Nocardiopsis* spp. by 16S rRNA sequencing (Supplementary Information). The maximum likelihood tree of 16S rRNA sequences from the GSL *Nocardiopsis* strains and 51 *Nocardiopsis* type strains (Fig. 4b) showed some clustering of the GSL strains together, but overall the GSL strains were distributed among the type strain sequences. Because 16S rRNA sequence similarity is typically very high among *Nocardiopsis* strains and can overestimate relatedness^30^, we also compared MALDI-TOF protein mass spectra (3–20 kDa) obtained by analysis of dendrograms in the IDBac bioinformatics platform^37^. The resulting dendrogram largely recapitulated the relationships suggested by the 16S rRNA maximum likelihood tree, but also revealed clear distinctions among the GSL isolates (Extended Data Fig. 4). Notably, despite several isolates sharing ≥99–100% 16S rRNA sequence similarity, the protein dendrogram indicated that all 14 GSL *Nocardiopsis* isolates, including *N. bonnivellensis*, represent unique strains (Extended Data Fig. 4).

**Figure 4:**
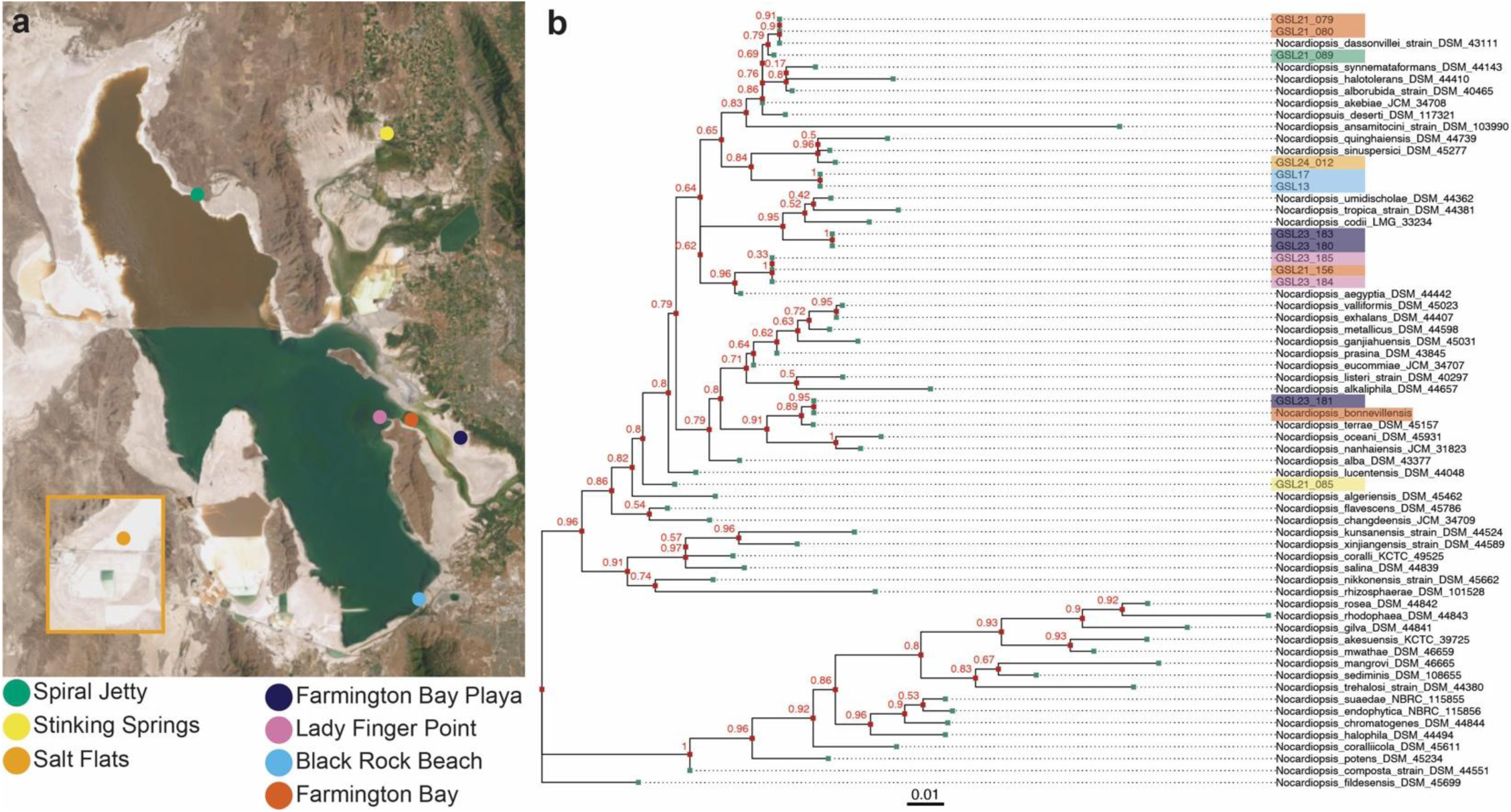
Map of GSL-derived *Nocardiopsis* strain sampling sites and their taxonomic distribution **a**, Satellite image of GSL (base image adapted from the Utah Geological Survey Geologic Map Portal). Sampling sites are indicated by colored circles. The inset (orange box) highlights the Salt Flats sampling location. **b**, Maximum likelihood tree of 16S rRNA sequences showing the current 51 validly published and correctly named *Nocardiopsis* spp., *N. bonnevillensis*, and 13 additional *Nocardiopsis* strains isolated from sediment collected from Great Salt Lake and the surrounding Great Salt Lake Desert. GSL-derived *Nocardiopsis* strains are color-coded according to their sediment sampling location (color key shown in panel 4a).

Acid-fast staining was positive for four isolates, *Nocardiopsis* spp. GSL13, GSL17, GSL21-085, and GSL23-181, while all other GSL-derived *Nocardiopsis* strains were acid-fast negative (Fig. 5). Among these, GSL23-181 was the most closely related to *N. bonnevillensis* based on both 16S rRNA similarity and protein dendrogram. The remaining three acid-fast positive strains, however, were more distantly related and positioned among different *Nocardiopsis* type strains rather than clustering with *N. bonnevillensis*. This phylogenetic distribution indicates that mycolic acid production is unlikely to represent a trait confined to a single evolutionary lineage within Great Salt Lake but may instead be a more broadly distributed adaptation across the genus.

**Figure 5:**
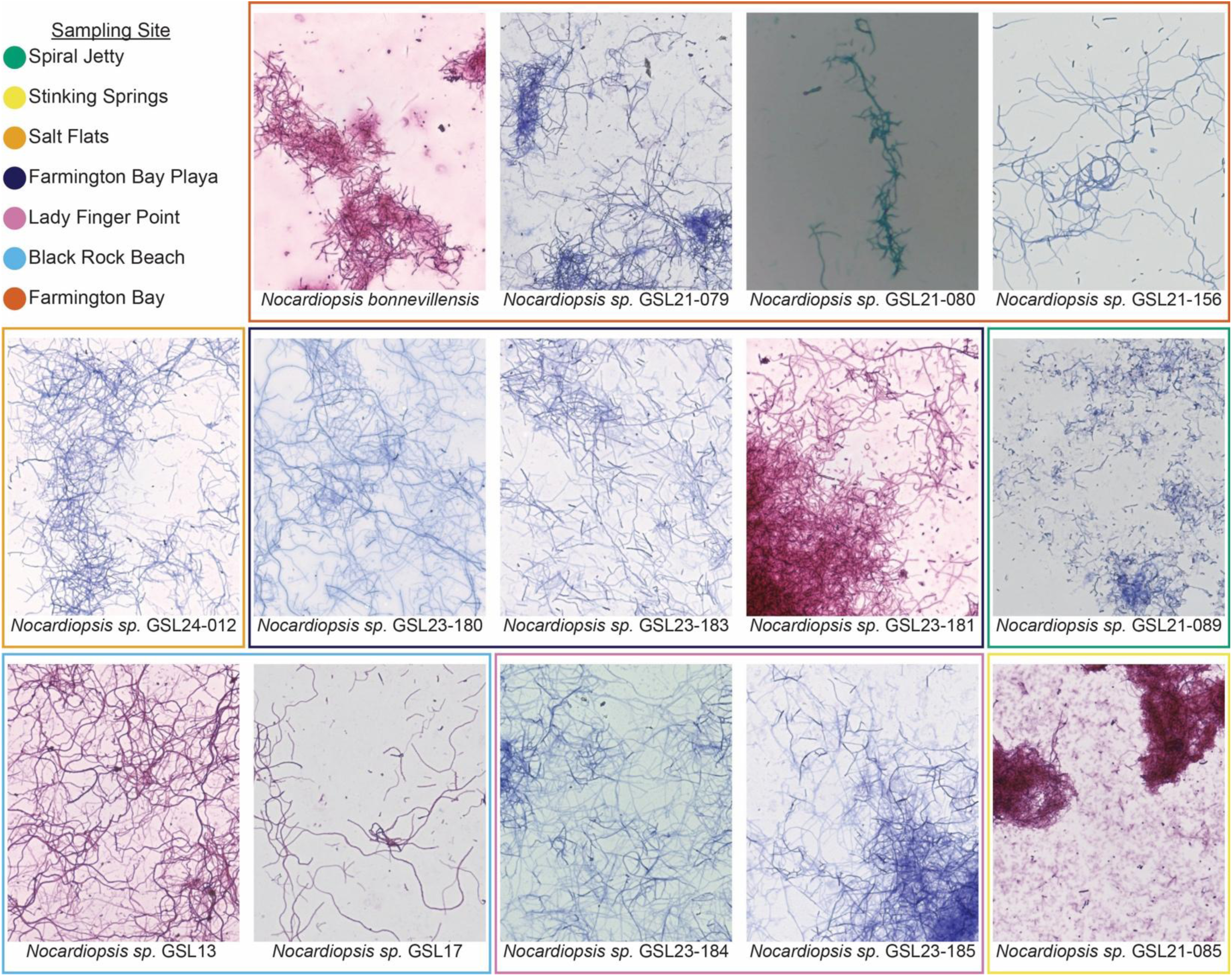
Acid-fast staining of *Nocardiopsis* isolates from Great Salt Lake Acid-fast phenotypes of GSL-derived *Nocardiopsis* spp.. Colored boxes around each image correspond to the sampling site shown in Figure 4a (color key, top left). Images were captured under brightfield microscopy at 40x magnification following modified Ziehl-Neelsen staining. Acid-fast positive isolates (*N. bonnevillensis*, GSL13, GSL17, GSL21-085, and GSL23-181) retain the characteristic red-pink stain, whereas acid-fast negative strains appear blue following counterstaining. Representative images are shown for all 14 isolates examined.

## DISCUSSION

This study identified *Nocardiopsis bonnevillensis* as a novel type strain from Great Salt Lake, notable for producing mycolic acids outside the order *Mycobacteriales* and for encoding diverse and largely uncharacterized biosynthetic gene clusters. Together, these findings expand the known taxonomic distribution of mycolic acid biosynthesis and reveal an unexpected intersection between cell envelope specialization, environmental adaptation, and natural product potential in extremophilic *Actinomycetota*.

Compared to other members of the genus, *N. bonnevillensis* exhibits several distinctive features, including unique menaquinone and phospholipid profiles, acid-fast staining behavior, diagnostic sugars, and differences in carbon source utilization, while genome size, % G+C content, and tolerance for temperature, salinity, and pH remain within typical *Nocardiopsis* ranges. Genomic comparisons revealed 96.1% ANI to *N. terrae* KCTC 19431 and DSM 45157, but digital DNA-DNA hybridization values below the 70% species threshold, indicating clear genomic divergence and supporting its designation as a new species. Chemotaxonomic differences, together with MALDI Biotyper analysis that yielded only genus-level identification, further reinforced the novelty of *N. bonnevillensis* within the *Nocardiopsis* genus.

The detection of mycolic acids in *N. bonnevillensis* challenges a long-held paradigm that these complex fatty acids are restricted to members of the order *Mycobacteriales*. This finding suggests a re-evaluation of their evolutionary history and biological function. The growth defect observed under high-salinity conditions in the presence of isoniazid suggests that mycolic acids contribute to desiccation resistance or osmotic stress tolerance, extending their known role beyond host persistence in pathogens such as *Mycobacterium turberculosis*. Previous work in *Rhodococcus opacus* demonstrated that mycolic acid chain length and saturation decrease in response to NaCl-induced osmotic stress, increasing cell surface hydrophobicity, while *Rhodococcus erythropolis* adjusts fatty acid saturation under pH and temperature extremes^38,39^. These findings, together with our results, suggest modulation of envelope composition may represent a conserved adaptive strategy among diverse *Actinomycetota* inhabiting harsh or stressful environments.

The absence of a FAS-I homolog in *N. bonnevillensis*, despite the presence of all other essential genes for mycolic acid biosynthesis, provides further evidence for evolutionary divergence. Because all previously described mycolic acid-producing bacteria rely on either FAS-I alone (*Corneybacterium*) or both FAS-I and FAS-II (*Mycobacterium*), the absence of FAS-I suggests that the biosynthetic pathway in *N. bonnevillensis* may have arisen through convergent evolution. The presence of GSL acid-fast-positive strains across multiple phylogenetic branches of *Nocardiopsis* supports this hypothesis and argues against a single lineage-specific origin. Instead, the scattered distribution of this phenotype implies either repeated evolutionary gains or widespread ancestral retention followed by differential loss. These findings highlight the limitations of acid-fast staining as a sole indicator of mycolic acid production, particularly since staining outcomes can vary with cell age, growth conditions, and protocol^33,40,41^. This suggests that the historical absence of mycolic acid reports in *Nocardiopsis* may reflect methodological bias rather than biological absence.

The biosynthetic capacity of *N. bonnevillensis* further highlights the ecological and biotechnological potential of extremophilic *Actinomycetota.* Genome mining revealed 19 biosynthetic gene clusters, most with ≤85% similarity to characterized pathways, suggesting significant untapped chemical diversity. Consistent with this analysis, fermentation of *N. bonnevillensis* led to the isolation of bonnevanoside, a thiophenyl nonulopyranoside previously known only as a synthetic compound ^42,43^. Although bonnevanoside lacked antibacterial activity, its structural novelty emphasizes the potential of extremophiles as sources of unprecedented natural product scaffolds. The co-occurrence of unique envelope traits and biosynthetic capacity raises the intriguing possibility that these features may co-evolve in response to environmental pressures, linking chemical novelty with ecological specialization.

Future work will aim to expand the chemical space associated with natural products produced by *N. bonnevillensis,* and to investigate the mechanistic basis and evolutionary trajectory of mycolic acid biosynthesis in non-*Mycobacteriales* lineages. More broadly, these findings emphasize the importance of revisiting classical chemotaxonomic assumptions using modern lipidomics, genome mining and functional assays. By integrating these approaches, we can better understand the diversity, distribution, and ecological significance of mycolate-containing envelopes, and ultimately, uncover new principles that govern microbial survival, evolution, and specialization metabolism in extreme environments.

## METHODS

### Sediment Collection and Bacterial Isolation

A 10 g sediment sample was collected in July 2021 from the shoreline of Great Salt Lake’s Farmington Bay (41° 3’ 58″ N, 112° 13’ 48″ W), located within Antelope Island State Park, Utah using a sterile 15 mL Falcon tube. The sediment was desiccated in a biological safety cabinet for 72 hours, then plated on yeast-peptone-mannitol (YPM) agar supplemented with 4 % (w/v) Instant Ocean (per liter: 4 g mannitol, 2 g yeast extract, 2 g peptone, 18 g agar, and 40 g Instant Ocean Aquarium Sea Salt Mixture, Spectrum Brands, USA). Desiccated sediment was plated on solid YPM media supplemented with 4 % (w/v) artificial sea salt and incubated at 30 °C for 3 weeks. Colonies were isolated by repeated subculturing on the same media, and *N. bonnevillensis* was identified from one of these isolates. Plates were incubated at 30 °C for 3 weeks. Colonies were then sub-plated on the same medium and incubated for 30 °C for 7 to 90 days. Pure isolates were obtained through repeated subculturing, and *N. bonnevillensis* was identified from one of these isolates.

### Sediment Conductivity Measurement

The desiccated sediment sample from which *N. bonnevillensis* was isolated from was diluted 1:5 (w:v) in milliQ water in a 50 mL conical tube, stirred for 30 minutes, and then allowed to settle until the sediment had precipitated out of solution. Conductivity of the supernatant was measured using an Oakton Instruments COND 6+ conductivity meter with a temperature-compensated probe, standardized to report values of 25°C.

### Fermentation studies

To evaluate growth conditions and nutrient tolerance, *N. bonnevillensis* was incubated on YPM agar supplemented with Instant Ocean at concentrations of 0%, 0.1%, 0.5%, and 1–20% (w/v) in 1% increments. Plates were incubated at nine temperatures ranging from 4–45°C. Additional growth assays were performed on YPM agar supplemented with 8% Instant Ocean and adjusted to pH 5–11. *N. bonnevillensis* was also grown on 8% Instant Ocean supplemented ISP2-5 agar (HiMedia Laboratories Private Limited, Maharashtra, India), Czapek-Dox agar, and Potato Dextrose agar (Difco Laboratories, Sparks, MD, USA). All growth condition and nutrient tolerance assays were performed in triplicate and incubated for 14 days at 30 °C unless otherwise specified. Plates were inoculated as previously described^44^.

### Whole genome sequencing

Genomic DNA (gDNA) was extracted from *N. bonnevillensis* using a FastDNASpin Kit (MP Biomedicals, Cat.No.:116540-600). A total of 1.2 µg of gDNA was submitted to Plasmidsaurus (Louisville, Kentucky, USA) for whole-genome sequencing. Sequencing was performed using Oxford Nanopore Technology with custom annotation and analysis provided by Plasmidsaurus.

### Phylogenomic characterization

Genome completeness and contamination were assessed using CheckM^23^. Phylogenomic analyses, including ANI and dDDH were performed using autoMLST^24^, fastANI (v1.33)^45^, and the Type Strain Genome Server^25,46^. The fastANI analysis was run with the parameter --fragLen 1000, while default settings were used for autoMLST and dDDH analyses. Reference *Nocardiopsis* genomes were downloaded using NCBI Datasets (v16.4.3)^26^. For downstream analyses, only genomes exhibiting ≥ 82% ANI to *N. bonnevillensis* were retained (110 genomes total). From each genome, 16S ribosomal RNA (rRNA) sequences were extracted using SeqForge (v0.0.1)^34^ with the 16S rRNA sequence from *N. coralli* (NCBI accession NR_181413.1) used as the query. Alignment of 16S rRNA sequences was performed using MUSCLE (v5.2)^47^, and a maximum likelihood phylogenetic tree was constructed using RAxML-NG (v1.2.2)^48^ with the following parameters: --all, --model GTR+G, and --bs-trees 2000. Branch support values were added using the transfer bootstrap expectation (TBE) method with --support and --bs-metric tbe. Whole-genome phylogenomic analysis was performed using PhyloPhlAn (v3.1.68)^27^. Marker gene amino acid sequences identified by PhyloPhlan were aligned and used to generate a maximum-likelihood tree with raxml-ng using the following parameters: --all, --model LG+G8+F, --tree pars{10}, and --bs-trees 200. Branch support was added using the TBE method as with the 16S rRNA tree. All phylogenetic trees were constructed and analyzed in R using the ggtree^49^ library. The multi-genus maximum likelihood tree (Fig. 1b) was constructed using 16S rRNA sequences extracted from type strains downloaded from the NCBI taxonomic database. From each genome, 16S rRNA sequences were extracted using SeqForge (v0.0.1)^34^ using the 16S sequence from *N. coralli* (NR_181413.1) as a BLAST query. This resulted in incomplete 16S sequence extractions from *N. prasina*, *D. cercidiphylli*, and *T. strandjordii*. To ensure all 16S sequences were of equal length, full-length sequences from these three strains were downloaded from the NCBI taxonomy database (Supplementary Table 8). The maximum likelihood tree was generated via RAxML using 25 starting parsimony and 25 starting random trees, 1000 bootstraps, a fixed seed of 12345, and the GTR+G model of nucleotide substitution. The final tree was visualized with iTOL.^50^

### Chemotaxonomic characterization

*N. bonnevillensis* was cultivated in A1 medium at 30 °C for cellular fatty acids analysis. For all other tests conducted by DMSZ, the strain was grown on M541 medium at 30 °C. Cellular fatty acids were converted to fatty acid methyl esters (FAMES) via saponification, methylation, and extraction^51^. FAMEs were analyzed by gas chromatography with flame ionization detection (GC-FID) followed by gas chromatography mass spectrometry (Agilent GC-MS 7000D system), and peaks were identified by comparing retention times and mass spectra to reference standards^52^. Respiratory quinones were extracted with hexanes from ∼50 mg of freeze-dried biomass, purified by a silica-based solid phase extraction, and detected by high performance liquid chromatography diode array detection mass spectrometry (HPLC-DAD-MS) using a reverse phase column^52^. Polar lipids were extracted from ∼200 mg of freeze-dried biomass using a choroform:methanol:0.3% aqueous NaCl mixture. Lipids were recovered from the chloroform phase^51^ and separated by two-dimensional thin-layer chromatography on silica gel plates. The first dimension was developed in chloroform:methanol:water, and the second in chloroform:methanol:acetic acid:water. Total lipids were visualized by spraying with 5% ethanolic molybdatophosphoric acid followed by heating at 140 °C ^52^. Mycolic acids were extracted from ∼300 mg of wet biomass and analyzed by HPLC as described previously^53^. Extracts were dried and reconstituted in chloroform:methanol, then analyzed by quadrupole time-of-flight mass spectrometry (QTOF-MS, Agilent) via direct infusion into an electrospray ionization (ESI) source. Mycolic acids were identified by comparing exact masses to those of known mycolic acid standards. Whole-cell hydrolysates were prepared using 1N H_2_SO_4_(100 °C, 2 hours) from ∼15 mg of freeze-dried cells to detect diagnostic sugars. Sugars were analyzed by thin layer chromatography (TLC) on cellulose plates as previously described^54^. Polyamines were extracted from ∼50 mg of wet biomass, derivatized, and analyzed by GC-MS as previously described^55,56^. The following polyamines and precursors were screened: agmatine, cadaverine, homospermidine, norspermidine, 1,2- and 1,3-diaminopropane, putrescine, *N*-acetyl-putrescine, 2-hydroxyputrescine, spermidine, and spermine. For detection of diaminopimelic acid (Dpm) isomers, including 2,6-diamino-3-hydroxypimelic acid (OH-Dpm), whole cell hydrolysates were prepared with 4N HCl (100 °C, 16 hours) and analyzed by TLC on cellulose plates^57^. Peptidoglycan was isolated and its structure was determined as previously described^57^. Amino acids were derivatized as *N*-heptafluorobutyryl isobutyl esters and identified and quantified by GC-MS based on comparison to authentic standards. Bacterial classification was performed using MALDI-TOF-MS and analyzed with the MALDI Biotyper software (Bruker Daltonics) as previously described^58^. The API Zym (bioMérieux) test was used to evaluate enzymatic activity, and the API CHB (bioMérieux) test was used to assess carbon source utilization.

### Genome mining

BGCs were initially identified using antiSMASH v8.0^28^. BGCs of interest were further analyzed and manually annotated using NCBI Blast/Blast+. To identify genes involved in mycolic acid biosynthesis, a curated list of protein sequences was compiled from 15 representative species (Supplementary Table 7) belonging to genera known to produce mycolic acids. These sequences were retrieved from the Kyoto Encyclopedia of Genes and Genomes (KEGG) ortholog tables for pathway modules M00886 (meromycolic acid biosynthesis) and M00887 (mycolic acid biosynthesis). The sequences were then queried against the *N. bonnevillensis* genome using Blast+. In addition, the genome was functionally annotated using the eggNOG-mapper web server with default parameters^35^. Orthologous genes identified in the *N. bonnevillensis* genome were aligned with their corresponding query sequences using the Clustal Omega multiple sequence alignment tool provided by the European Bioinformatics Institute (EBI)^59^. Alignments metrics for each homolog were obtained using NCBI Blast by comparing the *N. bonnevillensis* sequences with their highest-scoring matches. All gene annotations were documenting using SnapGene software (v.7.1.1).

### Isolation of compound 1

*N. bonnevillensis,* inoculated from a 10 mL seed culture, was cultivated in a 2.4 L Ultra Yield flask (Thomson Scientific, Oceanside, CA) containing 1 L of A1 broth (10 g soluble startch, 4 g yeast extract, 2 g peptone, 750 mL natural seawater, 250 mL distilled water) at 180 rpm and 27 °C for 3 days. After which, 25 mL of culture was transferred into ten 2.5 L Fernbach flasks, each containing 1 L of A1 broth. After 7 days of cultivation under the same conditions, sterile XAD-7 resin (25 g) was added to each flask and incubated for 3 hrs at 27 °C with constant shaking at 180 rpm. Cultures were then filtered through cheescloth, and the cells and resin were extract with acetone (2 L, for 4 h at 27 °C and constant shaking at 180 rpm). The extract was filtered through cotton wool, and the acetone was removed by rotary evaporation. The resulting crude extract (1.2 g) was partitioned between ethyl acetate and H_2_O (1:1, 600 mL total). The organic layer was collected, dried over anhydrodous Na_2_SO_4_, and concentrated via rotary evaporation. The extract was subjected to silica gel column chromatography and eluted with a step gradient of *n*-hexane/ethyl acetate/MeOH (1:0:0, 10:1:0, 3:1:0, 1:1:0, 1:5:0, 0:0:1) to afford six fractions (Supplemental Table 4). Fraction 4 (212 mg) was further purified by semi-preparative HPLC (Phenomenex Luna C18(2), 10 µm, 100 Å, 250 × 10 mm column; mobile phase: 30–50% MeCN in H2O with 0.02% TFA over 25 min, yielding compound **1** (t_R_ 25 min, 2.0 mg). The molecular formula of compound **1** was established as C₂₆H₃₃NO₁₂S based on HR-ESI-TOF-MS, which displayed a protonated molecule at m/z 584.1935 ([M + H]⁺; calcd for C₂₆H₃₄NO₁₂S, 584.1802). Compound **1** was obtained as a colorless solid.

### Acid-fast staining

Acid-fast staining was performed using the Ziehl–Neelsen method as previously described^31^. A modified Ziehl–Neelsen method optimized for weakly acid-fast specimens was also used^32^. Slides were examined at 10x and 20x magnification using an Olympus model CKX41SF microscope and at 40x using an Echo Revolution microscope imaged with the brightfield camera. To assess the impact of isoniazid (INH) on growth and acid-fastness, *N. bonnevillensis* was grown in triplicate on solid YPM supplemented with 8% Instant Ocean (w/v) and seven concentrations of INH (12.5–400 µg/mL) or a no-drug control. Plates were incubated at 30 °C for 5 days. For evaluating INH’s effect on cell wall composition, cultures were grown in liquid YPM media containing 8% Instant Ocean (w/v) and supplemented with 0, 12.5, or 25 µg/mL INH at 30 °C with shaking (180 RPM) for 5 days. Cells from liquid cultures were processed for acid-fast staining as described above. To evaluate the effect of INH on salinity tolerance, YPM agar was prepared with 0, 4, 8, 12, or 16% Instant Ocean (w/v) and further supplemented with INH to final concentrations of 0, 12.5, 25, or 200 µg/mL. Plates were inoculated as described above, incubated at 30 °C for 14 days, and photographed on days 5, 6, 10, and 14 (Supplemental Fig. 14).

### Isolation and identification of additional *Nocardiopsis* strains

Sediment samples were collected as described previously from Black Rock Beach (40°44′15″ N, 112°12′10″ W), Farmington Bay Playa (41° 2’ 57″ N, 112° 6’ 40″ W), Lady Finger Point (41° 3’ 21″ N, 112° 15’ 15″ W), Salt Flats (40° 44′ 31″ N, 113° 50′ 48″ W), Spiral Jetty (41° 26’ 17″ N, 112° 39’ 57″ W), and Stinking Springs (41° 34’ 37″ N, 112° 13’ 60″ W). Isolates were obtained from GSL sediment samples as described above. Spore PCR was performed to amplify 16S sequences using 27F and 1492R primers (Supplementary Table 9). Amplicons were sequenced using Oxford Nanopore Technologies-based linear amplicon sequencing (Plasmidsaurus). 16S rRNAsequences were queried using NCBI Blast for genus identification and their taxonomic placements were determined as described above.

### Matrix Assisted Laser Desorption Ionization-Time of Flight (MALDI-TOF) analysis of GSL-derived *Nocardiopsis* strains

Cultures of the GSL-derived *Nocardiopsis* strains on YPM agar supplemented with 5 or 8% salinity were analyzed by MALDI along with the corresponding media controls. Protein and metabolite spectra were processed using the IDBac platform^37^. Following incubation, a small volume of colony cell-mass was applied to designated spots of a 384-spot MALDI target plate (Bruker Daltonics, Billerica, MA, USA) using sterile toothpicks. Five biological replicates were plated per condition. Following cell-mass application, 1 μl of 70% formic acid (7:3 Optima, Fisher Chemical: Optima LC-MS Grade Water Fisher Chemical) was added to each well followed by an overlay of matrix (α-cyano-4-hydroxycinnamic acid [powder, 98% pure, Sigma-Aldrich, part-C2020] 50% acetonitrile, 47.5% water, and 2.5% trifluoroacetic acid). Solvents used in this preparation were LC-MS grade. This procedure has been previously published in detail^37^. MALDI-TOF MS data collection was conducted on a rapifleX MALDI Tissuetyper mass spectrometer (Bruker Daltonics) equipped with a smartbeam™ 3D laser (355 nm). Automated data acquisitions were performed using flexControl software v. 4.0.46.0 (Bruker Daltonics) and flexAnalysis software v. 3.4. Small molecule spectra were obtained using positive reflectron mode (1000 shots; RepRate: 5000 Hz; delay: 28272 ns; ion source 1 voltage: 20 kV; ion source 2 voltage: 18.45 kV; lens voltage: 9 kV; mass range: 60 Da to 2700 Da). Spectra were corrected with external Bruker Daltonics Peptide Calibration Standard.

## DATA AVAILABILITY

The genome of *N. bonnevillensis* is deposited in GenBank under BioProject accession number #. The GSL-derived *Nocardiopsis* 16S rRNA sequences are deposited under BioProject accession number #. *N. bonnevillensis* is deposited in the DSMZ open collection as DSM 118115.

## ACKNOWLEDGEMENTS

We thank Drs. Lou Barrows (University of Utah) and Allison Carey (University of Utah) for helpful advice and discussions on mycolic acid mass spectrometry results, Martha Trujillo (University of Salamanca) for helpful discussions about *Nocardiopsis* taxonomy, and Ana Beatriz DePaula-Silva (University of Utah) for microscopy assistance. We thank Giulia Napoli (University of Utah) for staining assistance. The authors gratefully acknowledge the Leibniz Institute DSMZ–German Collection of Microorganisms and Cell Cultures GmbH for their expertise and assistance with the chemotaxonomic characterization of *N. bonnevillensis*. This work was supported by a Skaggs Foundation for graduate research fellowship to AFS; a 3i graduate research fellowship to ERBH; the University of Utah Research Foundation, the Ben and Iris Margolis Foundation, and the National Institutes of Health [1R01AI155694] to JMW, the Chicago Biomedical Consortium with support from the Searle Funds at The Chicago Community Trust to BTM and Dr. Neil Kelleher, and in part by the National Cancer Institute CA044848 to WF and the Moore Foundation GBMF7621 to JMW.

## Author Contributions

### Conceptualization of the study

A.F.S. and J.M.W. Methodology: A.F.S., T.J., E.R.B.H., N.K.K., R.I., and A.B. Data analysis: all authors. Project administration: J.M.W. Writing–original draft: A.F.S. and J.M.W. Writing– review and editing: all authors.

### Ethics Declarations

The authors declare no competing interests.

## Extended Data

**Extended Data Figure 1:**
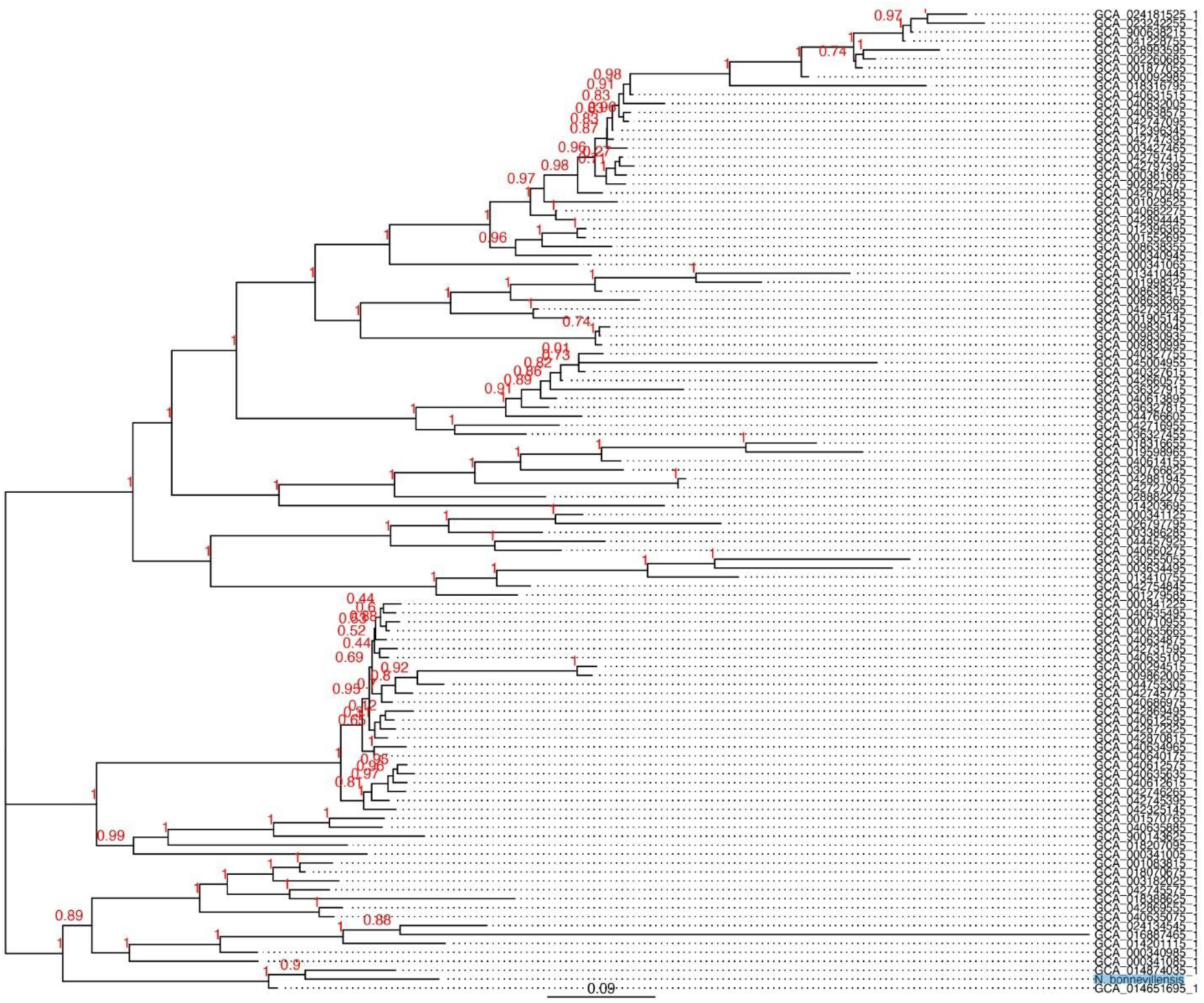
Whole-genome phylogeny of *N. bonnevillensis* and related *Nocardiopsis* genomes A total of 360 conserved single-copy marker genes were identified across a library of *Nocardiopsis* genomes using PhyloPhlAn^27^. Genomes exhibiting ≥ 82% ANI to *N. bonnevillensis* (n= 110 genomes) were included in the downstream phylogenomic analysis. Marker gene amino acid sequences were aligned in PhyloPhlAn and used to construct a maximum-likelihood tree with RAxML^48^ based on 200 bootstrap replicates.

**Extended Data Table 1:** Chemotaxonomic analyses of N. bonnevillensis. Polar lipids identified: diphosphatidylglycerol (DPG), phosphatidylglycerol (PG), aminophospholipid (APL), glycophospholipid (GPL), two unidentified glycolipids (GL), and three unidentified phospholipids (PL). Peptidoglycan hydrolysate contained diaminopimelic acid (Dpm).

| Cellular components | Composition |
| --- | --- |
| Major Fatty Acids (>20%) | 10-methyl-C<sub>18:0</sub> (24.8%), iso-C<sub>16:0</sub> (23.7%), anteiso-C<sub>17:0</sub> (22.1%). |
| Minor Fatty Acids (>1%) | cis-9-C<sub>18:1</sub> (5.1%), anteiso-C<sub>15:0</sub> (4.0%), iso-C<sub>18:0</sub> (3.0%), C<sub>18:0</sub> (2.6%), iso-cis-9-C<sub>18:1</sub> (2.4%), iso-C<sub>17:0</sub> (2.3%), 10-methyl-C<sub>17:0</sub> (1.9%), 10-methyl-C<sub>16:0</sub> (1.2%), iso-C<sub>14:0</sub> (1.2%), anteiso-C<sub>19:0</sub> (1.1%) |
| Additional Fatty acids (<1%) | anteiso-C<sub>13:0</sub>, C<sub>14:0</sub>, iso-C<sub>15:0</sub>, cis-9-C<sub>16:1</sub>, C<sub>16:0</sub>, anteiso-cis-9-C<sub>17:1</sub>, cis-9-C<sub>17:1</sub>, C<sub>17:0</sub>, iso-cis-9-C<sub>19:1</sub>, iso-C<sub>19:0</sub>, C<sub>19:0</sub>, iso-C<sub>20:0</sub>, C<sub>20:0</sub> |
| Menaquinones | MK-9 H<sub>2</sub> (1.0%), MK-9 H<sub>4</sub> (4.4%), MK-9 H<sub>6</sub> (4.9%), MK-9 H<sub>8</sub> (5.3%) |
| polar lipid profile | DPG, PG, APL, GPL, two unidentified GL, three unidentified PL |
| whole-cell sugars | glucose, ribose |
| peptidoglycan hydrolysate ratio | 3.9 Ala: 1.0 Glu : 0.3 Dpm |
| peptidoglycan hydrolysate peptides | M-Ala, Ala-Glu, Glu-Dpm, Dpm-Ala-Dpm |
| peptidoglycan type | A1γ |

**Extended Data Table 2:**
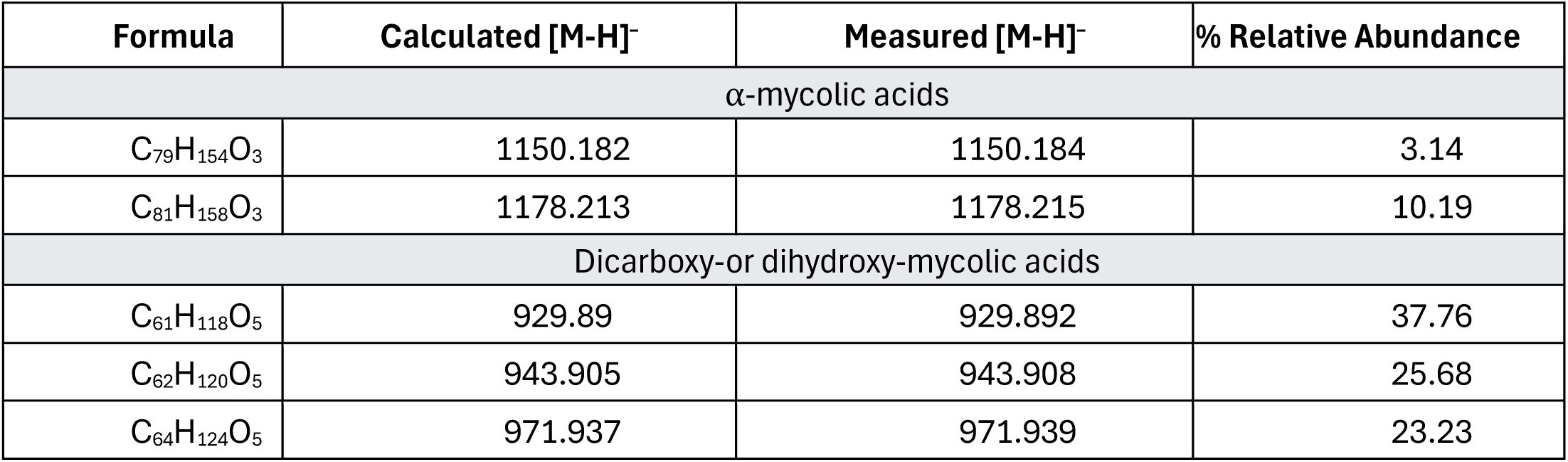
Detection of mycolic acids in N. bonnevillensis. Masses detected in biomass extracts of N. bonnevillensis by ESI-QTOF-MS were compared with standards of known mycolic acids.

| Formula | Calculated [M-H]<sup>-</sup> | Measured [M-H]<sup>-</sup> | % Relative Abundance |
| --- | --- | --- | --- |
| α-mycolic acids |  |  |  |
| C<sub>79</sub>H<sub>154</sub>O<sub>3</sub> | 1150.182 | 1150.184 | 3.14 |
| C<sub>81</sub>H<sub>158</sub>O<sub>3</sub> | 1178.213 | 1178.215 | 10.19 |
| Dicarboxy-or dihydroxy-mycolic acids |  |  |  |
| C<sub>61</sub>H<sub>118</sub>O<sub>5</sub> | 929.89 | 929.892 | 37.76 |
| C<sub>62</sub>H<sub>120</sub>O<sub>5</sub> | 943.905 | 943.908 | 25.68 |
| C<sub>64</sub>H<sub>124</sub>O<sub>5</sub> | 971.937 | 971.939 | 23.23 |

**Extended Data Table 3:** Enzymatic activity profile of N. bonnevillensis by API Zym. Enzymatic activities were assessed after 6 hour incubation at 37 °C. Activity scoring: strong (+), 3–5; weak (w), 1–2; negative (-), 0–0.5.

| Enzyme | Activity | Score |
| --- | --- | --- |
| alkaline phosphatase | + | 5 |
| leucine-arylamidase | + | 5 |
| esterase | w | 1 |
| α-glucosidase | w | 1 |
| esterase lipase | - | 0 |
| lipase | - | 0 |
| valine-arylamidase | - | 0.5 |
| cystine-arylamidase | - | 0 |
| trypsin | - | 0 |
| chymotrypsin | - | 0.5 |
| acid phosphatase | - | 0 |
| naphthol-AS-BI-phosphohydrolase | - | 0.5 |
| α-galactosidase | - | 0 |
| β-galactosidase | - | 0 |
| β-glucuronidase | - | 0 |
| N-acetyl-β-glucosaminidase | - | 0.5 |
| α-mannosidase | - | 0 |
| α-fucosidase | - | 0 |

**Extended Data Table 4:** Carbon source utilization of N. bonnevillensis by API 50 CHB. Carbon source utilization was assessed using the API 50 CHB assay strip after incubation at 30 °C for 5 days. Activity scoring: strong (+), weak (w), negative (-).

| Carbon Source | Utilization |
| --- | --- |
| glycerol | + |
| L-arabinose | + |
| ribose | + |
| D-xylose | + |
| glucose | + |
| rhamnose | + |
| esculine | + |
| cellobiose | + |
| starch | w |
| glycogen | w |
| D-arabitol | w |
| erythritol | - |
| D-arabinose | - |
| L-xylose | - |
| adonitol | - |
| β-Me-D-xyloside | - |
| galactose | - |
| fructose | - |
| mannose | - |
| sorbose | - |
| dulcitol | - |
| inositol | - |
| mannitol | - |
| sorbitol | - |
| α-Me- D-mannoside | - |
| α-Me-glucoside | - |
| N-acetylglucosmine | - |
| amygdalin | - |
| arbutin | - |
| salicin | - |
| maltose | - |
| lactose | - |
| melibiose | - |
| sucrose | - |
| trehalose | - |
| insulin | - |
| melizitose | - |
| raffinose | - |
| xylitol | - |
| gentibiose | - |
| D-turanose | - |
| D-lyxose | - |
| D-tagatose | - |
| D-fucose | - |
| L-fucose | - |
| L-arabitol | - |
| gluconate | - |
| 2-ketogluconate | - |
| 5-keogluconate | - |

**Extended Data Figure 2:**
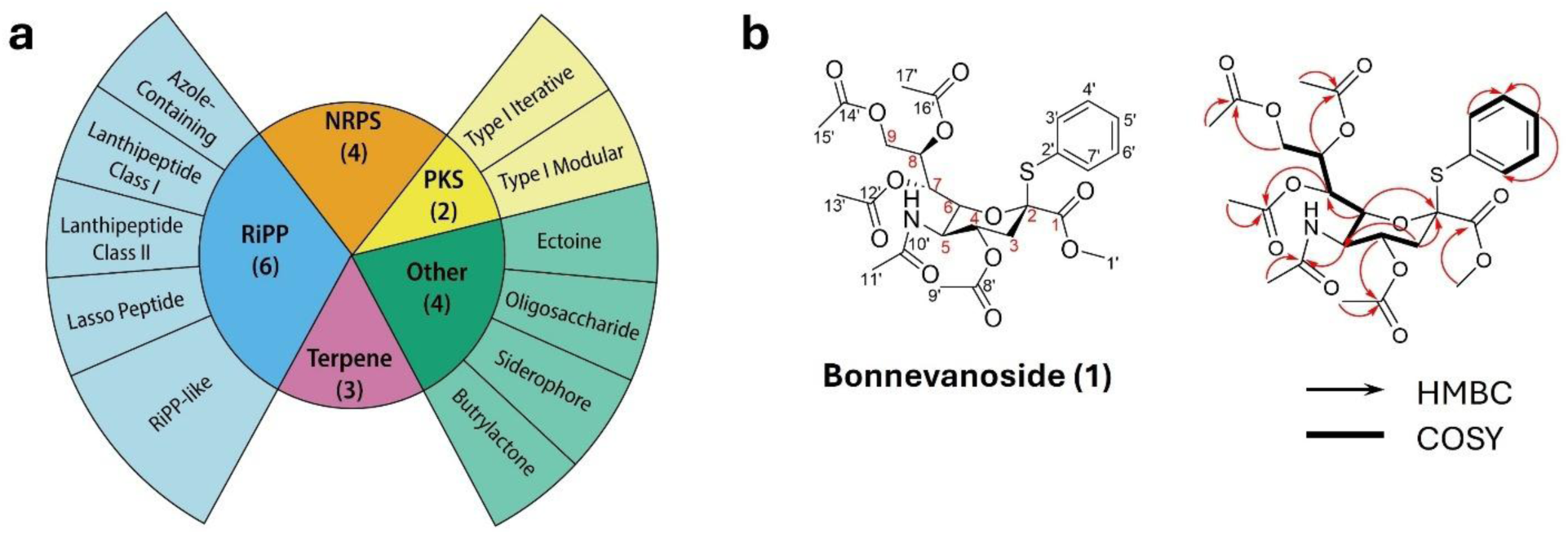
Biosynthetic gene clusters predicted in *N. bonnevillensis* and structure of bonnevanoside. **a**, Number of BGCs predicted from the *N. bonnevillensis* genome, grouped by natural product class (center) and subclass (outer ring). **b**, Molecular structure of bonnevanoside (**1**) and key COSY and HMBC correlations used to determine the structure of **1**.

**Extended Data Figure 3:**
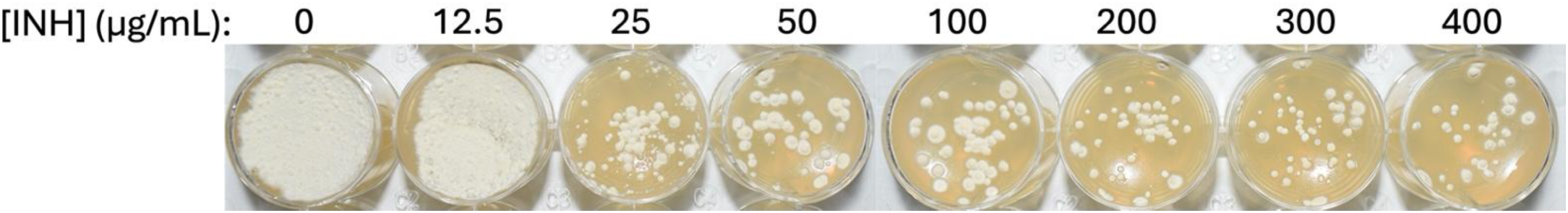
*N. bonnevillensis* after five days of growth on YPM media containing 8% salinity and increasing concentrations of INH.

**Extended Data Figure 4:**
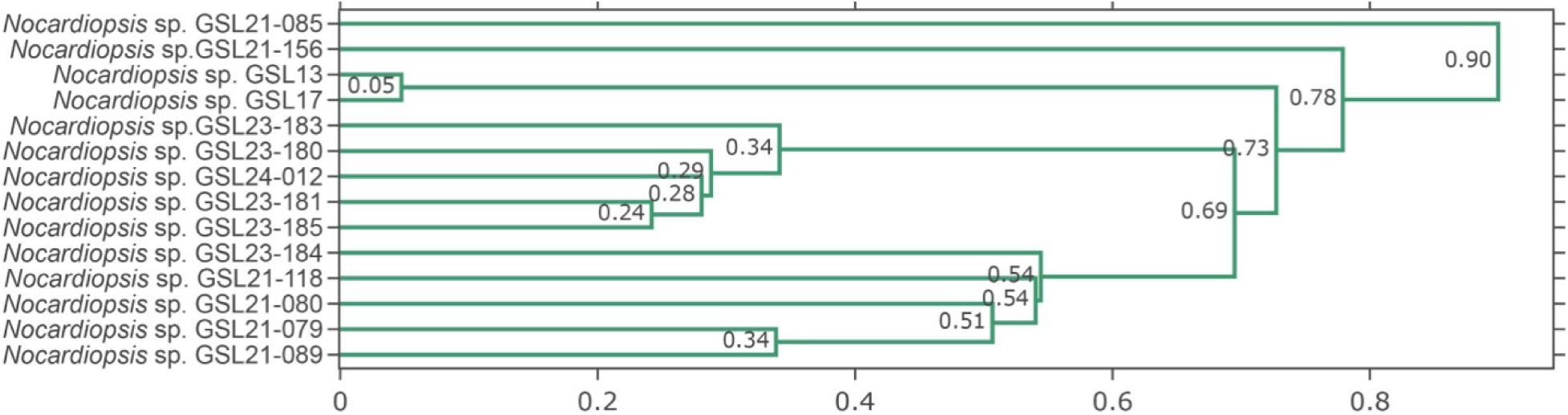
Protein dendrogram of 14 *Nocardiopsis* strains isolated from Great Salt Lake sediment. A protein dendrogram was generated from MALDI-TOF MS protein mass spectra (3–20 kDa) and visualized using the IDBac analysis pipeline and visualized with the IDBac^37^ Interactive Analysis Protein Dendrogram web analysis platform.

**Extended Data Table 5:**
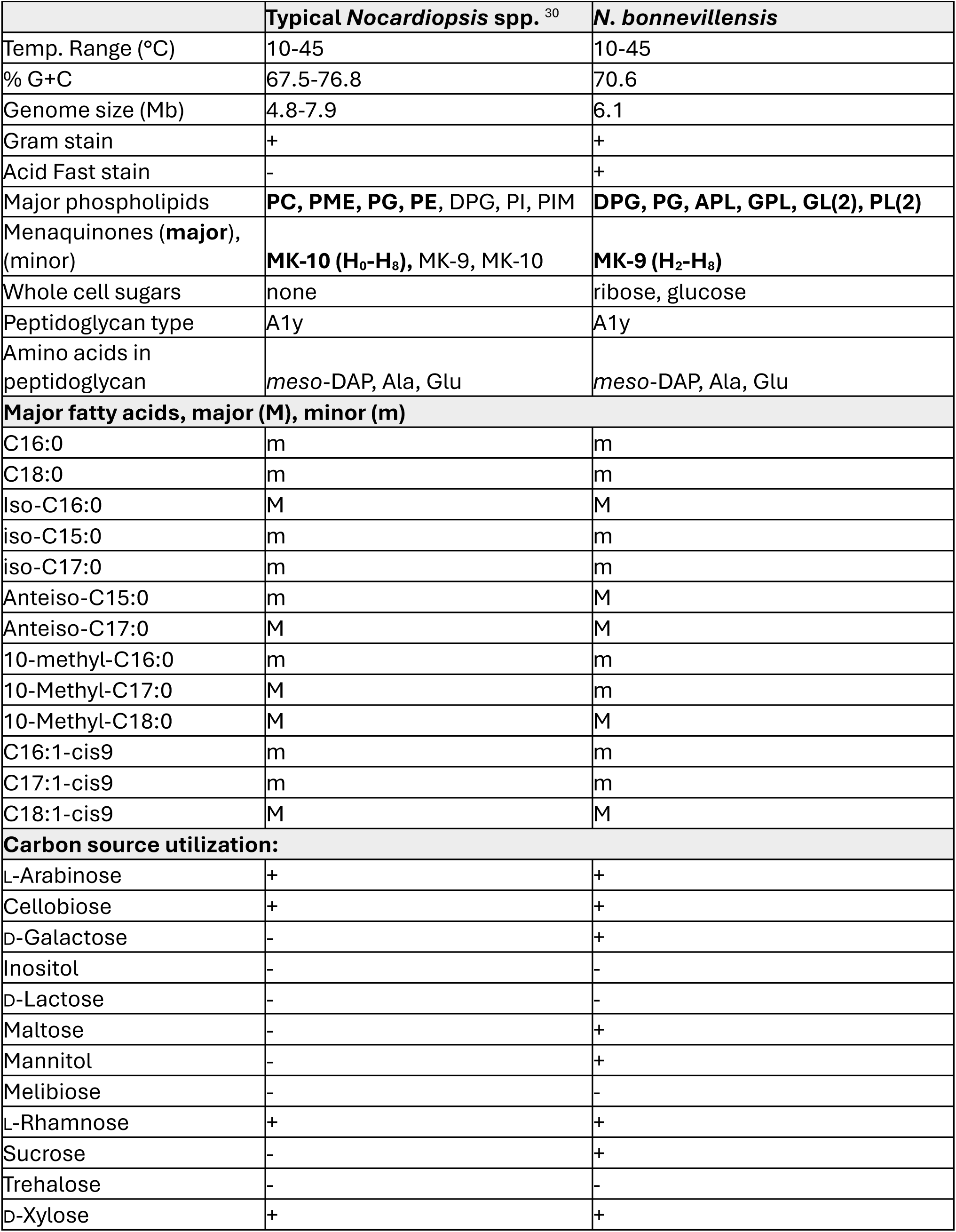
Chemotaxonomic characteristics distinguishing N. bonnevillensis from typical Nocardiopsis species. Carbon utilization pattens are compared with those of typical Nocardiopsis spp., where “+” indicates positive utilization reported for the majority of strains (Hozzein and Trujillo 2015).

**Extended Data Table 6:** Chemotaxonomic characteristics distinguishing N. bonnevillensis from N. terrae KCTC 19431. Comparisons include genomic features, physiological ranges, diagnostic sugars and amino acids, phospholipids, menaquinones, fatty acids, culture characteristics, and carbon source utilization.

| Characteristic | *N. bonnevillensis* | *N. terrae* KCTC 19431<sup>60</sup> |
| --- | --- | --- |
| Genome size (Mb) | 6.1 | 6 |
| % G+C | 70.6 | 71.5 |
| % NaCl tolerance | 0-16 | 1-15 |
| pH range | 5-11 | 6-10.5 |
| Temp Range (°C) | 10-45 | 10-45 |
| Diagnostic sugars | ribose, glucose | none |
| Diagnostic diamino acid | meso-Dpm | meso-Dpm |
| Major phospholipids | DPG, PG, APL, GPL, GL(2), PL(2) | DPG, PC, PG, PME |
| Major menaquinones | MK-9(H<sub>2</sub>-H<sub>8</sub>) | MK-9(H<sub>8</sub>), MK-10(H<sub>4</sub>-H<sub>8</sub>) |
| Major fatty acids (in bold): |  |  |
| 10-methyl-C<sub>18:0</sub> | **24.8** | **14.3** |
| iso-C<sub>16:0</sub> | **23.7** | **36.5** |
| anteiso-C<sub>17:0</sub> | **22.1** | **18.2** |
| C<sub>18:1</sub>ω<sub>9c</sub> | **5.1** | **3.5** |
| anteiso-C<sub>15:0</sub> | **4.0** | 2.0 |
| 10-methyl C<sub>17:0</sub> | 1.9 | **10.8** |
| Culture Characteristics on various media<sup>a</sup>: |  |  |
| ISP2 | Poor | Good* |
| ISP3 | Good | Moderate* |
| ISP4 | Good* | Good |
| ISP5 | Good | Poor* |
| Czapek-Dox | Moderate | Good* |
| Potato Dextrose | Poor | Moderate |
| Utilization of carbon sources<sup>b</sup>: |  |  |
| Glycerol | + | + |
| L-Arabinose | + | - |
| Ribose | + | + |
| D-Xylose | + | + |
| Adonitol | - | - |
| Galactose | - | - |
| Glucose | + | + |
| Fructose | - | + |
| Mannose | - | + |
| Rhamnose | + | - |
| Dulcitol | - | + |
| Inositol | - | - |
| D-Mannitol | - | - |
| Salicilin | - | - |
| Cellulobiose | + | + |
| Maltose | - | - |
| Lactose | - | - |
| Melibiose | - | - |
| Sucrose | - | + |
| Trehalose | - | + |
| Melizitose | - | - |
| Raffinose | - | + |
| D-Arabitol | w | - |
| Gluconate | - | + |

\*Soluble pigment observed.
<sup>a</sup>For direct comparison between strains with media conditions, NaCl was used instead of Instant Ocean. All media conditions with *N. bonnevillensis* were supplemented with 8% NaCl (w/v) and *N. terrae* was supplemented with 4% NaCl (w/v)
<sup>b</sup>positive (+), weakly positive (w), negative (-)

