## Supplemental Information for "An extremophilic *Nocardiopsis* strain from Great Salt Lake expands the taxonomic range of mycolic acid biosynthesis"

**Table of Contents**

**Supplementary Information	Page Number**

16S sequences of GSL-derived *Nocardiopsis* strains	S3

**Supplementary Results**

Supplementary Result 1: *N. bonnevillensis* growth across NaCl, temperature, and pH ranges	S11

Supplementary Result 2: Catalase and oxidase activity of *N. bonnevillensis*	S14

Supplementary Result 3: Biosynthetic gene cluster identification by genome mining	S15

Supplementary Result 4: Antibacterial activity of *N. bonnevillensis*	S16

Supplementary Result 5: Isolation of bonnevanoside	S17

**Supplementary Figures**

Figure S1. Maximum likelihood tree of 16S rRNA sequences from *Nocardiopsis* genomes	S10

Figure S2. Growth of *N. bonnevillensis* across temperature and NaCl ranges	S11

Figure S3. Metabolic Association Network (MAN) of *N. bonnevillensis*	S13

Figure S4. Negative HR-ESI-TOF-MS and UV spectra of compound **1**	S17

Figure S5. Structure and key 2D NMR NOESY correlations of compound **1**	S18

Figure S6. ^1^H NMR spectrum of compound **1** at 500 MHz in MeOD	S20

Figure S7. ^13^C NMR spectrum of compound **1** at 125 MHz in MeOD	S21

Figure S8. HSQC spectrum of compound **1** at 500 MHz in MeOD	S22

Figure S9. HMBC spectrum of compound **1** at 500 MHz in MeOD	S22

Figure S10. COSY spectrum of compound **1** at 500 MHz in MeOD	S23

Figure S11. NOESY spectrum of compound **1** at 500 MHz in MeOD	S23

Figure S12. ^1^H NMR spectrum of compound **1** at 500 MHz in CDCl_3_	S24

Figure S13. TOCSY spectrum of compound **1** at 500 MHz in CDCl_3_	S24

Figure S14: Growth of *N. bonnevillensis* across isonizid and salinity ranges	S29

**Supplementary Tables**

Table S1. Quality and color observations for *N. bonnevillensis* grown on standard media	S12

Table S2. antiSMASH summary of biosynthetic gene clusters identified in *N. bonnevillensis* 	S15

Table S3. Antimicrobial activity of crude extract from *N. bonnevillensis*	S16

Table S4. Antimicrobial activity of fractionated extracts from *N. bonnevillensis*	S16

Table S5. NMR spectral data of **1**	S19

Table S6. Antibacterial activity of **1**	S20

Table S7. List of known mycolate-producing species	S25

Table S8. Reference genomes used to create Fig. 1b.	S26

Table S9: 16S rRNA primers	S28

**Supplementary References**	S32

**Supplementary Information:** 16S rRNA sequences of GSL-derived *Nocardiopsis* strains

***Nocardiopsis bonnevillensis***

GACGAACGCTGGCGGCGTGCTTAACACATGCAAGTCGAGCGGTAAGGCCCTTCGGGGTACACGAGCGGCGAACGGGTGAGTAACACGTGAGCAACCTGCCCCTGACTCCGGGATAAGCGGTGGAAACGCCGTCTAATACCGGATATGACACACCATCTCCTGGTGGTGTGTGGAAAGTTTTGTCGGTCAGGGATGGGCTCGCGGCCTATCAGCTAGTTGGTGGGGTAAAGGCCTACCAAGGCGATTACGGGTAGCCGGCCTGAGAGGGCGACCGGCCACACTGGGACTGAGACACGGCCCAGACTCCTGCGGGAGGCAGCAGTGGGGAATATTGCACAATGGGCGAAAGCCTGATGCAGCGACGCCGCGTGGGGGATGACGGCCTTCGGGTTGTAAACCTCTTTTACCACCAACGCAGGCCCCGGGTTCTCTCGGGGTTGACGGTAGGTGGGGAATAAGGACCGGCTAACTACGTGCCAGCAGCCGCGGTAATACGTAGGGTCCGAGCGTTGTCCGGAATTATTGGGCGTAAAGAGCTCGTAGGCGGCGTGTCGCGTCTGCTGTGAAAGACCGGGGCTTAACCCCGGTTCTGCAGTGGATACGGGCATGCTAGAGGTAGGTAGGGGAGACTGGAATTCCTGGTGTAGCGGTGAAATGCGCAGATATCAGGAGGAACACCGGTGGCGAAGGCGGGTCTCTGGGCCTTACCTGACGCTGAGGAGCGAAAGCATGGGGAGCGAACAGGATTAGATACCCTGGTAGTCCATGCCGTAAACGTTGGGCGCTAGGTGTGGGGACTTTCCACGGTTTCCGCGCCGTAGCTAACGCATTAAGCGCCCCGCCTGGGGAGTACGGCCGCAAGGCTAAAACTCAAAGGAATTGACGGGGGCCCGCACAAGCGGCGGAGCATGTTGCTTAATTCGACGCAACGCGAAGAACCTTACCAAGGTTTGACATCACCCGTGGACCTGTAGAGATACAGGGTCATTTAGTTGGTGGGTGACAGGTGGTGCATGGCTGTCGTCAGCTCGTGTCGTGAGATGTTGGGTTAAGTCCCGCAACGAGCGCAACCCTTGTTCCATGTTGCCAGCACGTAATGGTGGGGACTCATGGGAGACTGCCGGGGTCAACTCGGAGGAAGGTGGGGACGACGTCAAGTCATCATGCCCCTTATGTCTTGGGCTGCAAACATGCTACAATGGCCGGTACAATGGGCGTGCGATACCGTAAGGTGGAGCGAATCCCTTAAAGCCGGTCTCAGTTCGGATTGGGGTCTGCAACTCGACCCCATGAAGGTGGAGTCGCTAGTAATCGCGGATCAGCAACGCCGCGGTGAATACGTTCCCGGGCCTTGTACACACCGCCCGTCACGTCATGAAAGTCGGCAACACCCGAAACTTGTGGCCTAACCCTTCGGGGAGGGAATGAGTGAAGGTGGGGCTGGCGATTGGGACG

***Nocardiopsis* sp. GSL13**

GACGAACGCTGGCGGCGTGCTTAACACATGCAAGTCGAGCGGTAAGGCCCTTCGGGGTACACGAGCGGCGAACGGGTGAGTAACACGTGAGCAACCTGCCCCCGACTCCGGGATAAGCGGTGGAAACGCCGTCTAATACCGGATACGACCTCGCATCTCCTGGTGCGGGGTGGAAAGTTCTTCGGTCGGGGATGGGCTCGCGGCCTATCAGCTTGTTGGTGGGGTAACGGCCTACCAAGGCGATTACGGGTAGCCGGCCTGAGAGGGCGACCGGCCACACTGGGACTGAGACACGGCCCAGACTCCTGCGGGAGGCAGCAGTGGGGAATATTGCGCAATGGGCGAAAGCCTGACGCAGCGACGCCGCGTGGGGGATGACGGCCTTCGGGTTGTAAACCTCTTTTACCACCAACGCAGGCTCCCGGTTCTCCGGGGGTTGACGGTAGGTGGGGAATAAGGACCGGCTAACTACGTGCCAGCAGCCGCGGTAATACGTAGGGTCCGAGCGTTGTCCGGAATTATTGGGCGTAAAGAGCTCGTAGGCGGCGTGTCGCGTCTGCTGTGAAAGACCGGGGCTTAACTCCGGTTCTGCAGTGGATACGGGCATGCTAGAGGTAGGTAGGGGAGACTGGAATTCCTGGTGTAGCGGTGAAATGCGCAGATATCAGGAGGAACACCGGTGGCGAAGGCGGGTCTCTGGGCCTTACCTGACGCTGAGGAGCGAAAGCATGGGGAGCGAACAGGATTAGATACCCTGGTAGTCCATGCCGTAAACGTTGGGCGCTAGGTGTGGGGACTTTCCACGGTTTCCGCGCCGTAGCTAACGCATTAAGCGCCCCGCCTGGGGAGTACGGCCGCAAGGCTAAAACTCAAAGGAATTGACGGGGGCCCGCACAAGCGGCGGAGCATGTTGCTTAATTCGACGCAACGCGAAGAACCTTACCAAGGTTTGACATCACCCGTGGACCTGCAGAGATGTGGGGTCATTTAGTTGGCGGGTGACAGGTGGTGCATGGCTGTCGTCAGCTCGTGTCGTGAGATGTTGGGTTAAGTCCCGCAACGAGCGCAACCCTTATTCCATGTTGCCAGCACGTGATGGTGGGGACTCATGGGAGACTGCCGGGGTCAACTCGGAGGAAGGTGGGGATGACGTCAAGTCATCATGCCCCTTATGTCTTGGGCTGCAAACATGCTACAATGGCCGGTACAGTGGGCGTGCGATGCCGTAAGGTGGAGCGAATCCCTAAAAGCCGGTCTCAGTTCGGATTGGGGTCTGCAACTCGACCCCATGAAGGTGGAGTCGCTAGTAATCGCGGATCAGCAACGCCGCGGTGAATACGTTCCCGGGCCTTGTACACACCGCCCGTCACGTCATGAAAGTCGGCAACACCCGAAACTTGCGGCCTAACCCTTCGGGGAGGGAGTGAGTGAAGGTGGGGCTGGCGATTGGGACG

***Nocardiopsis* sp. GSL17**

GACGAACGCTGGCGGCGTGCTTAACACATGCAAGTCGAGCGGTAAGGCCCTTCGGGGTACACGAGCGGCGAACGGGTGAGTAACACGTGAGCAACCTGCCCCCGACTCCGGGATAAGCGGTGGAAACGCCGTCTAATACCGGATACGACCTCGCATCTCCTGGTGCGGGGTGGAAAGTTCTTCGGTCGGGGATGGGCTCGCGGCCTATCAGCTTGTTGGTGGGGTAACGGCCTACCAAGGCGATTACGGGTAGCCGGCCTGAGAGGGCGACCGGCCACACTGGGACTGAGACACGGCCCAGACTCCTGCGGGAGGCAGCAGTGGGGAATATTGCGCAATGGGCGAAAGCCTGACGCAGCGACGCCGCGTGGGGGATGACGGCCTTCGGGTTGTAAACCTCTTTTACCACCAACGCAGGCTCCCGGTTCTCCGGGGGTTGACGGTAGGTGGGGAATAAGGACCGGCTAACTACGTGCCAGCAGCCGCGGTAATACGTAGGGTCCGAGCGTTGTCCGGAATTATTGGGCGTAAAGAGCTCGTAGGCGGCGTGTCGCGTCTGCTGTGAAAGACCGGGGCTTAACTCCGGTTCTGCAGTGGATACGGGCATGCTAGAGGTAGGTAGGGGAGACTGGAATTCCTGGTGTAGCGGTGAAATGCGCAGATATCAGGAGGAACACCGGTGGCGAAGGCGGGTCTCTGGGCCTTACCTGACGCTGAGGAGCGAAAGCATGGGGAGCGAACAGGATTAGATACCCTGGTAGTCCATGCCGTAAACGTTGGGCGCTAGGTGTGGGGACTTTCCACGGTTTCCGCGCCGTAGCTAACGCATTAAGCGCCCCGCCTGGGGAGTACGGCCGCAAGGCTAAAACTCAAAGGAATTGACGGGGGCCCGCACAAGCGGCGGAGCATGTTGCTTAATTCGACGCAACGCGAAGAACCTTACCAAGGTTTGACATCACCCGTGGACCTGCAGAGATGTGGGGTCATTTAGTTGGCGGGTGACAGGTGGTGCATGGCTGTCGTCAGCTCGTGTCGTGAGATGTTGGGTTAAGTCCCGCAACGAGCGCAACCCTTATTCCATGTTGCCAGCACGTGATGGTGGGGACTCATGGGAGACTGCCGGGGTCAACTCGGAGGAAGGTGGGGATGACGTCAAGTCATCATGCCCCTTATGTCTTGGGCTGCAAACATGCTACAATGGCCGGTACAGTGGGCGTGCGATGCCGTAAGGTGGAGCGAATCCCTAAAAGCCGGTCTCAGTTCGGATTGGGGTCTGCAACTCGACCCCATGAAGGTGGAGTCGCTAGTAATCGCGGATCAGCAACGCCGCGGTGAATACGTTCCCGGGCCTTGTACACACCGCCCGTCACGTCATGAAAGTCGGCAACACCCGAAACTTGCGGCCTAACCCTTCGGGGAGGGAGTGAGTGAAGGTGGGGCTGGCGATTGGGACG

***Nocardiopsis* sp. GSL21-079**

GACGAACGCTGGCGGCGTGCTTAACACATGCAAGTCGAGCGGTAAGGCCCTTCGGGGTACACGAGCGGCGAACGGGTGAGTAACACGTGAGCAACCTGCCCCTGACTCTGGGATAAGCGGTGGAAACGCCGTCTAATACCGGATACGACCCGCCACCTCATGGTGGAGGGTGGAAAGTTTTTCGGTCAGGGATGGGCTCGCGGCCTATCAGCTTGTTGGTGGGGTAACGGCCTACCAAGGCGATTACGGGTAGCCGGCCTGAGAGGGCGACCGGCCACACTGGGACTGAGACACGGCCCAGACTCCTGCGGGAGGCAGCAGTGGGGAATATTGCGCAATGGGCGAAAGCCTGACGCAGCGACGCCGCGTGGGGGATGACGGCCTTCGGGTTGTAAACCTCTTTTACCACCAACGCAGGCTTCCAGTTCTCTGGAGGTTGACGGTAGGTGGGGAATAAGGACCGGCTAACTACGTGCCAGCAGCCGCGGTAATACGTAGGGTCCGAGCGTTGTCCGGAATTATTGGGCGTAAAGAGCTCGTAGGCGGCGTGTCGCGTCTGCTGTGAAAGACCGGGGCTTAACTCCGGTTCTGCAGTGGATACGGGCATGCTAGAGGTAGGTAGGGGAGACTGGAATTCCTGGTGTAGCGGTGAAATGCGCAGATATCAGGAGGAACACCGGTGGCGAAGGCGGGTCTCTGGGCCTTACCTGACGCTGAGGAGCGAAAGCATGGGGAGCGAACAGGATTAGATACCCTGGTAGTCCATGCCGTAAACGTTGGGCGCTAGGTGTGGGGACTTTCCACGGTTTCCGCGCCGTAGCTAACGCATTAAGCGCCCCGCCTGGGGAGTACGGCCGCAAGGCTAAAACTCAAAGGAATTGACGGGGGCCCGCACAAGCGGCGGAGCATGTTGCTTAATTCGACGCAACGCGAAGAACCTTACCAAGGTTTGACATCACCCGTGGACTCGCAGAGATGTGAGGTCATTTAGTTGGCGGGTGACAGGTGGTGCATGGCTGTCGTCAGCTCGTGTCGTGAGATGTTGGGTTAAGTCCCGCAACGAGCGCAACCCTTGTTCCATGTTGCCAGCACGTAATGGTGGGGACTCATGGGAGACTGCCGGGGTCAACTCGGAGGAAGGTGGGGATGACGTCAAGTCATCATGCCCCTTATGTCTTGGGCTGCAAACATGCTACAATGGCCGGTACAATGGGCGTGCGATACCGTAAGGTGGAGCGAATCCCTAAAAGCCGGTCTCAGTTCGGATTGGGGTCTGCAACTCGACCCCATGAAGGTGGAGTCGCTAGTAATCGCGGATCAGCAACGCCGCGGTGAATACGTTCCCGGGCCTTGTACACACCGCCCGTCACGTCATGAAAGTCGGCAACACCCGAAACTTGCGGCCTAACCCCTTGTGGGAGGGAGTGAGTGAAGGTGGGGCTGGCGATTGGGACG

***Nocardiopsis* sp. GSL21-080**

GACGAACGCTGGCGGCGTGCTTAACACATGCAAGTCGAGCGGTAAGGCCCTTCGGGGTACACGAGCGGCGAACGGGTGAGTAACACGTGAGCAACCTGCCCCTGACTCTGGGATAAGCGGTGGAAACGCCGTCTAATACCGGATACGACCCGCCACCTCATGGTGGAGGGTGGAAAGTTTTTCGGTCAGGGATGGGCTCGCGGCCTATCAGCTTGTTGGTGGGGTAACGGCCTACCAAGGCGATTACGGGTAGCCGGCCTGAGAGGGCGACCGGCCACACTGGGACTGAGACACGGCCCAGACTCCTGCGGGAGGCAGCAGTGGGGAATATTGCGCAATGGGCGAAAGCCTGACGCAGCGACGCCGCGTGGGGGATGACGGCCTTCGGGTTGTAAACCTCTTTTACCACCAACGCAGGCTTCCAGTTCTCTGGAGGTTGACGGTAGGTGGGGAATAAGGACCGGCTAACTACGTGCCAGCAGCCGCGGTAATACGTAGGGTCCGAGCGTTGTCCGGAATTATTGGGCGTAAAGAGCTCGTAGGCGGCGTGTCGCGTCTGCTGTGAAAGACCGGGGCTTAACTCCGGTTCTGCAGTGGATACGGGCATGCTAGAGGTAGGTAGGGGAGACTGGAATTCCTGGTGTAGCGGTGAAATGCGCAGATATCAGGAGGAACACCGGTGGCGAAGGCGGGTCTCTGGGCCTTACCTGACGCTGAGGAGCGAAAGCATGGGGAGCGAACAGGATTAGATACCCTGGTAGTCCATGCCGTAAACGTTGGGCGCTAGGTGTGGGGACTTTCCACGGTTTCCGCGCCGTAGCTAACGCATTAAGCGCCCCGCCTGGGGAGTACGGCCGCAAGGCTAAAACTCAAAGGAATTGACGGGGGCCCGCACAAGCGGCGGAGCATGTTGCTTAATTCGACGCAACGCGAAGAACCTTACCAAGGTTTGACATCACCCGTGGACTCGCAGAGATGTGAGGTCATTTAGTTGGCGGGTGACAGGTGGTGCATGGCTGTCGTCAGCTCGTGTCGTGAGATGTTGGGTTAAGTCCCGCAACGAGCGCAACCCTTGTTCCATGTTGCCAGCACGTAATGGTGGGGACTCATGGGAGACTGCCGGGGTCAACTCGGAGGAAGGTGGGGATGACGTCAAGTCATCATGCCCCTTATGTCTTGGGCTGCAAACATGCTACAATGGCCGGTACAATGGGCGTGCGATACCGTAAGGTGGAGCGAATCCCTAAAAGCCGGTCTCAGTTCGGATTGGGGTCTGCAACTCGACCCCATGAAGGTGGAGTCGCTAGTAATCGCGGATCAGCAACGCCGCGGTGAATACGTTCCCGGGCCTTGTACACACCGCCCGTCACGTCATGAAAGTCGGCAACACCCGAAACTTGCGGCCTAACCCCTTGTGGGAGGGAGTGAGTGAAGGTGGGGCTGGCGATTGGGACG

***Nocardiopsis* sp. GSL21-085**

GACGAACGCTGGCGGCGTGCTTAACACATGCAAGTCGAGCGGTAAGGCCCTTCGGGGTACACGAGCGGCGAACGGGTGAGTAACACGTGAGCAACCTGCCCCTGACTCCGGGATAAGCGGTGGAAACGCCGTCTAATACCGGATACGACCCTCCGCCTCATGGTGGGGGGTGGAAAGTTGTGTCGGTCAGGGATGGGCTCGCGGCCTATCAGCTTGTTGGTGGGGTAACGGCCTACCAAGGCGATTACGGGTAGCCGGCCTGAGAGGGCGACCGGCCACACTGGGACTGAGACACGGCCCAGACTCCTGCGGGAGGCAGCAGTGGGGAATATTGCGCAATGGGCGAAAGCCTGACGCAGCGACGCCGCGTGGGGGATGACGGCCTTCGGGTTGTAAACCTCTTTTACCACCAACGCAGGCTCCGGGTGTTCTCGGGGTTGACGGTAGGTGGGGAATAAGGACCGGCTAACTACGTGCCAGCAGCCGCGGTAATACGTAGGGTCCGAGCGTTGTCCGGAATTATTGGGCGTAAAGAGCTCGTAGGCGGCGTGTCGCGTCTGCTGTGAAAGACCGGGGCTTAACTCCGGTTCTGCAGTGGATACGGGCATGCTAGAGGTAGGTAGGGGAGACTGGAATTCCTGGTGTAGCGGTGAAATGCGCAGATATCAGGAGGAACACCGGTGGCGAAGGCGGGTCTCTGGGCCTTACCTGACGCTGAGGAGCGAAAGCATGGGGAGCGAACAGGATTAGATACCCTGGTAGTCCATGCCGTAAACGTTGGGCGCTAGGTGTGGGGACTTTCCACGGTTTCCGCGCCGTAGCTAACGCATTAAGCGCCCCGCCTGGGGAGTACGGCCGCAAGGCTAAAACTCAAAGGAATTGACGGGGGCCCGCACAAGCGGCGGAGCATGTTGCTTAATTCGACGCAACGCGAAGAACCTTACCAAGGTTTGACATCACCCGTGGACCTGCAGAGATGTGGGGTCATTTAGTTGGTGGGTGACAGGTGGTGCATGGCTGTCGTCAGCTCGTGTCGTGAGATGTTGGGTTAAGTCCCGCAACGAGCGCAACCCTTGTTCCATGTTGCCAGCACGTAGTGGTGGGGACTCATGGGAGACTGCCGGGGTCAACTCGGAGGAAGGTGGGGACGACGTCAAGTCATCATGCCCCTTATGTCTTGGGCTGCAAACATGCTACAATGGCCGGTACAATGGGCGTGCGATACCGTGAGGTGGAGCGAATCCCTAAAAGCCGGTCTCAGTTCGGATTGGGGTCTGCAACTCGACCCCATGAAGGTGGAGTCGCTAGTAATCGCGGATCAGCAACGCCGCGGTGAATACGTTCCCGGGCCTTGTACACACCGCCCGTCACGTCATGAAAGTCGGCAACACCCGAAACTTGTGGCCCAACCCTTCGGGGAGGGAGTGAGTGAAGGTGGGGCTGGCGATTGGGACG

***Nocardiopsis* sp. GSL21-089**

GACGAACGCTGGCGGCGTGCTTAACACATGCAAGTCGAGCGGTAAGGCCCTTCGGGGTACACGAGCGGCGAACGGGTGAGTAACACGTGAGCAACCTGCCCCTGACTCCGGGATAAGCGGTGGAAACGCCGTCTAATACCGGATACGACCCTCCGCCTCATGGTGGGGGGTGGAAAGTTGTGTCGGTCAGGGATGGGCTCGCGGCCTATCAGCTTGTTGGTGGGGTAACGGCCTACCAAGGCGATTACGGGTAGCCGGCCTGAGAGGGCGACCGGCCACACTGGGACTGAGACACGGCCCAGACTCCTGCGGGAGGCAGCAGTGGGGAATATTGCGCAATGGGCGAAAGCCTGACGCAGCGACGCCGCGTGGGGGATGACGGCCTTCGGGTTGTAAACCTCTTTTACCACCAACGCAGGCTCCGGGTGTTCTCGGGGTTGACGGTAGGTGGGGAATAAGGACCGGCTAACTACGTGCCAGCAGCCGCGGTAATACGTAGGGTCCGAGCGTTGTCCGGAATTATTGGGCGTAAAGAGCTCGTAGGCGGCGTGTCGCGTCTGCTGTGAAAGACCGGGGCTTAACTCCGGTTCTGCAGTGGATACGGGCATGCTAGAGGTAGGTAGGGGAGACTGGAATTCCTGGTGTAGCGGTGAAATGCGCAGATATCAGGAGGAACACCGGTGGCGAAGGCGGGTCTCTGGGCCTTACCTGACGCTGAGGAGCGAAAGCATGGGGAGCGAACAGGATTAGATACCCTGGTAGTCCATGCCGTAAACGTTGGGCGCTAGGTGTGGGGACTTTCCACGGTTTCCGCGCCGTAGCTAACGCATTAAGCGCCCCGCCTGGGGAGTACGGCCGCAAGGCTAAAACTCAAAGGAATTGACGGGGGCCCGCACAAGCGGCGGAGCATGTTGCTTAATTCGACGCAACGCGAAGAACCTTACCAAGGTTTGACATCACCCGTGGACCTGCAGAGATGTGGGGTCATTTAGTTGGTGGGTGACAGGTGGTGCATGGCTGTCGTCAGCTCGTGTCGTGAGATGTTGGGTTAAGTCCCGCAACGAGCGCAACCCTTGTTCCATGTTGCCAGCACGTAGTGGTGGGGACTCATGGGAGACTGCCGGGGTCAACTCGGAGGAAGGTGGGGACGACGTCAAGTCATCATGCCCCTTATGTCTTGGGCTGCAAACATGCTACAATGGCCGGTACAATGGGCGTGCGATACCGTGAGGTGGAGCGAATCCCTAAAAGCCGGTCTCAGTTCGGATTGGGGTCTGCAACTCGACCCCATGAAGGTGGAGTCGCTAGTAATCGCGGATCAGCAACGCCGCGGTGAATACGTTCCCGGGCCTTGTACACACCGCCCGTCACGTCATGAAAGTCGGCAACACCCGAAACTTGTGGCCCAACCCTTCGGGGAGGGAGTGAGTGAAGGTGGGGCTGGCGATTGGGACG

***Nocardiopsis* sp. GSL21-156**

GACGAACGCTGGCGGCGTGCTTAACACATGCAAGTCGAGCGGTAAGGCCCTTCGGGGTACACGAGCGGCGAACGGGTGAGTAACACGTGAGCAACCTGCCCCCGACTCCGGGATAAGCGGTGGAAACGCCGTCTAATACCGGATACGACCGGCCACCTCATGGCGGCCGGTGGAAAGTTTTCTCGGTTGGGGATGGGCTCGCGGCCTATCAGCTAGTTGGTGGGGTAAAGGCCTACCAAGGCGATTACGGGTAGCCGGCCTGAGAGGGCGACCGGCCACACTGGGACTGAGACACGGCCCAGACTCCTGCGGGAGGCAGCAGTGGGGAATATTGCGCAATGGGCGAAAGCCTGACGCAGCGACGCCGCGTGGGGGATGACGGCCTTCGGGTTGTAAACCTCTTTTACCACCAACGCAGGCTCCCAGTTCTCTGGGGGTTGACGGTAGGTGGGGAATAAGGACCGGCTAACTACGTGCCAGCAGCCGCGGTAATACGTAGGGTCCGAGCGTTGTCCGGAATTATTGGGCGTAAAGAGCTCGTAGGCGGCGTGTCGCGTCTGCTGTGAAAGACCGGGGCTTAACTCCGGTTCTGCAGTGGATACGGGCATGCTAGAGGTAGGTAGGGGAGACTGGAATTCCTGGTGTAGCGGTGAAATGCGCAGATATCAGGAGGAACACCGGTGGCGAAGGCGGGTCTCTGGGCCTTACCTGACGCTGAGGAGCGAAAGCATGGGGAGCGAACAGGATTAGATACCCTGGTAGTCCATGCCGTAAACGTTGGGCGCTAGGTGTGGGGACTTTCCACGGTTTCCGCGCCGTAGCTAACGCATTAAGCGCCCCGCCTGGGGAGTACGGCCGCAAGGCTAAAACTCAAAGGAATTGACGGGGGCCCGCACAAGCGGCGGAGCATGTTGCTTAATTCGACGCAACGCGAAGAACCTTACCAAGGTTTGACATCACCCGTGGACCTGCAGAGATGTGGGGTCATTTAGTTGGTGGGTGACAGGTGGTGCATGGCTGTCGTCAGCTCGTGTCGTGAGATGTTGGGTTAAGTCCCGCAACGAGCGCAACCCTTATTCCATGTTGCCAGCACGTAATGGTGGGGACTCATGGGAGACTGCCGGGGTCAACTCGGAGGAAGGTGGGGACGACGTCAAGTCATCATGCCCCTTATGTCTTGGGCTGCAAACATGCTACAATGGCCGGTACAATGGGCGTGCGATACCGTAAGGTGGAGCGAATCCCTAAAAGCCGGTCTCAGTTCGGATTGGGGTCTGCAACTCGACCCCATGAAGGTGGAGTCGCTAGTAATCGCGGATCAGCAACGCCGCGGTGAATACGTTCCCGGGCCTTGTACACACCGCCCGTCACGTCATGAAAGTCGGCAACACCCGAAACTTGTGGCCTAACCCTTCGGGGAGGGAATGAGTGAAGGTGGGGCTGGCGATTGGGACG

***Nocardiopsis* sp. GSL23-180**

GACGAACGCTGGCGGCGTGCTTAACACATGCAAGTCGAGCGGTAAGGCCCTTCGGGGTACACGAGCGGCGAACGGGTGAGTAACACGTGAGCAACCTGCCCCCGACTCCGGGATAAGCGGCGGAAACGCCGTCTAATACCGGATACGACCCACCGCCTCATGGCGGAGGGTGGAAAGTTTTTCGGTTGGGGATGGGCTCGCGGCCTATCAGCTTGTTGGTGGGGTAAAGGCCTACCAAGGCGATTACGGGTAGCCGGCCTGAGAGGGCGACCGGCCACACTGGGACTGAGACACGGCCCAGACTCCTACGGGAGGCAGCAGTGGGGAATATTGCGCAATGGGCGAAAGCCTGACGCAGCGACGCCGCGTGGGGGATGACGGCCTTCGGGTTGTAAACCTCTTTTACCACCAACGCAGGCTCCCAGTACTCTGGGGGTTGACGGTAGGTGGGGAATAAGGACCGGCTAACTACGTGCCAGCAGCCGCGGTAATACGTAGGGTCCGAGCGTTGTCCGGAATTATTGGGCGTAAAGAGCTCGTAGGCGGCGTGTCGCGTCTGCTGTGAAAGACCGGGGCTTAACCCCGGTTCTGCAGTGGATACGGGCATGCTAGAGGTAGGTAGGGGAGACTGGAATTCCTGGTGTAGCGGTGAAATGCGCAGATATCAGGAGGAACACCGGTGGCGAAGGCGGGTCTCTGGGCCTTACCTGACGCTGAGGAGCGAAAGCATGGGGAGCGAACAGGATTAGATACCCTGGTAGTCCATGCCGTAAACGTTGGGCGCTAGGTGTGGGGACTTTCCACGGTTTCCGCGCCGCAGCTAACGCATTAAGCGCCCCGCCTGGGGAGTACGGCCGCAAGGCTAAAACTCAAAGGAATTGACGGGGGCCCGCACAAGCGGCGGAGCATGTTGCTTAATTCGACGCAACGCGAAGAACCTTACCAAGGTTTGACATCGCCCGTGGACCTGTAGAGATACAGGGTCATTTAGTTGGTGGGTGACAGGTGGTGCATGGCTGTCGTCAGCTCGTGTCGTGAGATGTTGGGTTAAGTCCCGCAACGAGCGCAACCCCTATTCTATGTTGCCAGCACGTAATGGTGGGGACTCATAGGAGACTGCCGGGGTCAACTCGGAGGAAGGTGGGGACGACGTCAAGTCATCATGCCCCTTATGTCTTGGGCTGCAAACATGCTACAATGGCCGGTACAATGGGCGTGCGAGACCGCAAGGTGGAGCGAATCCCTAAAAGCCGGTCTCAGTTCGGATTGGGGTCTGCAACTCGACCCCATGAAGGTGGAGTCGCTAGTAATCGCGGATCAGCAACGCCGCGGTGAATACGTTCCCGGGCCTTGTACACACCGCCCGTCACGTCATGAAAGTCGGCAACACCCGAAACTTGTGGCCTAACCCTTCGGGGAGGGAATGAGTGAAGGTGGGGCTGGCGATTGGGACG

***Nocardiopsis* sp. GSL23-181**

CGAACGCTGGCGGCGTGCTTAACACATGCAAGTCGAGCGGTAAGGCCCTTCGGGGTACACGAGCGGCGAACGGGTGAGTAACACGTGAGCAACCTGCCCCTGACTCCGGGATAAGCGGTGGAAACGCCGTCTAATACCGGATATGACACACCATCTCCTGGTGGTGTGTGGAAAGTTTTGTCGGTCAGGGATGGGCTCGCGGCCTATCAGCTAGTTGGTGGGGTAAAGGCCTACCAAGGCGATTACGGGTAGCCGGCCTGAGAGGGCGACCGGCCACACTGGGACTGAGACACGGCCCAGACTCCTGCGGGAGGCAGCAGTGGGGAATATTGCACAATGGGCGAAAGCCTGATGCAGCGACGCCGCGTGGGGGATGACGGCCTTCGGGTTGTAAACCTCTTTTACCACCAACGCAGGCCCCGGGTTCTCTCGGGGTTGACGGTAGGTGGGGAATAAGGACCGGCTAACTACGTGCCAGCAGCCGCGGTAATACGTAGGGTCCGAGCGTTGTCCGGAATTATTGGGCGTAAAGAGCTCGTAGGCGGCGTGTCGCGTCTGCTGTGAAAGACCGGGGCTTAACCCCGGTTCTGCAGTGGATACGGGCATGCTAGAGGTAGGTAGGGGAGACTGGAATTCCTGGTGTAGCGGTGAAATGCGCAGATATCAGGAGGAACACCGGTGGCGAAGGCGGGTCTCTGGGCCTTACCTGACGCTGAGGAGCGAAAGCATGGGGAGCGAACAGGATTAGATACCCTGGTAGTCCATGCCGTAAACGTTGGGCGCTAGGTGTGGGGACTTTCCACGGTTTCCGCGCCGTAGCTAACGCATTAAGCGCCCCGCCTGGGGAGTACGGCCGCAAGGCTAAAACTCAAAGGAATTGACGGGGGCCCGCACAAGCGGCGGAGCATGTTGCTTAATTCGACGCAACGCGAAGAACCTTACCAAGGTTTGACATCACCCGTGGACCTGTAGAGATACAGGGTCATTTAGTTGGTGGGTGACAGGTGGTGCATGGCTGTCGTCAGCTCGTGTCGTGAGATGTTGGGTTAAGTCCCGCAACGAGCGCAACCCTTGTTCCATGTTGCCAGCACGTAATGGTGGGGACTCATGGGAGACTGCCGGGGTCAACTCGGAGGAAGGTGGGGACGACGTCAAGTCATCATGCCCCTTATGTCTTGGGCTGCAAACATGCTACAATGGCCGGTACAATGGGCGTGCGATACCGTAAGGTGGAGCGAATCCCTTAAAGCCGGTCTCAGTTCGGATTGGGGTCTGCAACTCGACCCCATGAAGGTGGAGTCGCTAGTAATCGCGGATCAGCAACGCCGCGGTGAATACGTTCCCGGGCCTTGTACACACCGCCCGTCACGTCATGAAAGTCGGCAACACCCGAAACTTGTGGCCTAACCCTTCGGGGAGGGAATGAGTGAAGGTGGGGCTGGCGATTGGGA

***Nocardiopsis* sp. GSL23-183**

GACGAACGCTGGCGGCGTGCTTAACACATGCAAGTCGAGCGGTAAGGCCCTTCGGGGTACACGAGCGGCGAACGGGTGAGTAACACGTGAGCAACCTGCCCCCGACTCCGGGATAAGCGGCGGAAACGCCGTCTAATACCGGATACGACCCACCGCCTCATGGCGGAGGGTGGAAAGTTTTTCGGTTGGGGATGGGCTCGCGGCCTATCAGCTTGTTGGTGGGGTAAAGGCCTACCAAGGCGATTACGGGTAGCCGGCCTGAGAGGGCGACCGGCCACACTGGGACTGAGACACGGCCCAGACTCCTACGGGAGGCAGCAGTGGGGAATATTGCGCAATGGGCGAAAGCCTGACGCAGCGACGCCGCGTGGGGGATGACGGCCTTCGGGTTGTAAACCTCTTTTACCACCAACGCAGGCTCCCAGTACTCTGGGGGTTGACGGTAGGTGGGGAATAAGGACCGGCTAACTACGTGCCAGCAGCCGCGGTAATACGTAGGGTCCGAGCGTTGTCCGGAATTATTGGGCGTAAAGAGCTCGTAGGCGGCGTGTCGCGTCTGCTGTGAAAGACCGGGGCTTAACCCCGGTTCTGCAGTGGATACGGGCATGCTAGAGGTAGGTAGGGGAGACTGGAATTCCTGGTGTAGCGGTGAAATGCGCAGATATCAGGAGGAACACCGGTGGCGAAGGCGGGTCTCTGGGCCTTACCTGACGCTGAGGAGCGAAAGCATGGGGAGCGAACAGGATTAGATACCCTGGTAGTCCATGCCGTAAACGTTGGGCGCTAGGTGTGGGGACTTTCCACGGTTTCCGCGCCGCAGCTAACGCATTAAGCGCCCCGCCTGGGGAGTACGGCCGCAAGGCTAAAACTCAAAGGAATTGACGGGGGCCCGCACAAGCGGCGGAGCATGTTGCTTAATTCGACGCAACGCGAAGAACCTTACCAAGGTTTGACATCGCCCGTGGACCTGTAGAGATACAGGGTCATTTAGTTGGTGGGTGACAGGTGGTGCATGGCTGTCGTCAGCTCGTGTCGTGAGATGTTGGGTTAAGTCCCGCAACGAGCGCAACCCCTATTCTATGTTGCCAGCACGTAATGGTGGGGACTCATAGGAGACTGCCGGGGTCAACTCGGAGGAAGGTGGGGACGACGTCAAGTCATCATGCCCCTTATGTCTTGGGCTGCAAACATGCTACAATGGCCGGTACAATGGGCGTGCGAGACCGCAAGGTGGAGCGAATCCCTAAAAGCCGGTCTCAGTTCGGATTGGGGTCTGCAACTCGACCCCATGAAGGTGGAGTCGCTAGTAATCGCGGATCAGCAACGCCGCGGTGAATACGTTCCCGGGCCTTGTACACACCGCCCGTCACGTCATGAAAGTCGGCAACACCCGAAACTTGTGGCCTAACCCTTCGGGGAGGGAATGAGTGAAGGTGGGGCTGGCGATTGGGACG

***Nocardiopsis* sp. GSL23-184**

GACGAACGCTGGCGGCGTGCTTAACACATGCAAGTCGAGCGGTAAGGCCCTTCGGGGTACACGAGCGGCGAACGGGTGAGTAACACGTGAGCAACCTGCCCCCGACTCCGGGATAAGCGGTGGAAACGCCGTCTAATACCGGATACGACCGGCCACCTCATGGCGGCCGGTGGAAAGTTTTCTCGGTTGGGGATGGGCTCGCGGCCTATCAGCTAGTTGGTGGGGTAAAGGCCTACCAAGGCGATTACGGGTAGCCGGCCTGAGAGGGCGACCGGCCACACTGGGACTGAGACACGGCCCAGACTCCTGCGGGAGGCAGCAGTGGGGAATATTGCGCAATGGGCGAAAGCCTGACGCAGCGACGCCGCGTGGGGGATGACGGCCTTCGGGTTGTAAACCTCTTTTACCACCAACGCAGGCTCCCAGTTCTCTGGGGGTTGACGGTAGGTGGGGAATAAGGACCGGCTAACTACGTGCCAGCAGCCGCGGTAATACGTAGGGTCCGAGCGTTGTCCGGAATTATTGGGCGTAAAGAGCTCGTAGGCGGCGTGTCGCGTCTGCTGTGAAAGACCGGGGCTTAACTCCGGTTCTGCAGTGGATACGGGCATGCTAGAGGTAGGTAGGGGAGACTGGAATTCCTGGTGTAGCGGTGAAATGCGCAGATATCAGGAGGAACACCGGTGGCGAAGGCGGGTCTCTGGGCCTTACCTGACGCTGAGGAGCGAAAGCATGGGGAGCGAACAGGATTAGATACCCTGGTAGTCCATGCCGTAAACGTTGGGCGCTAGGTGTGGGGACTTTCCACGGTTTCCGCGCCGTAGCTAACGCATTAAGCGCCCCGCCTGGGGAGTACGGCCGCAAGGCTAAAACTCAAAGGAATTGACGGGGGCCCGCACAAGCGGCGGAGCATGTTGCTTAATTCGACGCAACGCGAAGAACCTTACCAAGGTTTGACATCACCCGTGGACCTGCAGAGATGTGGGGTCATTTAGTTGGTGGGTGACAGGTGGTGCATGGCTGTCGTCAGCTCGTGTCGTGAGATGTTGGGTTAAGTCCCGCAACGAGCGCAACCCTTATTCCATGTTGCCAGCACGTAATGGTGGGGACTCATGGGAGACTGCCGGGGTCAACTCGGAGGAAGGTGGGGACGACGTCAAGTCATCATGCCCCTTATGTCTTGGGCTGCAAACATGCTACAATGGCCGGTACAATGGGCGTGCGATACCGTAAGGTGGAGCGAATCCCTAAAAGCCGGTCTCAGTTCGGATTGGGGTCTGCAACTCGACCCCATGAAGGTGGAGTCGCTAGTAATCGCGGATCAGCAACGCCGCGGTGAATACGTTCCCGGGCCTTGTACACACCGCCCGTCACGTCATGAAAGTCGGCAACACCCGAAACTTGTGGCCTAACCCTTCGGGGAGGGAATGAGTGAAGGTGGGGCTGGCGATTGGGACG

***Nocardiopsis* sp. GSL23-185**

GACGAACGCTGGCGGCGTGCTTAACACATGCAAGTCGAGCGGTAAGGCCCTTCGGGGTACACGAGCGGCGAACGGGTGAGTAACACGTGAGCAACCTGCCCCCGACTCCGGGATAAGCGGTGGAAACGCCGTCTAATACCGGATACGACCGGCCACCTCATGGCGGCCGGTGGAAAGTTTTCTCGGTTGGGGATGGGCTCGCGGCCTATCAGCTAGTTGGTGGGGTAAAGGCCTACCAAGGCGATTACGGGTAGCCGGCCTGAGAGGGCGACCGGCCACACTGGGACTGAGACACGGCCCAGACTCCTGCGGGAGGCAGCAGTGGGGAATATTGCGCAATGGGCGAAAGCCTGACGCAGCGACGCCGCGTGGGGGATGACGGCCTTCGGGTTGTAAACCTCTTTTACCACCAACGCAGGCTCCCAGTTCTCTGGGGGTTGACGGTAGGTGGGGAATAAGGACCGGCTAACTACGTGCCAGCAGCCGCGGTAATACGTAGGGTCCGAGCGTTGTCCGGAATTATTGGGCGTAAAGAGCTCGTAGGCGGCGTGTCGCGTCTGCTGTGAAAGACCGGGGCTTAACTCCGGTTCTGCAGTGGATACGGGCATGCTAGAGGTAGGTAGGGGAGACTGGAATTCCTGGTGTAGCGGTGAAATGCGCAGATATCAGGAGGAACACCGGTGGCGAAGGCGGGTCTCTGGGCCTTACCTGACGCTGAGGAGCGAAAGCATGGGGAGCGAACAGGATTAGATACCCTGGTAGTCCATGCCGTAAACGTTGGGCGCTAGGTGTGGGGACTTTCCACGGTTTCCGCGCCGTAGCTAACGCATTAAGCGCCCCGCCTGGGGAGTACGGCCGCAAGGCTAAAACTCAAAGGAATTGACGGGGGCCCGCACAAGCGGCGGAGCATGTTGCTTAATTCGACGCAACGCGAAGAACCTTACCAAGGTTTGACATCACCCGTGGACCTGCAGAGATGTGGGGTCATTTAGTTGGTGGGTGACAGGTGGTGCATGGCTGTCGTCAGCTCGTGTCGTGAGATGTTGGGTTAAGTCCCGCAACGAGCGCAACCCTTATTCCATGTTGCCAGCACGTAATGGTGGGGACTCATGGGAGACTGCCGGGGTCAACTCGGAGGAAGGTGGGGACGACGTCAAGTCATCATGCCCCTTATGTCTTGGGCTGCAAACATGCTACAATGGCCGGTACAATGGGCGTGCGATACCGTAAGGTGGAGCGAATCCCTAAAAGCCGGTCTCAGTTCGGATTGGGGTCTGCAACTCGACCCCATGAAGGTGGAGTCGCTAGTAATCGCGGATCAGCAACGCCGCGGTGAATACGTTCCCGGGCCTTGTACACACCGCCCGTCACGTCATGAAAGTCGGCAACACCCGAAACTTGTGGCCTAACCCTTCGGGGAGGGAATGAGTGAAGGTGGGGCTGGCGATTGGGACG

***Nocardiopsis* sp. GSL24-012**

GACGAACGCTGGCGGCGTGCTTAACACATGCAAGTCGAGCGGTAAGGCCCTTCGGGGTACACGAGCGGCGAACGGGTGAGTAACACGTGAGCAACCTGCCCCTGACTCCGGGATAAGCGGTGGAAACGCCGTCTAATACCGGATACGACCCAAGGCCTCCTGGCCATGGGTGGAAAGTTTTTCGGTCGGGGATGGGCTCGCGGCCTATCAGCTTGTTGGTGGGGTAACAGCCTACCAAGGCGATTACGGGTAGCCGGCCTGAGAGGGCGACCGGCCACACTGGGACTGAGACACGGCCCAGACTCCTACGGGAGGCAGCAGTGGGGAATATTGCGCAATGGGCGAAAGCCTGACGCAGCGACGCCGCGTGGGGGATGACGGCCTTCGGGTTGTAAACCTCTTTTACCACCAACGCAGGCTCCCAGTTCTCTGGGGGTTGACGGTAGGTGGGGAATAAGGACCGGCTAACTACGTGCCAGCAGCCGCGGTAATACGTAGGGTCCGAGCGTTGTCCGGAATTATTGGGCGTAAAGAGCTCGTAGGCGGCGTGTCGCGTCTGCTGTGAAAGACCGGGGCTTAACCCCGGTTCTGCAGTGGATACGGGCATGCTAGAGGTAGGTAGGGGAGACTGGAATTCCTGGTGTAGCGGTGAAATGCGCAGATATCAGGAGGAACACCGGTGGCGAAGGCGGGTCTCTGGGCCTTACCTGACGCTGAGGAGCGAAAGCATGGGGAGCGAACAGGATTAGATACCCTGGTAGTCCATGCCGTAAACGTTGGGCGCTAGGTGTGGGGACTTTCCACGGTTTCCGCGCCGTAGCTAACGCATTAAGCGCCCCGCCTGGGGAGTACGGCCGCAAGGCTAAAACTCAAAGGAATTGACGGGGGCCCGCACAAGCGGCGGAGCATGTTGCTTAATTCGACGCAACGCGAAGAACCTTACCAAGGTTTGACATCACCCGTGGACCTGCAGAGATGTGGGGTCATTTAGTTGGCGGGTGACAGGTGGTGCATGGCTGTCGTCAGCTCGTGTCGTGAGATGTTGGGTTAAGTCCCGCAACGAGCGCAACCCTTATTCCATGTTGCCAGCACGTAATGGTGGGGACTCATGGGAGACTGCCGGGGTCAACTCGGAGGAAGGTGGGGATGACGTCAAGTCATCATGCCCCTTATGTCTTGGGCTGCAAACATGCTACAATGGCCGGTACAATGGGCGTGCGATGCCGCAAGGTGGAGCGAATCCCTAAAAGCCGGTCTCAGTTCGGATTGGGGTCTGCAACTCGACCCCATGAAGGTGGAGTCGCTAGTAATCGCGGATCAGCAACGCCGCGGTGAATACGTTCCCGGGCCTTGTACACACCGCCCGTCACGTCATGAAAGTCGGCAACACCCGAAACTTGCGGCCTAACCCTTCGGGGAGGGAGTGAGTGAAGGTGGGGCTGGCGATTGGGACG

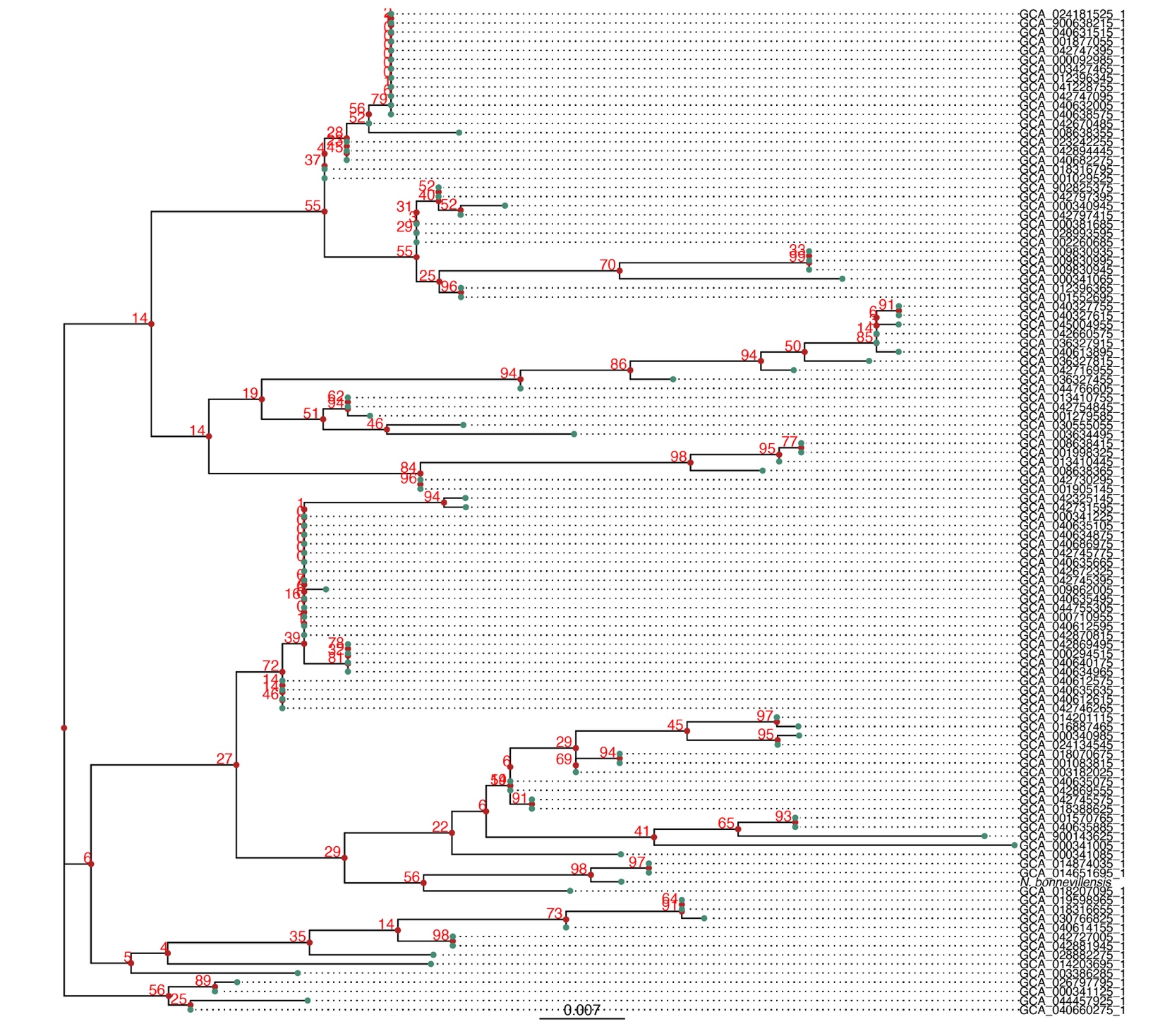

**Supplementary Figure 1**: Maximum-likelihood phylogenetic tree of 16S rRNA gene sequences from *Nocardiopsis* genomes exhibiting ≥ 82% ANI to *N. bonnevillensis* (n= 110 genomes). The tree was constructed using RAxML^1^ with 1000 bootstrap replicates, and bootstrap support values are shown at the nodes. *N. bonnevillensis* is highlighted with a blue arrow to indicate its phylogenetic placement within the genus.

**Supplementary Result 1:** *N. bonnevillensis* growth across NaCl, temperature, and pH ranges

To enable direct comparison with previously described *Nocardiopsis* species, we report salt tolerance in terms of NaCl concentration rather than overall salinity, as this is sometimes the standard format used in other taxonomic descriptions. Growth was observed from 0–16% NaCl (optimal 2–9%),10–45°C (optimal 25–37°C), and pH 5–11 (optimal pH 7–11) (Supplementary Fig. 2). Tolerated salinity ranges varied at temperature extremes. *N. bonnevillensis* exhibited poor growth on ISP2 and Potato Dextrose agar but grew well on ISP3–5 and Czapek-Dox agar. Diffuse yellow pigmentation was observed on ISP4 containing 8% NaCl and on YPM supplemented with 0–4% NaCl, with variation depending on incubation temperature (Supplementary Table 1).

MALDI-based metabolite profiling across a range of fermentation conditions revealed a largely conserved small-molecule profile, as shown by the resulting Metabolite Association Network (MAN). However, distinct small molecules were detected under conditions where growth was suboptimal (Supplementary Fig. 3). These conditions-specific features will guide efforts to identify and isolate novel small molecules.

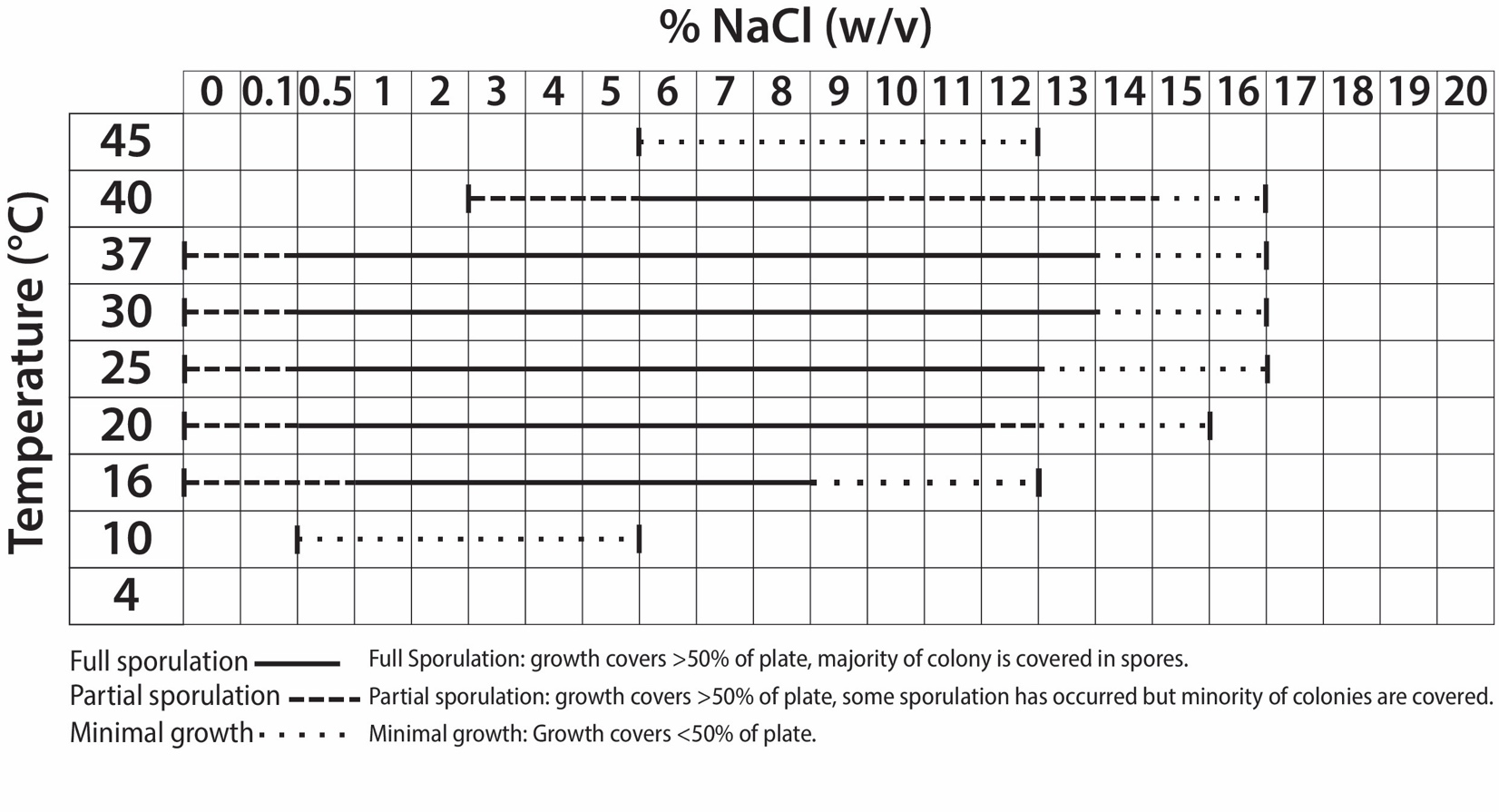

**Supplementary Figure 2**: Growth of *N. bonnevillensis* across temperature and NaCl ranges.

Growth was assessed on YPM agar supplemented with 0–16% NaCl (w/v) and incubated for 14 days at temperatures ranging from 4–45 °C.

**Supplementary Table 1**: Quality and color observations for *N. bonnevillensis* grown on standard media and YPM agar^2^. Substrate and mycelia were assessed after 14 days of growth. Color assignments follow Ridgway, R (1912)^3^ .

| **Media** | **Growth** | **Substrate mycelia** | **Aerial mycelia** | **Soluble Pigment** |
| --- | --- | --- | --- | --- |
| ISP2 | Poor | light yellowish olive | white | - |
| ISP3 | Good | light yellowish olive | white | citron yellow |
| ISP4 | Good | clay color | white to pale drab-gray | citron yellow |
| ISP5 | Good | Saccardo's umber | cream color | - |
| Czapek-Dox | moderate | light yellowish olive | white to pale drab-gray | - |
| Potato Dextrose | Poor | light yellowish olive | white | - |
| YPM | Good | clay color | white to cream buff | citron yellow* |

*Soluble pigment was observed at 16 °C (0–4% NaCl), 20 °C (0–3% NaCl), 25 °C (0–2% NaCl), and 30 °C (0–1% NaCl).

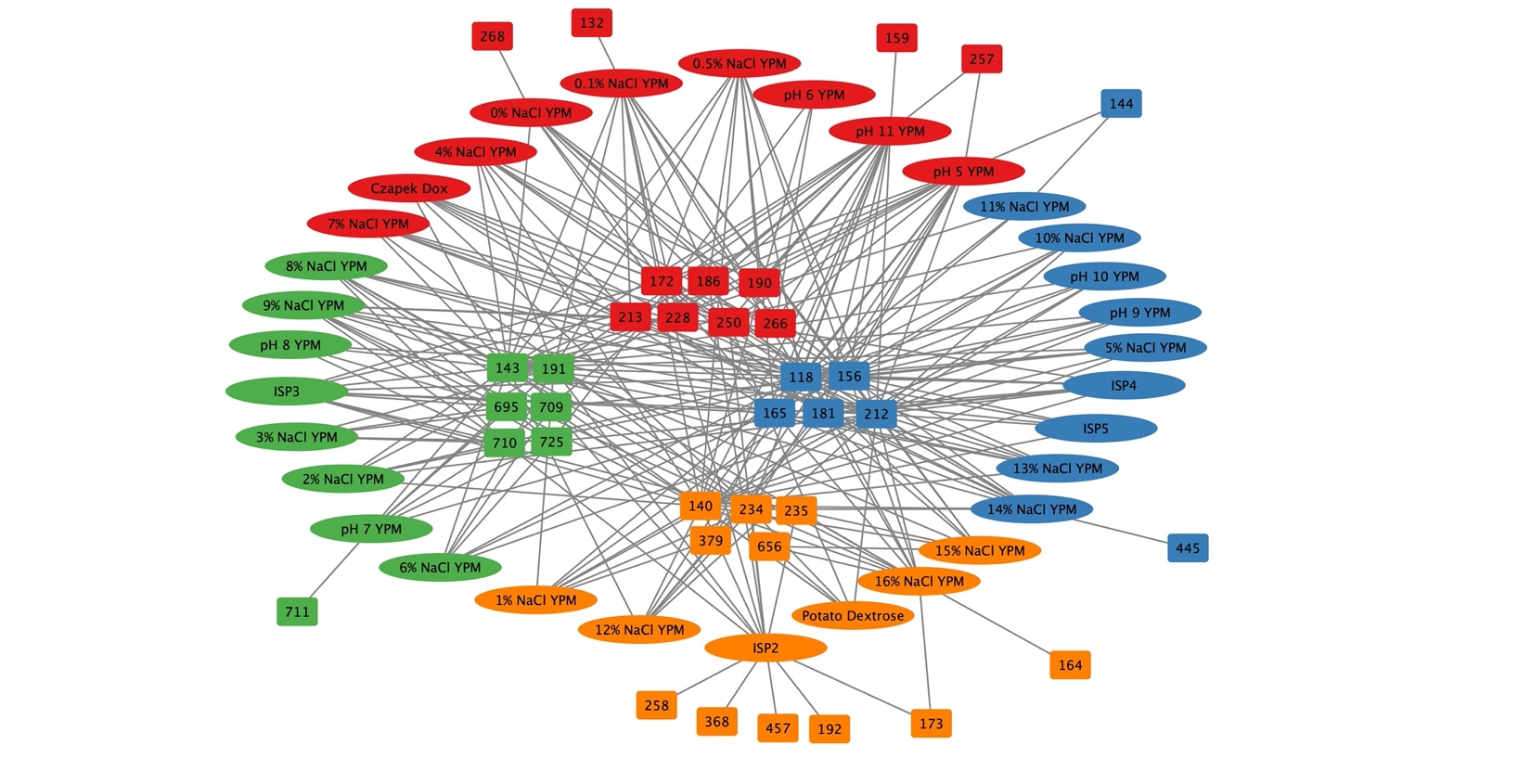

**Supplementary Figure 3**: Metabolic Association Network (MAN) of condition-dependent small molecule production by *N. bonnevillensis.* A Metabolic Association Network (MAN) generated using the IDBac web analysis platform^4^. Relative intensity threshold = 0.05, relative frequency threshold = 1.00, m/z tolerance = 0.10. Elliptical nodes represent media conditions, rectangular nodes represent small molecules (m/z). Node colors are based on network community detection. Edges between small molecules and media condition nodes indicate where the m/z was detected in all three replicates for that fermentation condition. Interior small-molecule nodes were detected in ≥ 3 media conditions, whereas exterior nodes were detected in 1–2 media conditions. All media were supplemented with 8% NaCl (w/v) unless otherwise specified. Media control spectra were subtracted prior to MAN assembly.

**Supplementary Result 2:** Catalase and oxidase activity of *N. bonnevillensis*

Catalase activity was assessed by transferring a colony of *N. bonnevillensis* grown for 18 hours at 30 °C on YPM agar supplemented with 5% salinity (Instant Ocean, w/v) into a sterile 5 mL glass test tube containing 2 mL of 2% hydrogen peroxide. Immediate effervescence was observed, indicating that *N. bonnevillensis* is catalase-positive.

Oxidase activity was evaluated using a separate colony grown under the same conditions, which was transferred using a sterile toothpick to filter paper impregnated with a 1% solution of *N*,*N*,*N*’,*N*’-tetramethyl-*p*-phenylenediamine dihydrochloride (TMPD). The colony turned blue within 10–15 seconds, demonstrating that *N. bonnevillensis* is positive for oxidase activity.

**Supplementary Result 3:** Biosynthetic gene cluster identification by genome mining. Genome mining of the *N. bonnevillensis* genome using antiSMASH^5^ revealed 19 putative biosynthetic gene clusters (BGCs) (Extended Fig. 2a), none of which were located at contig edges, suggesting they were likely annotated in their entirety. The majority of the 19 BGCs showed ≤ 85% similarity to characterized clusters in the MiBIG v.3 database^6^, classifying them as “uncharacterized” (Supplemental Table 2) and indicative of substantial unexplored biosynthetic diversity. The predicted BGCs encompassed diverse classes, including type I and type II polyketide synthases (PKSs), nonribosomal peptide synthetases (NRPSs), hybrid PKS-NRPS systems, terpenes, siderophores, and ribosomally synthesized and post-translationally modified peptides (RiPPs). The absence of high-similarity matches to known clusters suggests that *N. bonnevillensis* may encode novel metabolic pathways and potentially produce structurally distinct metabolites or natural products not yet observed.

**Supplementary Table 2**: antiSMASH^5^ summary of biosynthetic gene clusters identified in *N. bonnevillensis,*

| Region | Type | From | To | Similarity Confidence | Most similar known cluster |
| --- | --- | --- | --- | --- | --- |
| [Region 4.1](https://word-edit.officeapps.live.com/we/wordeditorframe.aspx?ui=en-US&rs=en-US&IsLicensedUser=1&WOPISrc=https%3A%2F%2Fapi.box.com%2Fwopi%2Ffiles%2F1977927011490#RANGE!r4c1) ^*^ | terpene,lassopeptide | 444,759 | 488,322 |  |  |
| [Region 5.1](https://word-edit.officeapps.live.com/we/wordeditorframe.aspx?ui=en-US&rs=en-US&IsLicensedUser=1&WOPISrc=https%3A%2F%2Fapi.box.com%2Fwopi%2Ffiles%2F1977927011490#RANGE!r5c1) | [ectoine](https://docs.antismash.secondarymetabolites.org/glossary/#ectoine) | 54,093 | 64,494 | Medium | [ectoine](https://mibig.secondarymetabolites.org/go/BGC0002052.3) |
| [Region 5.2](https://word-edit.officeapps.live.com/we/wordeditorframe.aspx?ui=en-US&rs=en-US&IsLicensedUser=1&WOPISrc=https%3A%2F%2Fapi.box.com%2Fwopi%2Ffiles%2F1977927011490#RANGE!r5c2) | [butyrolactone](https://docs.antismash.secondarymetabolites.org/glossary/#butyrolactone) | 546,142 | 556,924 |  |  |
| [Region 5.3](https://word-edit.officeapps.live.com/we/wordeditorframe.aspx?ui=en-US&rs=en-US&IsLicensedUser=1&WOPISrc=https%3A%2F%2Fapi.box.com%2Fwopi%2Ffiles%2F1977927011490#RANGE!r5c3) | [T1PKS](https://docs.antismash.secondarymetabolites.org/glossary/#t1pks) | 575,707 | 631,726 |  |  |
| [Region 6.1](https://word-edit.officeapps.live.com/we/wordeditorframe.aspx?ui=en-US&rs=en-US&IsLicensedUser=1&WOPISrc=https%3A%2F%2Fapi.box.com%2Fwopi%2Ffiles%2F1977927011490#RANGE!r6c1) | [RiPP-like](https://docs.antismash.secondarymetabolites.org/glossary/#ripp-like) | 488,568 | 499,494 |  |  |
| [Region 6.2](https://word-edit.officeapps.live.com/we/wordeditorframe.aspx?ui=en-US&rs=en-US&IsLicensedUser=1&WOPISrc=https%3A%2F%2Fapi.box.com%2Fwopi%2Ffiles%2F1977927011490#RANGE!r6c2) | [azole-containing-RiPP](https://docs.antismash.secondarymetabolites.org/glossary/#azole-containing-ripp) | 1,067,428 | 1,095,366 |  |  |
| [Region 6.3](https://word-edit.officeapps.live.com/we/wordeditorframe.aspx?ui=en-US&rs=en-US&IsLicensedUser=1&WOPISrc=https%3A%2F%2Fapi.box.com%2Fwopi%2Ffiles%2F1977927011490#RANGE!r6c3) ^*^ | NRPS,lanthipeptide-class-ii | 1,099,998 | 1,169,451 |  |  |
| [Region 6.4](https://word-edit.officeapps.live.com/we/wordeditorframe.aspx?ui=en-US&rs=en-US&IsLicensedUser=1&WOPISrc=https%3A%2F%2Fapi.box.com%2Fwopi%2Ffiles%2F1977927011490#RANGE!r6c4) | [NRPS](https://docs.antismash.secondarymetabolites.org/glossary/#nrps) | 1,335,684 | 1,400,513 |  |  |
| [Region 6.5](https://word-edit.officeapps.live.com/we/wordeditorframe.aspx?ui=en-US&rs=en-US&IsLicensedUser=1&WOPISrc=https%3A%2F%2Fapi.box.com%2Fwopi%2Ffiles%2F1977927011490#RANGE!r6c5) | [terpene](https://docs.antismash.secondarymetabolites.org/glossary/#terpene) | 2,135,074 | 2,163,338 | High | [isorenieratene](https://mibig.secondarymetabolites.org/go/BGC0001456.3) |
| [Region 6.6](https://word-edit.officeapps.live.com/we/wordeditorframe.aspx?ui=en-US&rs=en-US&IsLicensedUser=1&WOPISrc=https%3A%2F%2Fapi.box.com%2Fwopi%2Ffiles%2F1977927011490#RANGE!r6c6) | [lanthipeptide-class-i](https://docs.antismash.secondarymetabolites.org/glossary/#lanthipeptide-class-i) | 2,274,037 | 2,299,469 |  |  |
| [Region 6.7](https://word-edit.officeapps.live.com/we/wordeditorframe.aspx?ui=en-US&rs=en-US&IsLicensedUser=1&WOPISrc=https%3A%2F%2Fapi.box.com%2Fwopi%2Ffiles%2F1977927011490#RANGE!r6c7) | [terpene-precursor](https://docs.antismash.secondarymetabolites.org/glossary/#terpene-precursor) | 2,759,894 | 2,780,970 |  |  |
| [Region 6.8](https://word-edit.officeapps.live.com/we/wordeditorframe.aspx?ui=en-US&rs=en-US&IsLicensedUser=1&WOPISrc=https%3A%2F%2Fapi.box.com%2Fwopi%2Ffiles%2F1977927011490#RANGE!r6c8) | [oligosaccharide](https://docs.antismash.secondarymetabolites.org/glossary/#oligosaccharide) | 3,004,104 | 3,049,129 | Low | [calicheamicin](https://mibig.secondarymetabolites.org/go/BGC0000033.5) |
| [Region 6.9](https://word-edit.officeapps.live.com/we/wordeditorframe.aspx?ui=en-US&rs=en-US&IsLicensedUser=1&WOPISrc=https%3A%2F%2Fapi.box.com%2Fwopi%2Ffiles%2F1977927011490#RANGE!r6c9) | [T1PKS](https://docs.antismash.secondarymetabolites.org/glossary/#t1pks) | 3,147,902 | 3,193,727 |  |  |
| [Region 6.10](https://word-edit.officeapps.live.com/we/wordeditorframe.aspx?ui=en-US&rs=en-US&IsLicensedUser=1&WOPISrc=https%3A%2F%2Fapi.box.com%2Fwopi%2Ffiles%2F1977927011490#RANGE!r6c10) ^*^ | NRPS,RiPP-like | 3,297,609 | 3,344,838 | Low | [clipibicyclene/ azabicyclene B/ azabicyclene C/ azabicyclene D](https://mibig.secondarymetabolites.org/go/BGC0002697.3) |
| [Region 6.11](https://word-edit.officeapps.live.com/we/wordeditorframe.aspx?ui=en-US&rs=en-US&IsLicensedUser=1&WOPISrc=https%3A%2F%2Fapi.box.com%2Fwopi%2Ffiles%2F1977927011490#RANGE!r6c11) | [NI-siderophore](https://docs.antismash.secondarymetabolites.org/glossary/#ni-siderophore) | 3,438,715 | 3,470,370 | Low | [nonactin/ monactin/ dinactin/ trinactin/ tetranactin](https://mibig.secondarymetabolites.org/go/BGC0000244.4) |
| [Region 6.12](https://word-edit.officeapps.live.com/we/wordeditorframe.aspx?ui=en-US&rs=en-US&IsLicensedUser=1&WOPISrc=https%3A%2F%2Fapi.box.com%2Fwopi%2Ffiles%2F1977927011490#RANGE!r6c12) | [NRPS-like](https://docs.antismash.secondarymetabolites.org/glossary/#nrps-like) | 3,546,654 | 3,590,517 |  |  |

*Three hybrid biosynthetic gene clusters are predicted, but due to the distance between the core genes, these are considered as two independent gene clusters in our total count.

**Supplementary Result 4:** Antibacterial activity of *N. bonnevillensis* crude extract and fractions. A crude extract of a *N. bonnevillensis* fermentation was screened for antimicrobial activity (Supplemental Table 3), which showed moderate to strong activity against several bacterial pathogens. This screening was part of a broader effort to evaluate extremophilic *Actinomycetota* from Great Salt Lake as a resource of novel antibiotics. Bioactivity was also observed in the fractionated crude extract (Supplemental Table 4), and these fractions were further evaluated to identify individual constituents responsible for the observed bioactivity. The observed antimicrobial activity is consistent with the genomic prediction of substantial biosynthetic potential (Extended Fig. 2a and Supplementary Table 2), suggesting *N. bonnevillensis* encodes pathways capable of producing previously uncharacterized natural products.

**Supplementary Table 3**: Antimicrobial activity of crude extract from *N. bonnevillensis* cultured in A1 media. Activity and zone of inhibition diameters are reported as: NA, no activity; P, ≤ 10 mm (weak activity); PP, 10–15 mm (moderate activity); PPP, > 15 mm (strong activity). Fractions screened at 10 mg/mL and 1 mg/mL, 20 μL was added to each disc. Ciprofloxacin: 5 μg was included as a positive control.

|  | MRSA TCH1516 | *E. coli* LptD4213 | *P. aeruginosa* PA01 | *B. oceanisediminis* CNY997 | *S. marcescens* SNA111 | *V. haveryi* SNA117 |
| --- | --- | --- | --- | --- | --- | --- |
| Total Crude Extract | PP | PP | NA | PPP | NA | NA |
| Ciprofloxacin | PPP | PPP | PPP | PPP | PPP | PPP |

**Supplementary Table 4**: Antimicrobial activity of fractionated extracts from *N. bonnevillensis* cultured in A1 media. Activity and zone of inhibition diameters are reported as: NA, no activity; P, ≤ 10 mm (weak activity); PP, 10–15 mm (moderate activity); PPP, > 15 mm (strong activity). Crude extract concentration: 20 mg/mL, 20 μL was added to each disc. Ciprofloxacin: 5 μg was included as a positive control.

|  | MRSA TCH1516 | | *E. coli* LptD4213 | |
| --- | --- | --- | --- | --- |
|  | 10 mg/mL | 1 mg/mL | 10 mg/mL | 1 mg/mL |
| Fraction 1 | NA | NA | NA | NA |
| Fraction 2 | NA | NA | NA | NA |
| Fraction 3 | PPP | NA | PPP | PP |
| Fraction 4 | PPP | NA | PP | PP |
| Fraction 5 | P | NA | P | NA |
| Fraction 6 | PPP | NA | PPP | PPP |
| Ciprofloxacin | - | PPP | - | PPP |

**Supplementary Result 5:** Isolation of bonnevanoside

Guided by the bioactivity results (Supplemental Tables 3 and 4), fraction 4 was selected for further chemical characterization. Bioassay-guided purification of this fraction led to the isolation of bonnevanoside (**1**). High-resolution electrospray ionization mass spectrometry established the molecular formula as C₂₆H₃₃NO₁₂S (Supplementary Fig. 4), and 1D & 2D NMR analyses confirmed the structure as a thiophenyl nonulopyranoside (Supplemental Fig. 5 and Supplemental Table 5). Spectroscopic analysis of 1D and 2D NMR data (Supplementary Fig. 6–13), combined with comparison to the synthetic standard thiophenyl(methyl-5-acetamido-4,7,8,9-tetra-O-acetyl-3,5-dideoxy-β-d-glycero-d-galacto-2-nonulopyranoside), confirmed that the two molecules are structurally identical^7,8^. The syn-orientation of H-3a, H-4, and H-6 was further supported by NOE correlations between H-3a/H-4 and H-4/H-6. Likewise, NOE correlations between H-3b/H-5 and H-5/H-7 supported a syn-orientation of H-3b, H-5, and H-7, consistent with a large coupling constant (J₄,₅ = 10.4 Hz) between H-4 and H-5. An NOE correlation between H-6 and H-3′, together with the downfield shift of H-4 (δH 5.35) and a small J₇,₈ (2.6 Hz), established a β-anomeric configuration, which was further validated by molecular modeling. These findings, along with comparison to the synthetic standard, confirmed the structural assignment. Although **1** has been previously synthesized, this study represents the first report of its isolation from a natural source. Bonnevanoside did not display significant antibacterial activity, showing no inhibition against methicillin-resistant *S. aureus* or *E. coli* in agar disk diffusion assays at 100 μg/mL (Supplementary Table 6).

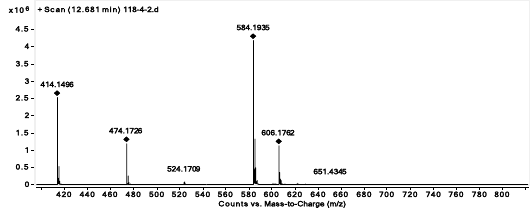

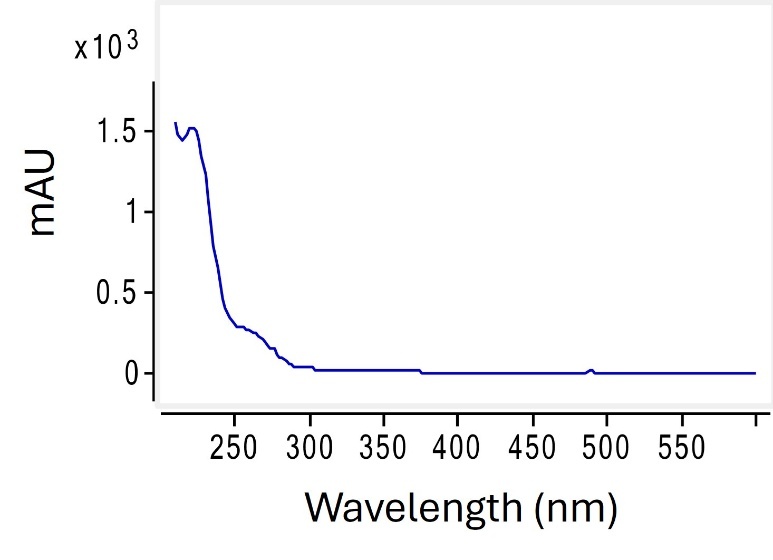

**Supplementary Figure 4**. Negative HR-ESI-TOF-MS and UV spectra of bonnevanoside (**1**).

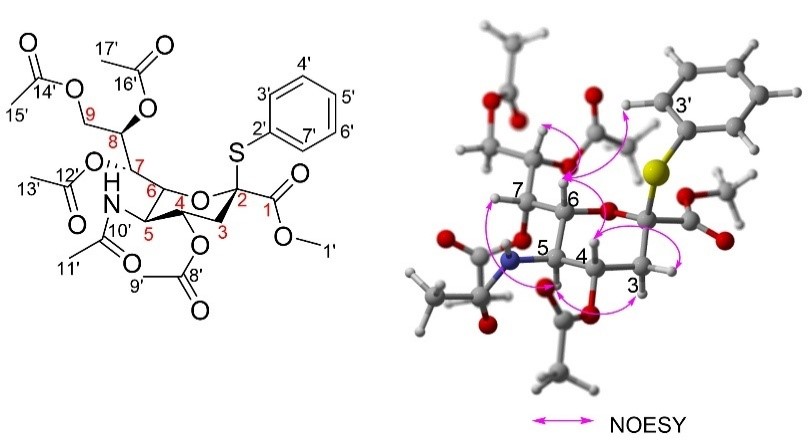

**Supplementary Figure 5**. Structure and key 2D NMR NOESY correlations of bonnevanoside (**1**).

**Supplementary Table 5**. NMR spectral data of **1** in MeOD

| Position | δC*^a^* | Type | δH*^b^*, mult (*J* in Hz) | HMBC | COSY | NOESY |
| --- | --- | --- | --- | --- | --- | --- |
| 1 | 168.8 | C |  |  |  |  |
| 2 | 88.7 | C |  |  |  |  |
| 3 | 37.0 | CH_2_ | a 2.63, dd (13.9, 4.9) | C-2, 4, 5 |  | H-4, 6 |
|  |  |  | b 2.04, d (3.3) |  | H-4 |  |
| 4 | 69.2 | CH | 5.32, ddd (11.6, 10.4, 4.8) | C-3, 5, 8' |  | H-6 |
| 5 | 48.9 | CH | 3.97, m | C-3, 4, 6, 10' | H-4, 6 | H-7, 3b |
| 6 | 72.4 | CH | 4.70, dd (4.6, 2.5) | C-2, 4, 5, 8 |  | H-7, 8, 9a |
| 7 | 68.9 | CH | 5.43, t (2.6) | C-6, 9, 12' | H-6, 8 | H-8 |
| 8 | 72.7 | CH | 4.94, dt (8.4, 2.5) | C-6, 9, 14' | H-9a | H-5 |
| 9 | 62.5 | CH_2_ | a 4.40, dd (12.2, 2.3) | C-7, 8, 16' |  |  |
|  |  |  | b 3.93, m |  |  |  |
| 1' | 51.8 | CH_3_ | 3.59, s | C-1 |  |  |
| 2' | 128.9 | C |  |  | H-3' |  |
| 3' | 136.0 | CH | 7.46, dd (8.5, 1.5) | C-5', 7' | H-4' | H-6 |
| 4' | 129.1 | CH | 7.35, t (8.5) | C-6' | H-5' |  |
| 5' | 129.7 | CH | 7.40, t (8.5) | C-3', 7' | H-6' |  |
| 6' | 129.1 | CH | 7.35, t (8.5) | C-4' |  |  |
| 7' | 136.0 | CH | 7.46, dd (8.5, 1.5) | C-3', 5' |  |  |
| 8' | 170.5 | C |  |  |  |  |
| 9' | 19.6 | CH_3_ | 1.97, s | C-8' |  |  |
| 10' | 172.2 | C |  |  |  |  |
| 11' | 21.3 | CH_3_ | 1.84, s | C-10' |  |  |
| 12' | 171.0 | C |  |  |  |  |
| 13' | 19.4 | CH_3_ | 1.99, d (6.3) | C-12' |  |  |
| 14' | 170.3 | C |  |  |  |  |
| 15' | 19.6 | CH_3_ | 2.04, s | C-14' |  |  |
| 16' | 170.8 | C |  |  |  |  |
| 17' | 19.6 | CH_3_ | 1.93, s | C-16' |  |  |

*^a^* 125 MHz, *^b^* 500 MHz

**Supplementary Table 6**. Antibacterial activity of bonnevanoside (**1**). Agar disk diffusion assays were performed in triplicate. Ciprofloxacin (1 µg/mL) was used as a positive control and consistently fell within the expected inhibition range, confirming assay validity. Test concentrations of **1** were performed at 0.01–0.1 mg/mL. Inhibitions zones were categorized as follows: P, ≤ 10 mm (weak activity); P.P., 10–15 mm (moderate activity); P.P.P., > 15 mm (strong activity).

|  | Compound **1**  100 μg/mL | Compound **1**  10 μg/mL | Ciprofloxacin  10 μg/mL |
| --- | --- | --- | --- |
| Methicillin-resistant *Staphyloccus aureus* TCH1516 | NA | NA | P.P.P |
| *Escherichia coli*  LptD4213 | NA | NA | P.P.P |

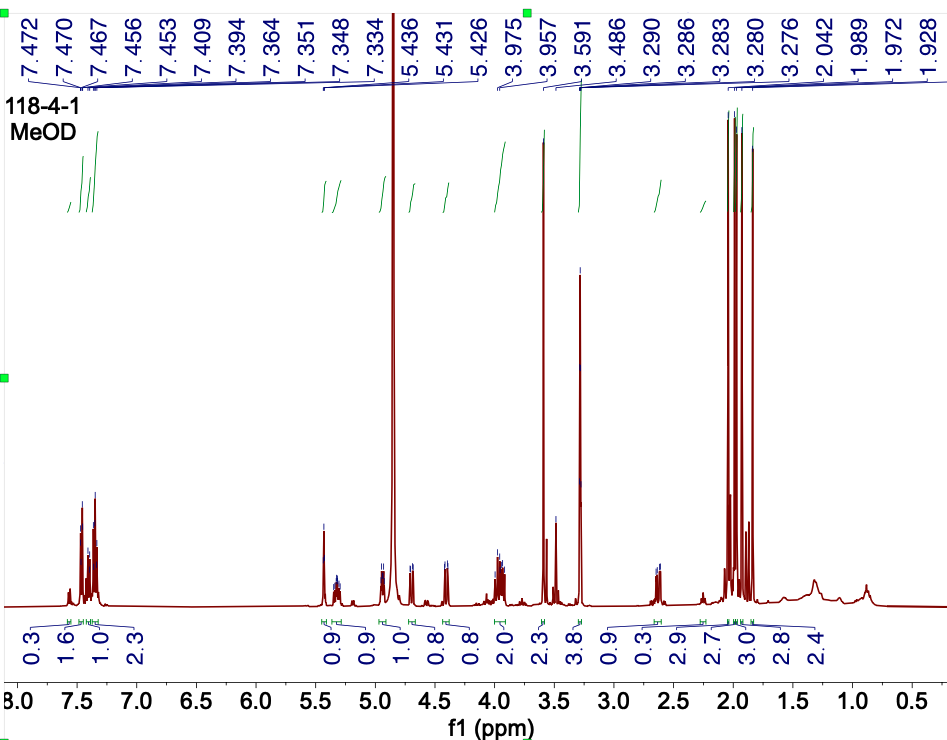

**Supplementary Figure 6**. ^1^H NMR spectrum of **1** at 500 MHz in MeOD.

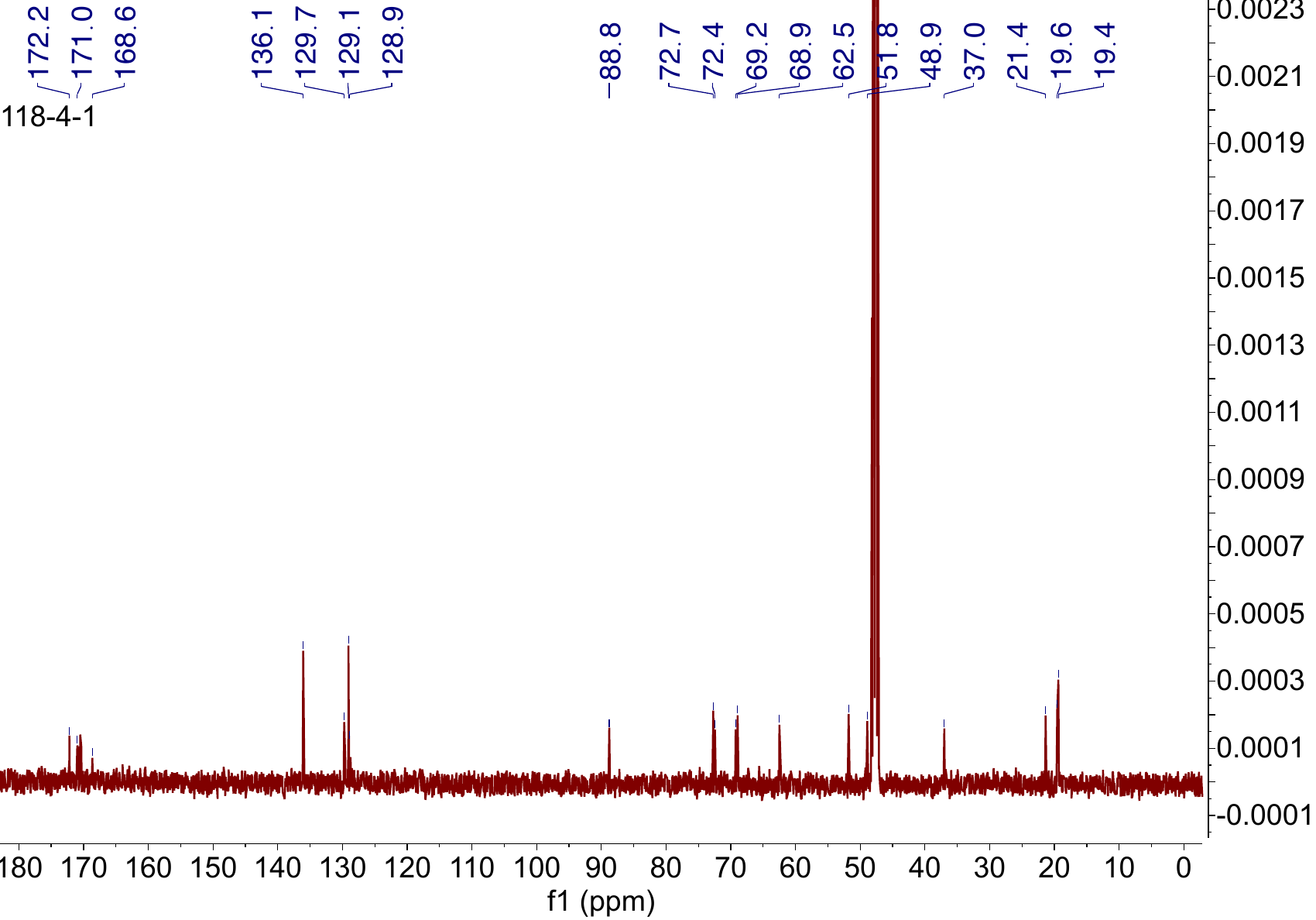

**Supplementary Figure 7**. ^13^C NMR spectrum of **1** at 125 MHz in MeOD.

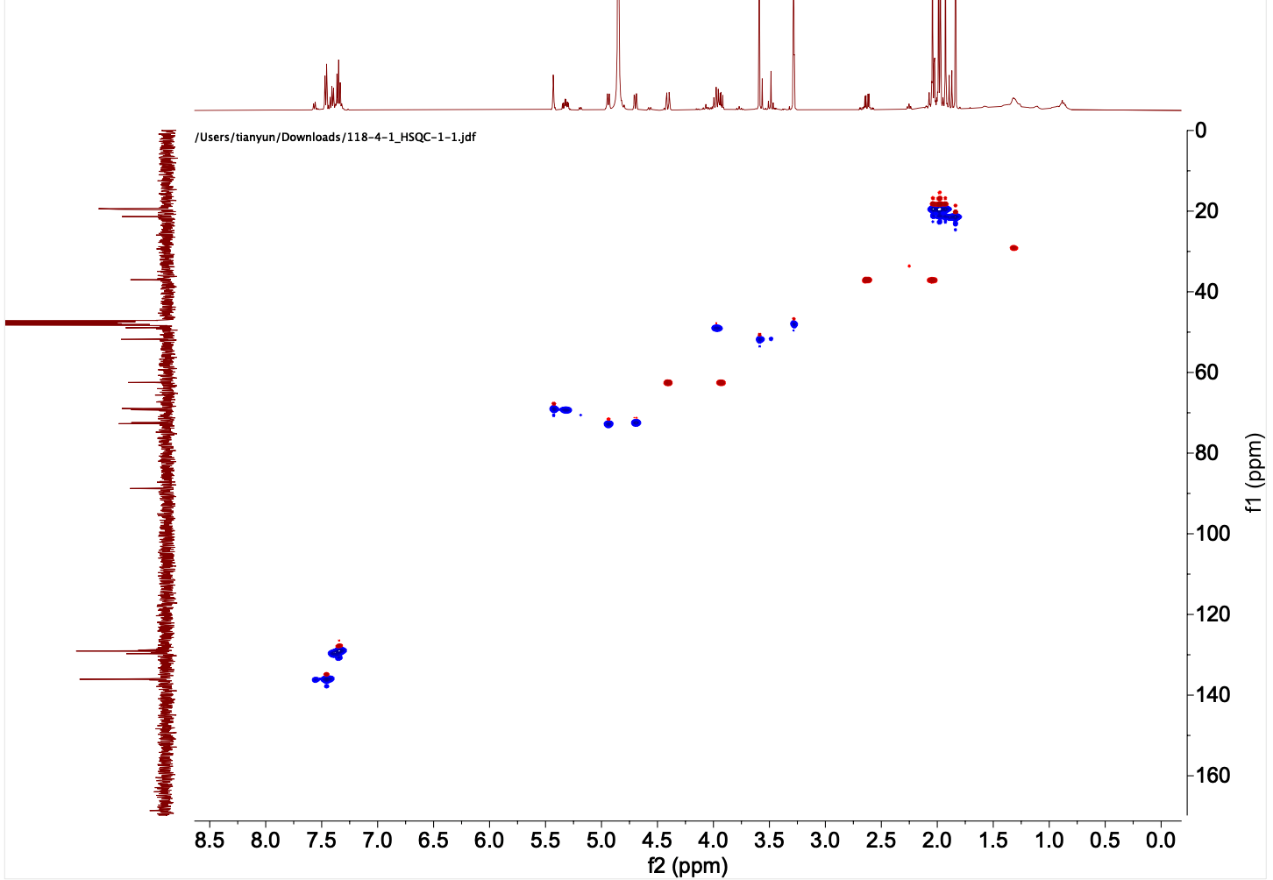

**Supplementary Figure 8**. HSQC NMR spectrum of **1** at 500 MHz in MeOD.

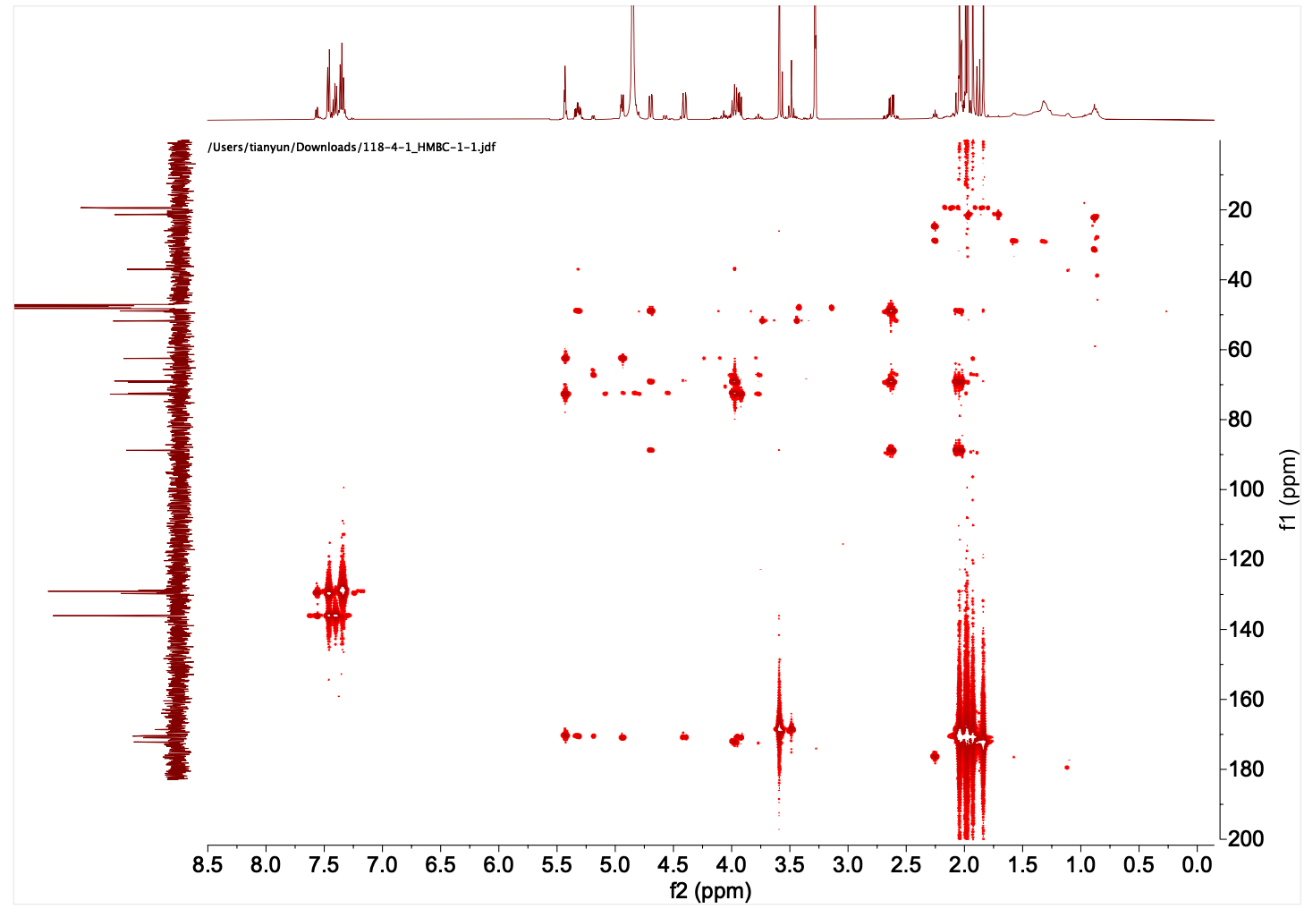

**Supplementary Figure 9**. HMBC NMR spectrum of **1** at 500 MHz in MeOD.

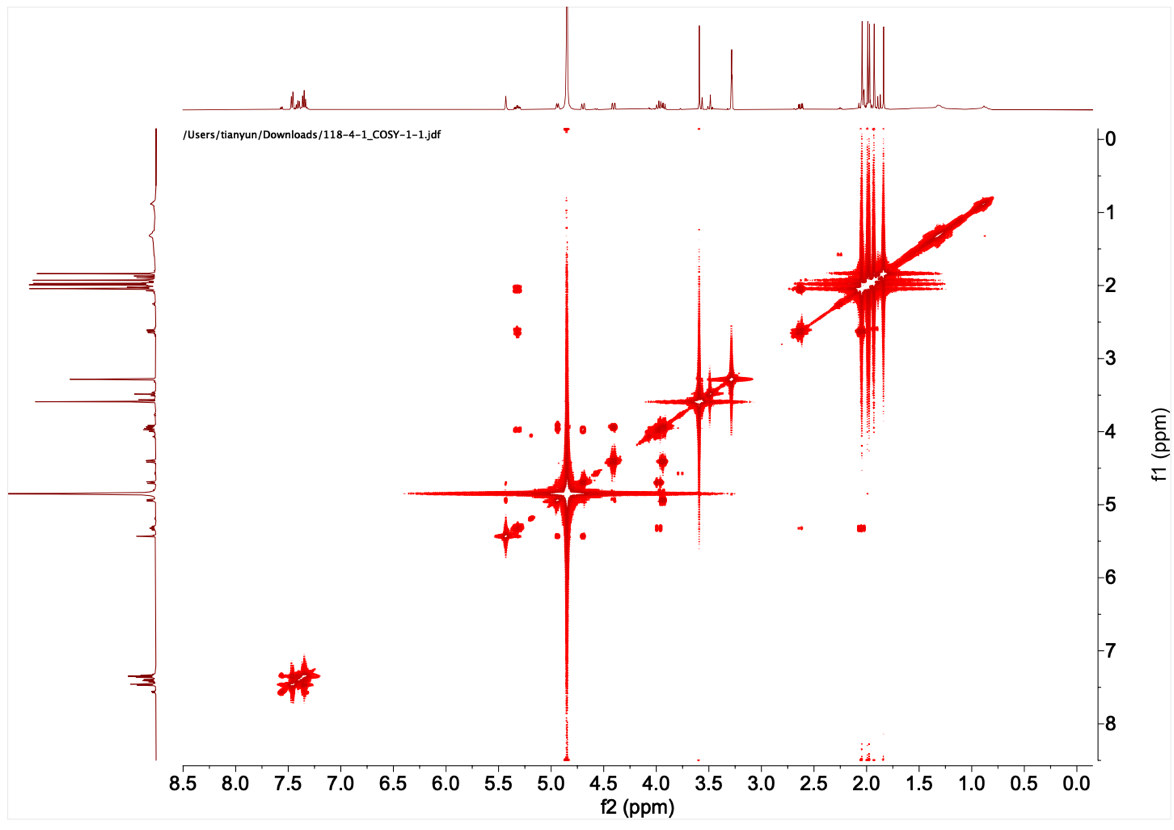

**Supplementary Figure 10**. COSY NMR spectrum of **1** at 500 MHz in MeOD.

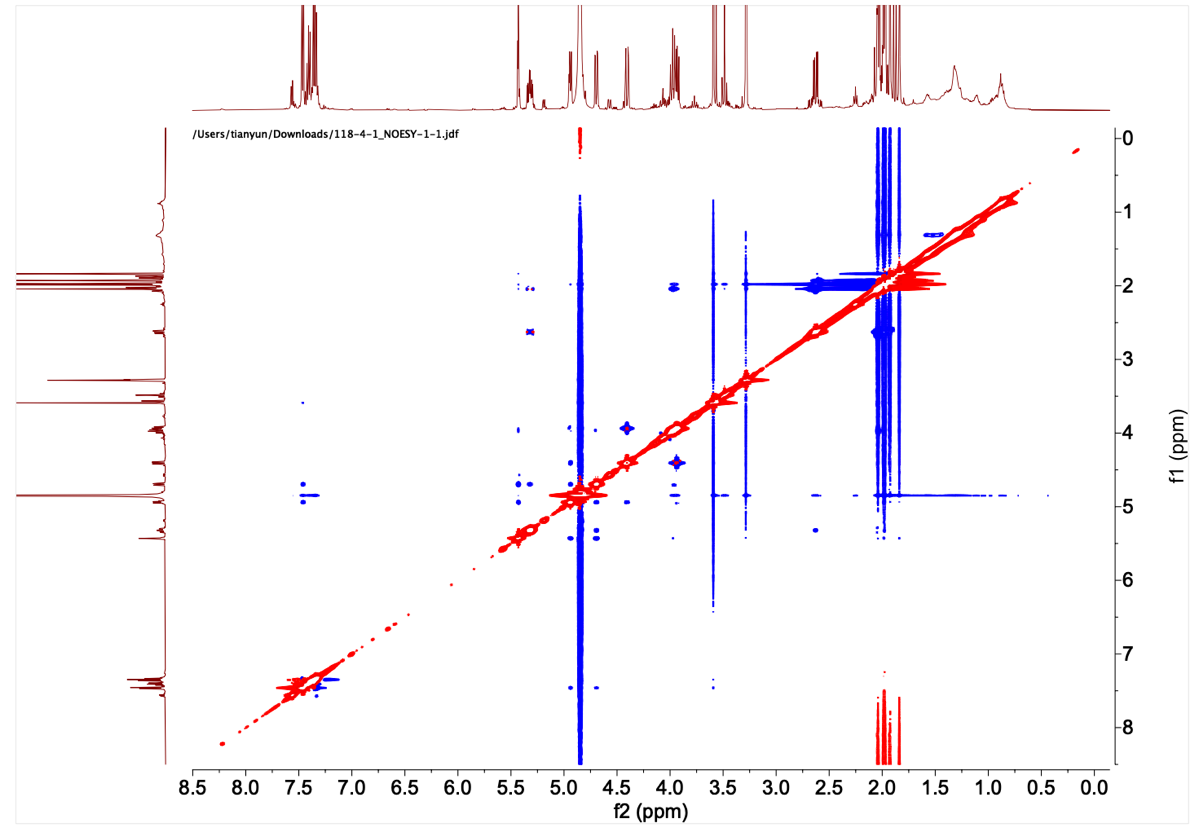

**Supplementary Figure 11**. NOESY NMR spectrum of **1** at 500 MHz in MeOD.

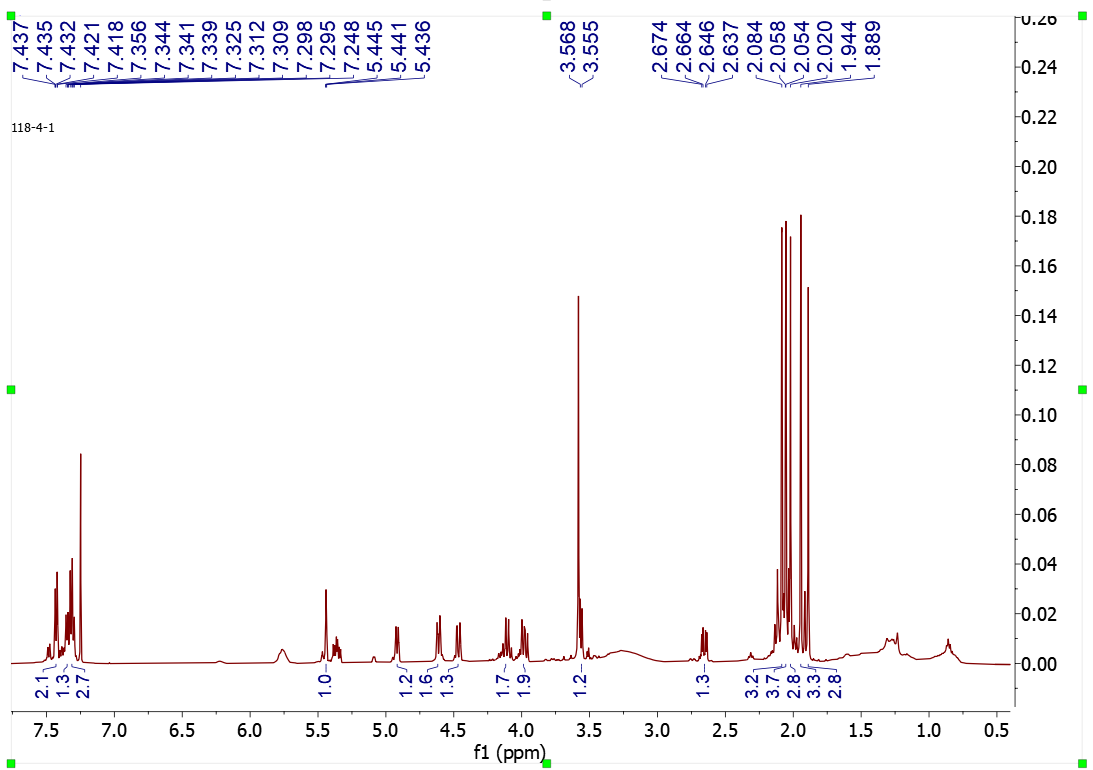

**Supplementary Figure 12**. ^1^H NMR NMR spectrum of **1** at 500 MHz in CDCl_3_.

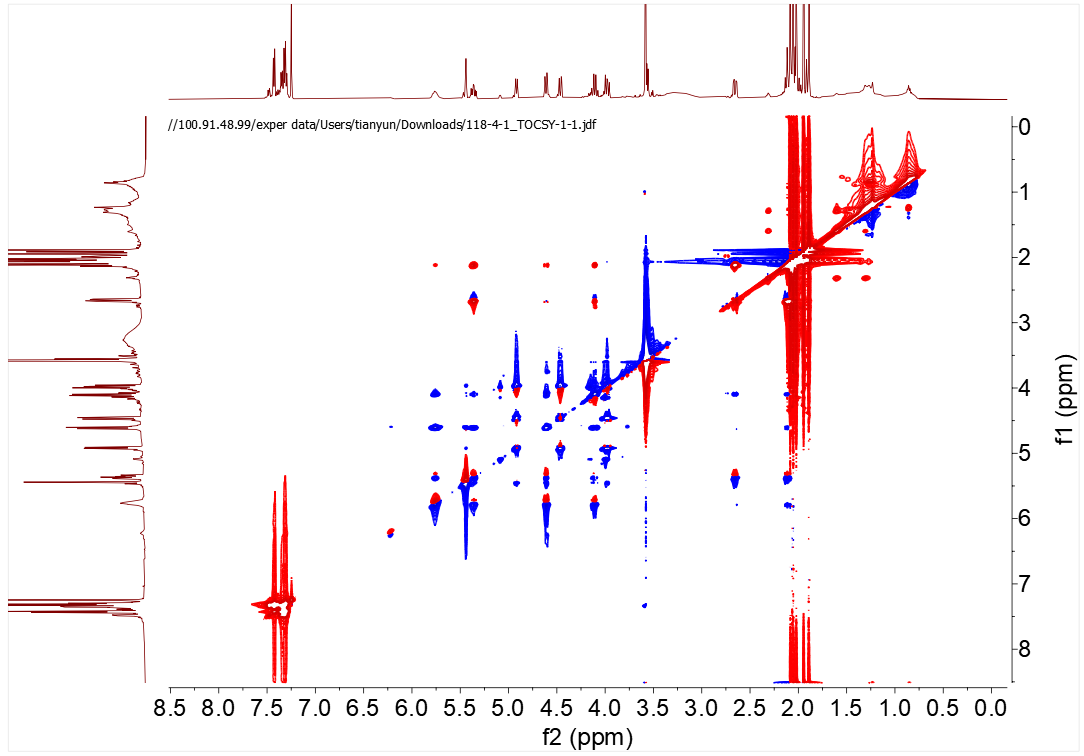

**Supplementary Figure 13**. TOCSY NMR spectrum of **1** at 500 MHz in CDCl_3_.
**Supplementary Table 7**: List of known mycolate-producing species used to compile protein sequences for mycolic acid biosynthesis gene homolog identification in *N. bonnevillensis* genome.

| **Organism** | **GenBank Accession Number** |
| --- | --- |
| *Mycobacterium tuberculosis* H37Rv | GCA_000195955.2 |
| *Mycobacterium smegmatis* MC2 155 | GCA_000767605.1 |
| *Mycobacterium koreense* JCM 19956 | GCA_010731835.1 |
| *Mycobacterium abscessus* ATCC 19977 | GCA_000069185.1 |
| *Mycobacterium sinense* JDM601 | GCA_000214155.1 |
| *Corynebacterium glutamicum* ATCC 13032 | GCA_000011325.1 |
| *Nocardia farcinica* DSM 43257 | GCA_000009805.1 |
| *Rhodococcus jostii* RHA1 | GCA_000014565.1 |
| *Rhodococcus equi* 103S | GCA_000196695.1 |
| *Williamsia herbipolensis* NBC 00319 | GCA_036153515.1 |
| *Gordonia bronchialis* DSM 43247 | GCA_000024785.1 |
| *Skermania pinensis* DSM 43998 | GCA_019285775.1 |
| *Tsukamurella paurometabola* DSM 20162 | GCA_000092225.1 |
| *Dietzia timorensis* ID05-A0528 | GCA_001659785.1 |
| *Tomitella fengzijianii* HY188 | GCA_007559025.1 |

**Supplementary Table 8**: Reference genomes used to create Fig. 1b.

| Strain | NCBI Accession Number | 16S Sequence Length (bp) |
| --- | --- | --- |
| *Streptomyces avermitilis* MA-4680 = NBRC 14893 | GCA_000009765.2 | 1448 |
| *Corynebacterium glutamicum* ATCC 13032 | GCA_000011325.1 | 1446 |
| *Gordonia bronchialis* DSM 43247 | GCA_000024785.1 | 1445 |
| *Tsukamurella paurometabola* DSM 20162 | GCA_000092225.1 | 1441 |
| *Mycobacterium kansasii* ATCC 12478 | GCA_000157895.2 | 1453 |
| *Corynebacterium resistens* DSM 45100 | GCA_000177535.2 | 1443 |
| *Mycobacterium tuberculosis* H37Rv | GCA_000195955.2 | 1454 |
| *Streptomyces xinghaiensis* S187 | GCA_000220705.2 | 1448 |
| *Dietzia alimentaria* 72 | GCA_000226215.2 | 1442 |
| *Mycobacterium paraintracellulare* | GCA_000276825.1 | 1453 |
| *Nocardia pneumoniae* NBRC 100136 | GCA_000308755.1 | 1441 |
| *Nocardia veterana* NBRC 100344 | GCA_000308855.1 | 1441 |
| *Streptomyces rimosus* subsp. rimosus ATCC 10970 | GCA_000331185.2 | 1449 |
| *Mycobacterium haemophilum* DSM 44634 | GCA_000340435.3 | 1454 |
| *Nocardiopsis valliformis* DSM 45023 | GCA_000340985.1 | 1455 |
| *Nocardiopsis alkaliphila* YIM 80379 | GCA_000341005.1 | 1233 |
| *Nocardiopsis ganjiahuensis* DSM 45031 | GCA_000341085.1 | 1455 |
| *Williamsia herbipolensis* | GCA_000964005.1 | 1438 |
| *Mycobacterium lepromatosis* | GCA_000975265.2 | 1470 |
| *Gordonia phthalatica* | GCA_001305675.1 | 1443 |
| *Corynebacterium diphtheriae* | GCA_001457455.1 | 1441 |
| *Mycolicibacterium smegmatis* | GCA_001457595.1 | 1444 |
| *Nocardiopsis listeri* NBRC 13360 | GCA_001570765.1 | 1457 |
| *Dietzia papillomatosis* NBRC 105045 | GCA_001570845.1 | 1440 |
| *Tsukamurella pseudospumae* | GCA_001575195.1 | 1133 |
| *Mycobacterium adipatum* | GCA_001644575.1 | 1440 |
| *Dietzia timorensis* | GCA_001659785.1 | 1438 |
| *Streptomyces lincolnensis* | GCA_001685355.1 | 1450 |
| *Gordonia terrae* | GCA_001698225.1 | 1445 |
| *Nocardia mangyaensis* | GCA_001886715.1 | 1439 |
| *Rhodococcus maanshanensis* NBRC 100610 | GCA_001894865.1 | 1438 |
| *Rhodococcus zopfii* | GCA_001895025.1 | 1445 |
| *Corynebacterium pseudotuberculosis* | GCA_002155265.1 | 1442 |
| *Dietzia lutea* | GCA_003096075.1 | 1442 |
| *Dietzia psychralcaliphila* | GCA_003096095.1 | 1439 |
| *Rhodococcus oxybenzonivorans* | GCA_003130705.1 | 1438 |
| *Williamsia limnetica* | GCA_003217475.1 | 1443 |
| *Dietzia kunjamensis* | GCA_003610395.1 | 1439 |
| *Nocardia yunnanensis* | GCA_003626895.1 | 1441 |
| *Williamsia muralis* | GCA_003634525.1 | 1443 |
| *Gordonia insulae* | GCA_003855095.1 | 1443 |
| *Nocardia bhagyanarayanae* | GCA_006716565.1 | 1441 |
| *Gordonia zhaorongruii* | GCA_007559005.1 | 1443 |
| *Tsukamurella asaccharolytica* | GCA_007858435.1 | 1441 |
| *Tsukamurella sputi* | GCA_007858445.1 | 1390 |
| *Tsukamurella conjunctivitidis* | GCA_007858475.1 | 1311 |
| *Rhodococcus rhodnii* | GCA_008011915.1 | 1444 |
| *Streptomyces venezuelae* ATCC 10712 | GCA_008639165.1 | 1448 |
| *Streptomyces fradiae* ATCC 10745 = DSM 40063 | GCA_008704425.1 | 1450 |
| *Mycobacterium avium* subsp. avium | GCA_009741445.1 | 1455 |
| *Mycobacterium conspicuum* | GCA_010730195.1 | 1453 |
| *Tsukamurella spumae* | GCA_012396015.1 | 1441 |
| *Nocardia huaxiensis* | GCA_013744875.1 | 1439 |
| *Gordonia jinghuaiqii* | GCA_014041935.1 | 1445 |
| *Nocardiopsis metallicus* | GCA_014201115.1 | 1455 |
| *Williamsia phyllosphaerae* | GCA_014635305.1 | 1425 |
| *Nocardiopsis terrae* DSM 45157 | GCA_014874035.1 | 1455 |
| *Corynebacterium amycolatum* | GCA_016889425.1 | 1442 |
| *Rhodococcus pseudokoreensis* | GCA_017068395.1 | 1441 |
| *Williamsia soli* | GCA_017377795.1 | 1444 |
| *Nocardiopsis eucommiae* | GCA_018388625.1 | 1455 |
| *Dietzia massiliensis* | GCA_018390595.1 | 1442 |
| *Nocardia iowensis* | GCA_019222765.1 | 1441 |
| *Mycobacterium malmoense* | GCA_019645855.1 | 1453 |
| *Streptomyces mobaraensis* | GCA_020099395.1 | 1448 |
| *Nocardia pulmonis* | GCA_023740755.1 | 1441 |
| *Nocardiopsis exhalans* | GCA_024134545.1 | 1455 |
| *Williamsia deligens* | GCA_024171765.1 | 1434 |
| *Williamsia maris* | GCA_024171815.1 | 1439 |
| *Gordonia mangrovi* | GCA_024734075.1 | 1445 |
| *Streptomyces murinus* | GCA_025231465.1 | 1447 |
| *Gordonia pseudamarae* | GCA_025273675.1 | 1444 |
| *Rhodococcus sacchari* | GCA_025837095.1 | 1445 |
| *Nocardia sputorum* | GCA_027924405.1 | 1441 |
| *Corynebacterium jeikeium* | GCA_028609885.1 | 1450 |
| *Corynebacterium durum* | GCA_030408675.1 | 1465 |
| *Streptomyces collinus* | GCA_031348265.1 | 1448 |
| *Gordonia hydrophobica* | GCA_038024885.1 | 1441 |
| *Dietzia natronolimnaea* | GCA_039525805.1 | 1439 |
| *Williamsia serinedens* | GCA_039530165.1 | 1432 |
| *Tsukamurella soli* | GCA_039541155.1 | 1440 |
| *Dietzia aerolata* | GCA_042685785.1 | 1439 |
| *Gordonia amarae* | GCA_044358485.1 | 1443 |
| *Rhodococcus parequi* | GCA_046042785.1 | 1441 |
| *Rhodococcus tukisamuensis* | GCA_900101735.1 | 1439 |
| *Corynebacterium mycetoides* | GCA_900103625.1 | 1443 |
| *Rhodococcus jostii* | GCA_900105375.1 | 1440 |
| *Williamsia sterculiae* | GCA_900156495.1 | 1442 |
| *Corynebacterium urealyticum* | GCA_900187235.1 | 1443 |
| *Corynebacterium cystitidis* | GCA_900187295.1 | 1444 |
| *Tsukamurella pulmonis* | GCA_900460155.1 | 1443 |
| *Rhodococcus coprophilus* | GCA_900478115.1 | 1445 |
| *Nocardia asteroides* | GCA_900637185.1 | 1439 |
| *Tsukamurella tyrosinosolvens* | GCA_900637875.1 | 1443 |
| *Streptomyces clavuligerus* | GCA_005519465.1 | 1450 |
| *Tsukamurella strandjordii* | NR_025113.1 | 1512 |
| *Dietzia cercidiphylli* | NR_044483.1 | 1456 |
| *Nocardiopsis prasina* DSM 43845 | NR_044906.1 | 1461 |
| *Nocardiopsis bonnevillensis* | NA | 1455 |

**Supplementary Table 9**: 16S rRNA primers

| Primer Name | Sequence (5'→ 3') |
| --- | --- |
| 27F | AGAGTTTGATCMTGGCTCAG |
| 1492R | CGGTTACCTTGTTACGACTT |

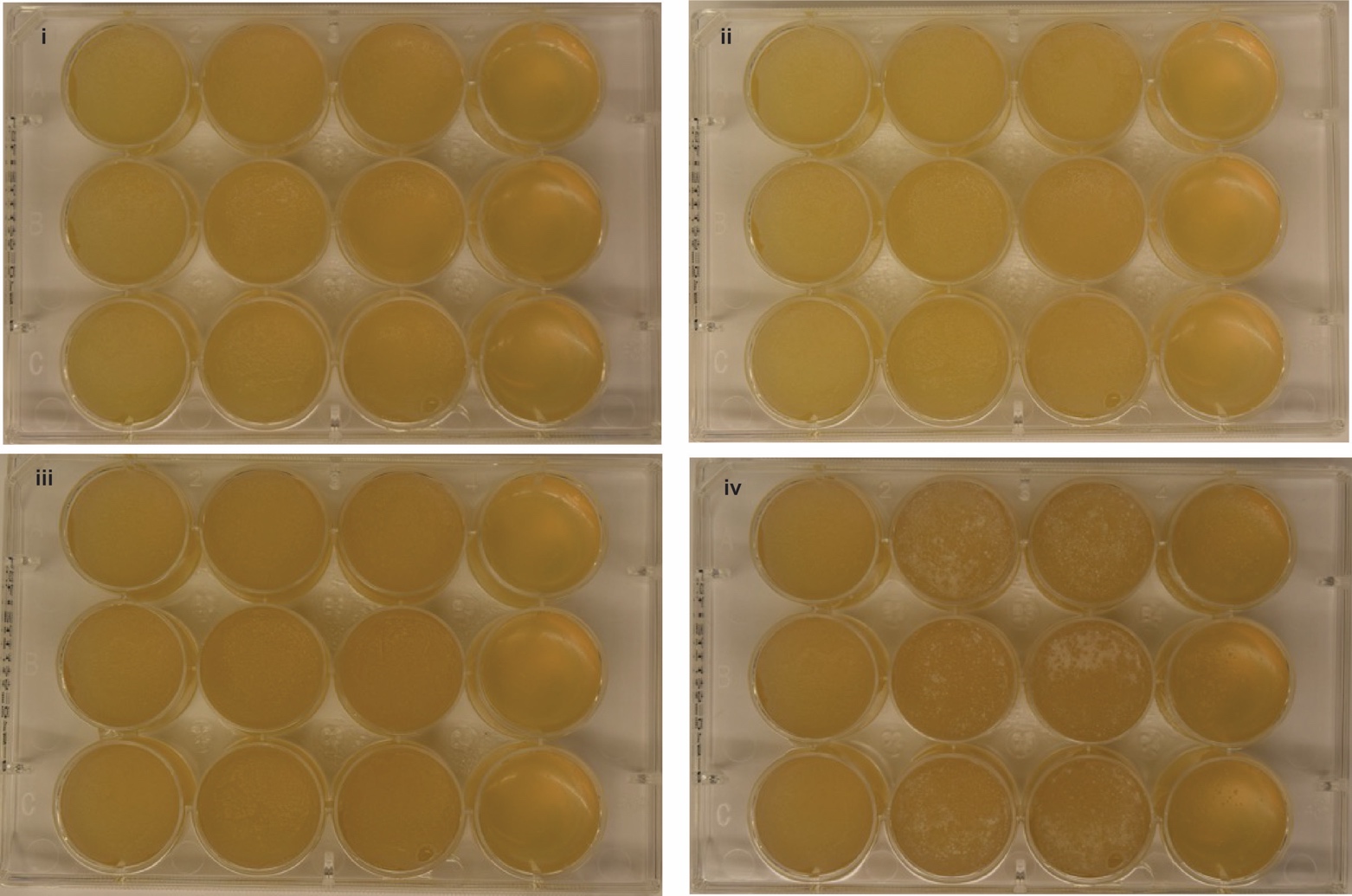

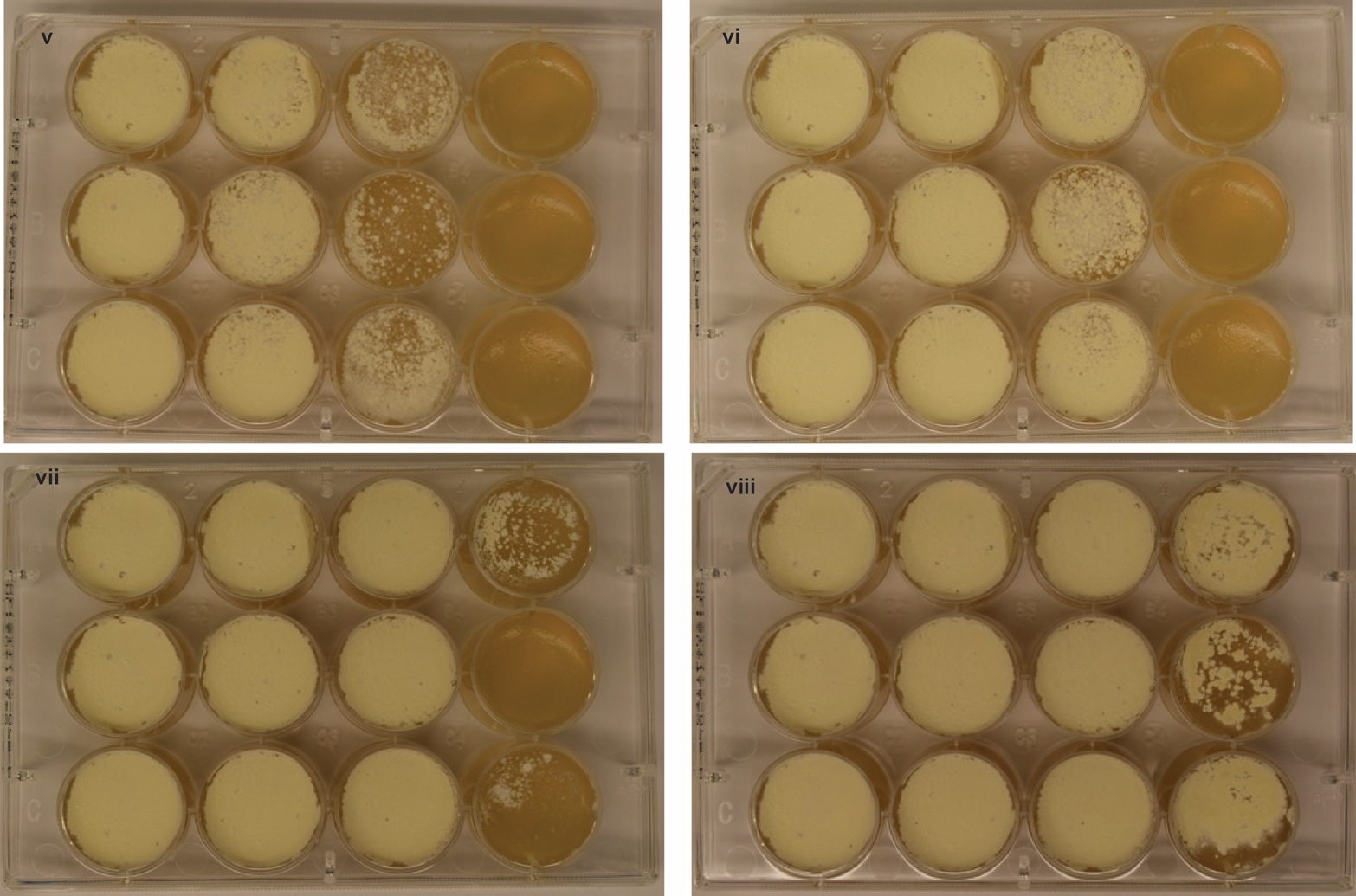

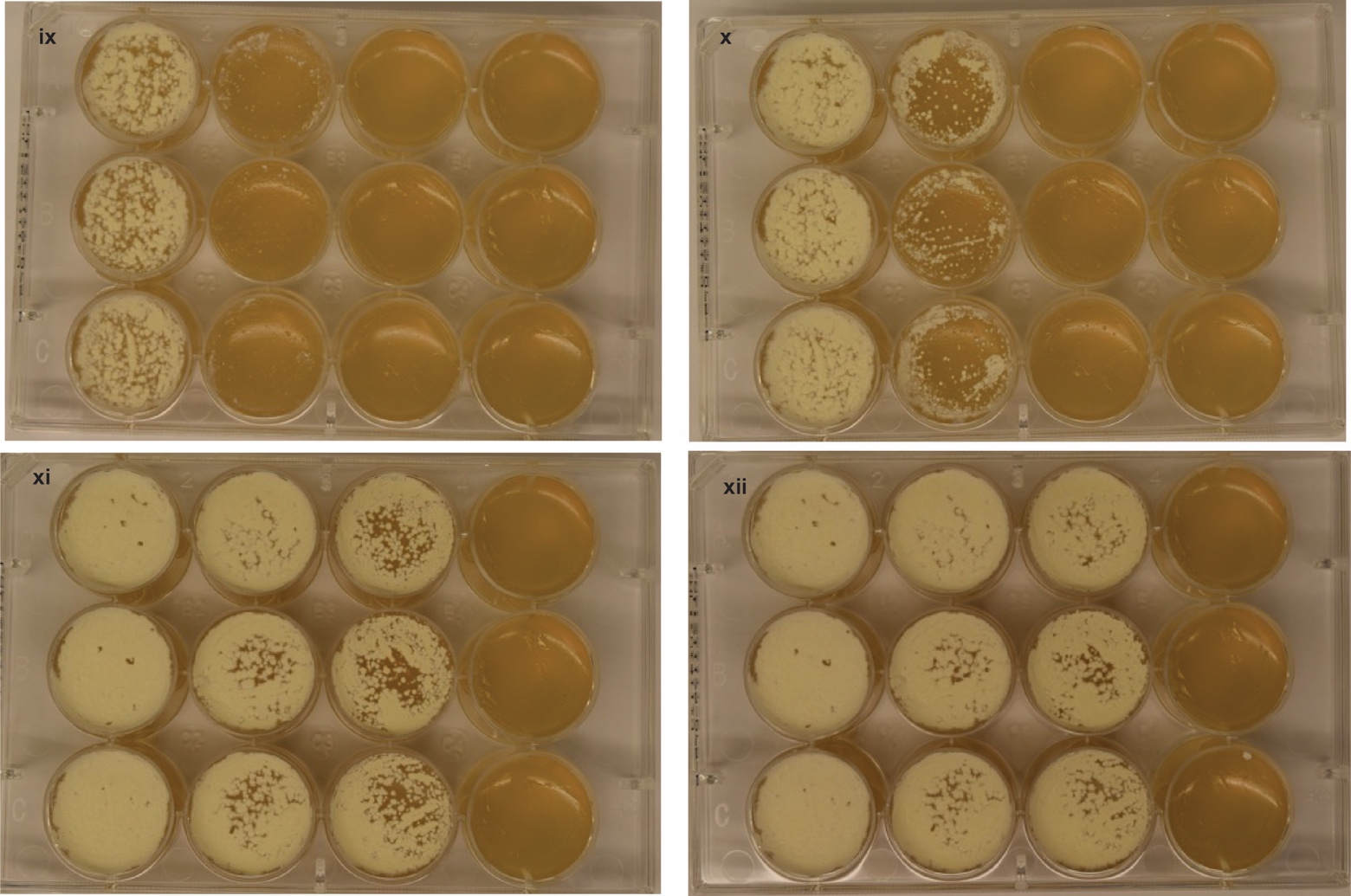

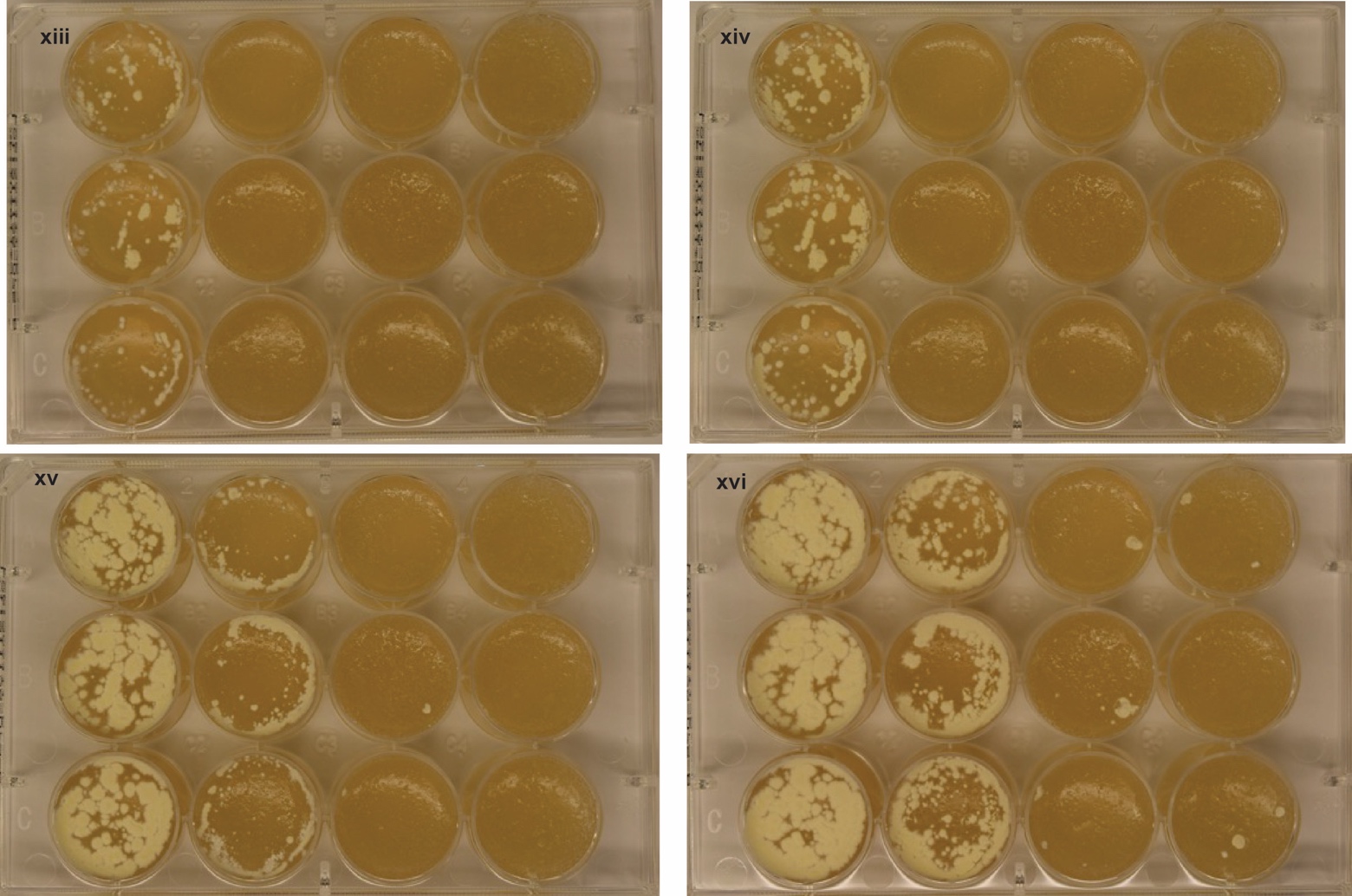

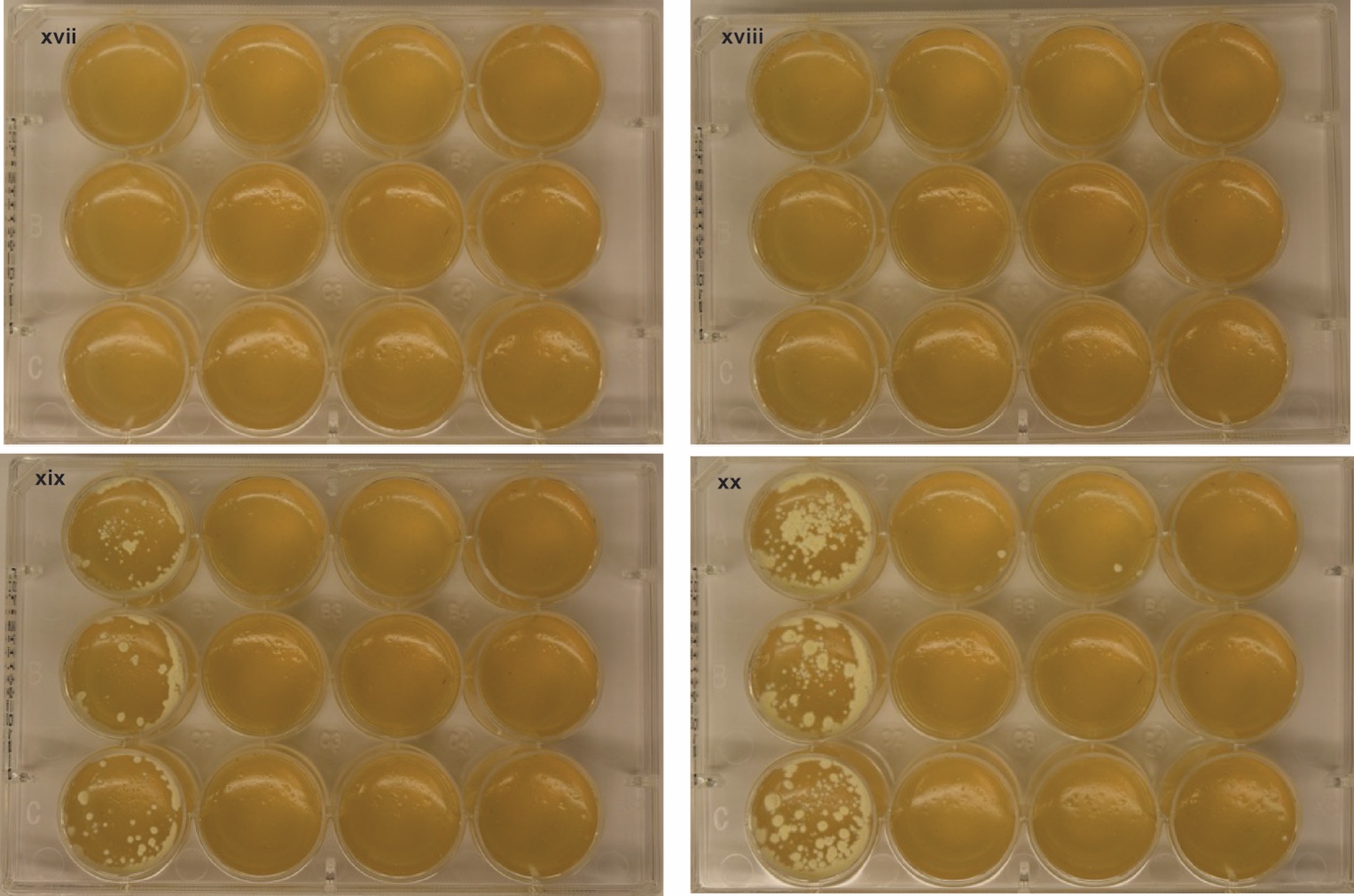

**Supplementary Figure 14**: Growth of *N. bonnevillensis* across a range of isoniazid and salinity concentrations. Growth of *N. bonnevillensis* was monitored on YPM agar plates supplemented with increasing concentrations of isoniazid (INH) (columns 1–4 for all plates at: 0, 12.5, 25, and 200 µg mL⁻¹, respectively). Rows are replicates for each condition being tested. Within each salinity condition, replicate plates were imaged at four time points: Day 5, Day 6, Day 10, and Day 14.. i-iv (YPM + 0% salinity): i, Day 5; ii, Day 6; iii, Day 10; and iv, Day 14. v-viii (YPM + 4% salinity): v, Day 5; vi, Day 6; vii, Day 10; and viii, Day 14. ix-xii (YPM + 8% salinity): ix, Day 5; x, Day 6; xi, Day 10; and xii, Day 14. xiii-xvi (YPM + 12% salinity): xiii, Day 5; xiv, Day 6; xv, Day 10; and xvi, and Day 14. xvii-xx (YPM + 16% salinity): xvii, Day 5; xviii, Day 6; xix, Day 10; and xx, Day 14.
